# Suprathreshold suppression underlies spatial selectivity in temporal interference stimulation

**DOI:** 10.64898/2026.11.22.768107

**Authors:** Jeongju Moon, Ji-Hyun Jeong, Younghoon Park, Seung Hyun Min, Ha Rim Oh, Hong Ju Lee, Jeongho Choi, Sung-Yun Park, Byung Tae Choi, Chin Su Koh, Hyun Ho Jung, Seungsoo Chung, Kyungsik Eom

## Abstract

Temporal interference stimulation (TIS) has been proposed for non-invasive deep-brain targeting using temporally interfering kilohertz-frequency electric fields, but the neuronal basis of its spatial selectivity remains unclear. Here we show that differential suppression of suprathreshold firing can support spatially selective recruitment. Multiscale simulations revealed no activation-threshold advantage for TIS over unmodulated high-frequency stimulation (HFS). At higher currents, HFS produced sustained depolarization and firing suppression, whereas strongly modulated TIS permitted periodic repolarization and maintained beat-locked firing. Spatially varying responses supported selective firing and current-ratio-dependent steering over an intensity-dependent range. Whole-cell recordings in mouse cortical slices under pharmacological synaptic blockade supported the predicted membrane-potential and firing dynamics. Retinal multielectrode-array recordings showed spatially patterned preservation of firing and current-ratio-dependent steering. In vivo intracortical stimulation in rats produced stronger beat-frequency locking of motor responses with a premodulated TIS waveform than with HFS. Our findings link suprathreshold neuronal response dynamics to spatial selectivity and characterize an intensity-dependent operating regime for TIS.

## Introduction

Non-invasive electrical neuromodulation provides a means to probe and modulate neural circuits without surgically implanted electrodes^1, 2, 3^. Transcranial electrical stimulation (tES) modulates neuronal excitability and network dynamics using currents delivered through electrodes placed on the scalp^3^. However, selective stimulation of deep brain structures remains challenging because increasing electric-field strength at depth generally increases exposure in superficial regions, limiting spatial focality^4, 5^. High-definition multielectrode configurations can improve cortical focality, but do not eliminate this depth– focality constraint^4, 6, 7^. Improving deep targeting with tES therefore requires strategies that increase the selectivity of neuronal responses at depth despite substantial electric-field exposure in superficial and intervening regions.

Temporal interference stimulation (TIS) was introduced as a spatiotemporal strategy to address this limitation. Two independent electrode pairs deliver kilohertz-frequency currents at slightly different carrier frequencies, generating spatially varying amplitude-modulated electric fields at the difference frequency. The modulation depth depends on the relative amplitudes and orientations of the local fields and decreases where one field dominates^8^. TIS was initially proposed to preferentially activate neurons in regions with strong modulation. Grossman et al. reported hippocampal activation with limited recruitment of the overlying cortex in mice and demonstrated steering of motor responses by adjusting the relative stimulation currents^8^. Subsequent studies optimized electrode configurations, stimulation current amplitudes and relative phase to improve deep-brain targeting^9, 10, 11, 12, 13^ and investigated TIS in humans^14, 15, 16^. Many targeting strategies aim to maximize the envelope amplitude at the target or improve its spatial focality, treating these field-based quantities as proxies for neural engagement^10, 17^. However, these field-based strategies provide limited mechanistic insight into how local stimulation waveforms produce spatially selective neuronal responses.

Early explanations of TIS proposed that neuronal membranes act as low-pass filters, responding to the slowly varying interference envelope rather than the kilohertz carriers^8^. Subsequent biophysical modeling challenged this interpretation, showing that passive membrane filtering alone cannot extract the envelope. The analysis instead suggested a role for ion-channel-mediated rectification and predicted tonic firing and conduction block in off-target axons^18^. Cortical-neuron models further showed comparable activation thresholds for modulated and unmodulated kilohertz stimulation and predicted conduction block under strong carrier-dominated stimulation, while amplitude modulation permits periodic responses^19^. Peripheral nerve experiments likewise reported largely overlapping activation thresholds and strength–frequency relationships for modulated and unmodulated kilohertz stimulation^20^. Cortical-slice recordings implicated network effects, showing that TIS-selective responses in pyramidal neurons were largely lost after synaptic blockade and proposing inhibition mediated by parvalbumin-positive (PV) interneurons as a contributor to selectivity^21^. Together, these findings challenge a simple envelope-based explanation of TIS selectivity and implicate both intrinsic neuronal dynamics and network-dependent mechanisms. The comparable activation thresholds raise the possibility that selectivity emerges from differences in neuronal responses as stimulation intensity increases beyond threshold. However, a unified, experimentally supported framework linking suprathreshold neuronal responses to the spatial selectivity and operating limits of TIS across neuronal populations and stimulation conditions remains lacking.

Here we show that differential firing suppression underlies spatial selectivity in suprathreshold TIS. Multiscale simulations revealed no consistent activation-threshold advantage for TIS over unmodulated high-frequency stimulation (HFS). At higher currents, HFS produced sustained depolarization and firing suppression, whereas strong amplitude modulation permitted periodic membrane recovery and maintained beat-locked firing, supporting selective recruitment and current-ratio-dependent steering under spatially heterogeneous electric fields. An atlas-based rat-brain model further revealed how stimulation intensity and neuronal properties shaped spatial selectivity. Whole-cell recordings under pharmacological synaptic blockade supported the predicted firing and membrane-potential dynamics, while retinal multielectrode-array recordings demonstrated differential suppression and spatial steering. In vivo motor-cortex stimulation showed stronger beat-frequency locking of motor responses under TIS than HFS. Together, these findings provide an experimentally supported framework linking suprathreshold neuronal responses to spatial selectivity, steering and intensity-dependent operating windows of TIS.

## Results

### TIS confers no activation-threshold advantage over HFS

We first tested whether TIS reduces the stimulation intensity required to activate a neuron. We constructed a simplified multiscale model consisting of a homogeneous tissue phantom, two opposing electrode pairs and a morphologically reduced, biophysically detailed hippocampal CA1 pyramidal neuron, with its soma positioned at the center of the phantom (Fig. 1a, Extended Data Fig. 1, and Supplementary Fig. 1)^22, 23^. TIS was delivered with equal-amplitude currents at 2.00 and 2.01 kHz, producing maximal amplitude modulation at the soma, whereas HFS used a common 2.00 kHz carrier with the same current ratio (Fig. 1a,b). Activation threshold was defined as the minimum total current amplitude that evoked at least one action potential (AP) during the stimulation plateau, excluding the ramp-up and ramp-down epochs. At the phantom center, the activation thresholds were 12.21 mA for TIS and 11.89 mA for HFS, indicating no activation-threshold advantage for TIS (Fig. 1b).

**Fig. 1.**
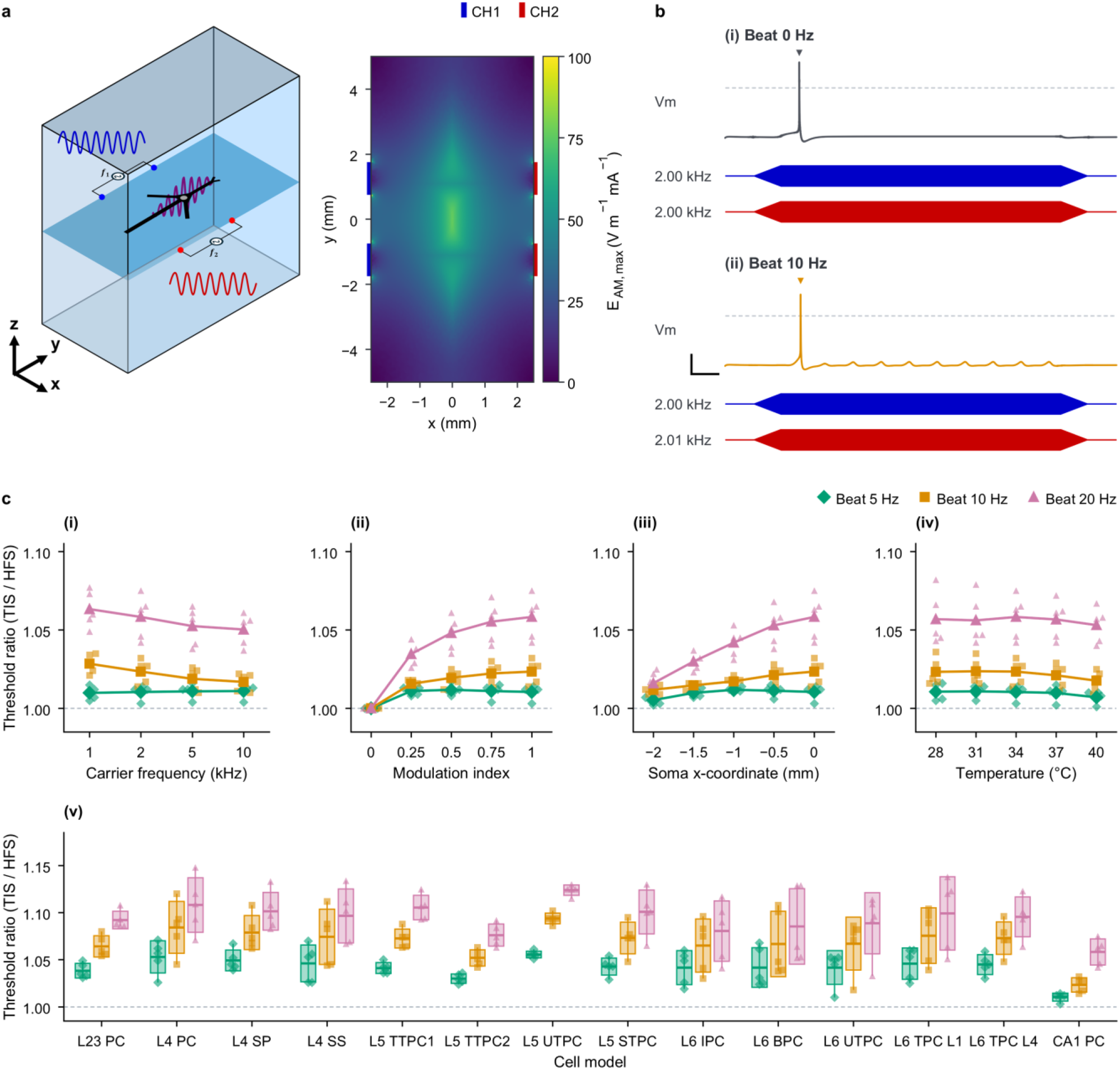
| Temporal interference stimulation has no systematic activation-threshold advantage over high-frequency stimulation. **a**, Homogeneous grey-matter phantom, electrode arrangement and maximum envelope-modulation amplitude, E_AM,max_, at z = 0, normalized by total current. Blue and red bars mark the CH1 and CH2 electrodes, respectively. **b**, Filtered somatic membrane potential (Vm) at activation threshold (Methods): (i) HFS, both channels at 2,000 Hz, 11.89 mA total; (ii) TIS, 2,000 and 2,010 Hz, 12.21 mA total. Below, normalized channel waveforms. Dashed line, 0 mV; filled triangles, plateau APs. Scale bars, 100 ms and 30 mV. **c**, TIS-to-HFS activation-threshold ratios for beat frequencies of 5 Hz (green diamonds), 10 Hz (orange squares) and 20 Hz (pink triangles), comparing the same clone under matched conditions. (i) Carrier frequency. (ii) Somatic modulation index (MI), varied through the channel-current ratio. At MI = 0, only CH1 was active. (iii) Soma x-coordinate. (iv) Temperature. (v) Thirteen excitatory cortical model types and the CA1 PC model, each represented by five clones (Supplementary Table 1). Small symbols, individual clones. In (i)–(iv), larger symbols and connecting lines show five-clone means. In (v), boxes span mean ± s.d., horizontal lines mark means and thin vertical lines span the minimum-to-maximum range across clones. Dashed line, ratio 1. Unless varied, simulations used the morphologically simplified, unmyelinated CA1 model at the phantom origin, 34 °C, a 2 kHz carrier and equal channel amplitudes.

We next assessed whether this relationship persisted across temporal, spatial and biophysical conditions (Fig. 1c). The relative activation threshold was defined as R_th = I_th(TIS)/I_th(HFS), with values above unity indicating a higher threshold for TIS (Methods). Activation thresholds increased with carrier frequency for both TIS and HFS, while R_th remained above unity across the tested carrier and beat frequencies (Fig. 1c(i) and Supplementary Fig. 2a). We next varied the modulation index (MI), with 0 representing unmodulated HFS and 1 representing maximal amplitude modulation (Methods). Increasing MI generally increased R_th, indicating that stronger amplitude modulation did not confer an activation-threshold advantage over HFS (Fig. 1c(ii)). Displacing the soma from the phantom center reduced the activation thresholds of both TIS and HFS, while R_th remained above unity across the tested positions (Fig. 1c(iii) and Supplementary Fig. 2c). For the subsequent biophysical comparisons of temperature and neuronal models, maximally modulated TIS (MI = 1) was compared with unmodulated HFS. Increasing temperature reduced the activation thresholds of both stimulation modes, whereas R_th remained above unity across all temperatures tested (Fig. 1c(iv) and Supplementary Fig. 2d). Across 14 neuronal models (Supplementary Table 1), each represented by five clones, absolute activation thresholds varied substantially, whereas R_th remained above unity for all models examined (Fig. 1c(v) and Supplementary Fig. 2e)^24^. Extending the analysis across carrier frequencies, beat frequencies and temperatures yielded the same overall relationship across the 14 models, with R_th generally increasing with beat frequency (Supplementary Fig. 3). Together, these results do not support a lower activation threshold as an explanation for TIS selectivity under the conditions examined.

### TIS maintains beat-locked firing while HFS undergoes suprathreshold suppression

Having found no activation-threshold advantage for TIS, we compared suprathreshold firing responses under TIS and HFS. Firing rates initially increased with total current under both conditions but diverged at higher amplitudes (Fig. 2a). Under HFS, firing ceased at 15.7 mA and remained absent at all higher currents tested. By contrast, TIS firing declined from its maximum and stabilized at 10 spikes/s from 17.2 mA, corresponding to one AP per beat cycle at 10 Hz. We therefore defined the TIS operating window (TOW) for each tested condition as the set of suprathreshold currents at which HFS evoked no APs during the stimulation plateau while TIS elicited beat-locked firing (Methods). Representative filtered membrane-potential traces illustrated these contrasting responses at matched stimulation currents (Fig. 2b(iii,iv)). The same divergence was observed at beat frequencies of 5 and 20 Hz (Supplementary Fig. 4). Across 14 neuronal models, TIS generally maintained higher normalized firing rates than HFS at stronger stimulation currents, although response profiles varied among models (Extended Data Fig. 2).

**Fig. 2.**
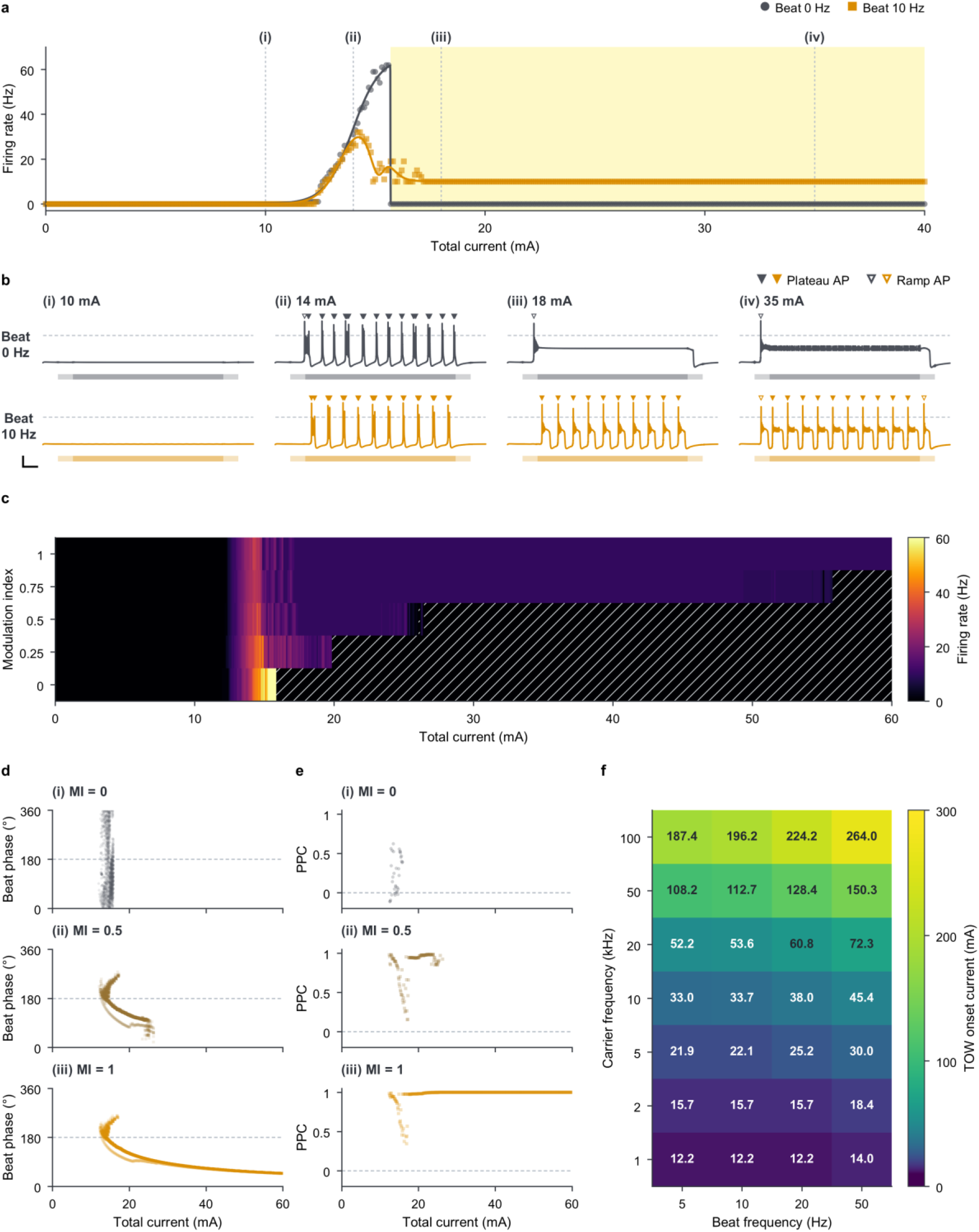
| Temporal interference stimulation maintains beat-locked somatic firing at suprathreshold currents that suppress high-frequency-stimulation responses. **a**, Firing rate versus total current. Symbols, simulation results; lines, fitted guides. Yellow shading, TIS operating window (TOW). **b**, Filtered somatic Vm at (i) 10, (ii) 14, (iii) 18 and (iv) 35 mA, corresponding to the currents marked in a; HFS above TIS. Grey and orange horizontal bars, stimulation periods; lighter segments, ramps. Dashed line, 0 mV; filled and open triangles, plateau and ramp APs, respectively. Scale bars, 100 ms and 30 mV. **c**, Firing rate across total current and somatic MI; the colour scale saturates at 60 Hz. Hatching marks zero-AP conditions above the first firing current at each MI. At MI = 0, only CH1 was active. **d**, Phase of individual plateau APs at MI = 0, 0.5 and 1. For MI > 0, 0°/360° marks the envelope minimum and 180° (dashed line) the maximum; MI = 0 uses a reference 10 Hz clock. **e**, Pairwise phase consistency (PPC) for each current with at least four plateau APs, using the APs in d; dashed line, zero. **f**, TOW onset current across carrier and beat frequencies, with logarithmic colour scaling. Unless varied, simulations used the morphologically simplified, unmyelinated CA1 model at the phantom origin, 34 °C, a 2 kHz carrier and equal channel amplitudes, with a 10 Hz beat for TIS.

To determine how amplitude modulation affected these response transitions, we mapped firing rates across total current and MI (Fig. 2c). At MI = 0, increasing current drove the neuron sequentially from a subthreshold state to firing and then to sustained suppression. As MI increased, the activation boundary changed comparatively little, whereas the suppression boundary generally shifted to higher currents. At MI = 1, sustained suppression was not reached within the tested current range. Thus, increasing modulation primarily delayed the onset of sustained suppression rather than lowering the activation threshold.

We next examined how stimulation current and MI affected spike timing within the beat cycle (Fig. 2d and Methods). For modulated waveforms (MI > 0), spikes near the activation threshold occurred preferentially around the envelope maximum at 180°. As stimulation current increased, the preferred firing phase shifted from the envelope maximum toward the envelope minimum near 0° and 360°. Pairwise phase consistency (PPC) approached unity over a broad current range at MI = 1, indicating strong phase locking to the beat cycle (Fig. 2e and Methods)^25^. As MI decreased, the current range supporting phase-locked firing narrowed because sustained suppression occurred at lower stimulation currents. Simulations using a slowly varying envelope showed suppression at higher envelope amplitudes and firing at lower amplitudes, consistent with the shift in firing towards the envelope minimum at stronger stimulation currents (Extended Data Fig. 3).

Finally, we examined the frequency dependence of the TOW onset across carrier frequencies of 1–100 kHz and beat frequencies of 5–50 Hz, with TIS evaluated at MI = 1 (Fig. 2f). The TOW onset increased markedly with carrier frequency, from approximately 12.2–14.0 mA at 1 kHz to 187.4–264.0 mA at 100 kHz, depending on beat frequency. The onset also generally increased with beat frequency, with this dependence becoming more pronounced at higher carrier frequencies. Together, these results indicate that the TOW was not restricted to a specific frequency combination, although its onset depended on both carrier and beat frequencies.

### Suprathreshold suppression enables spatially selective steering

We next tested whether the response transitions identified in Fig. 2 could support spatially selective firing and steering. We varied the current ratio between the two stimulation channels while matching the total current across steering conditions. The MI maximum shifted from x = 0 mm at a current ratio of 10:10 to approximately 1 and 2 mm at ratios of 16.4:3.6 and 19:1, respectively (Fig. 3a and Extended Data Fig. 4). For each current ratio, we defined the intended target as the region along the x-axis where MI ≥ 0.9, with all remaining sampled locations classified as off-target (MI < 0.9). The target length remained nearly constant across most current ratios but increased when the channel currents became highly unequal (Extended Data Fig. 4b).

**Fig. 3.**
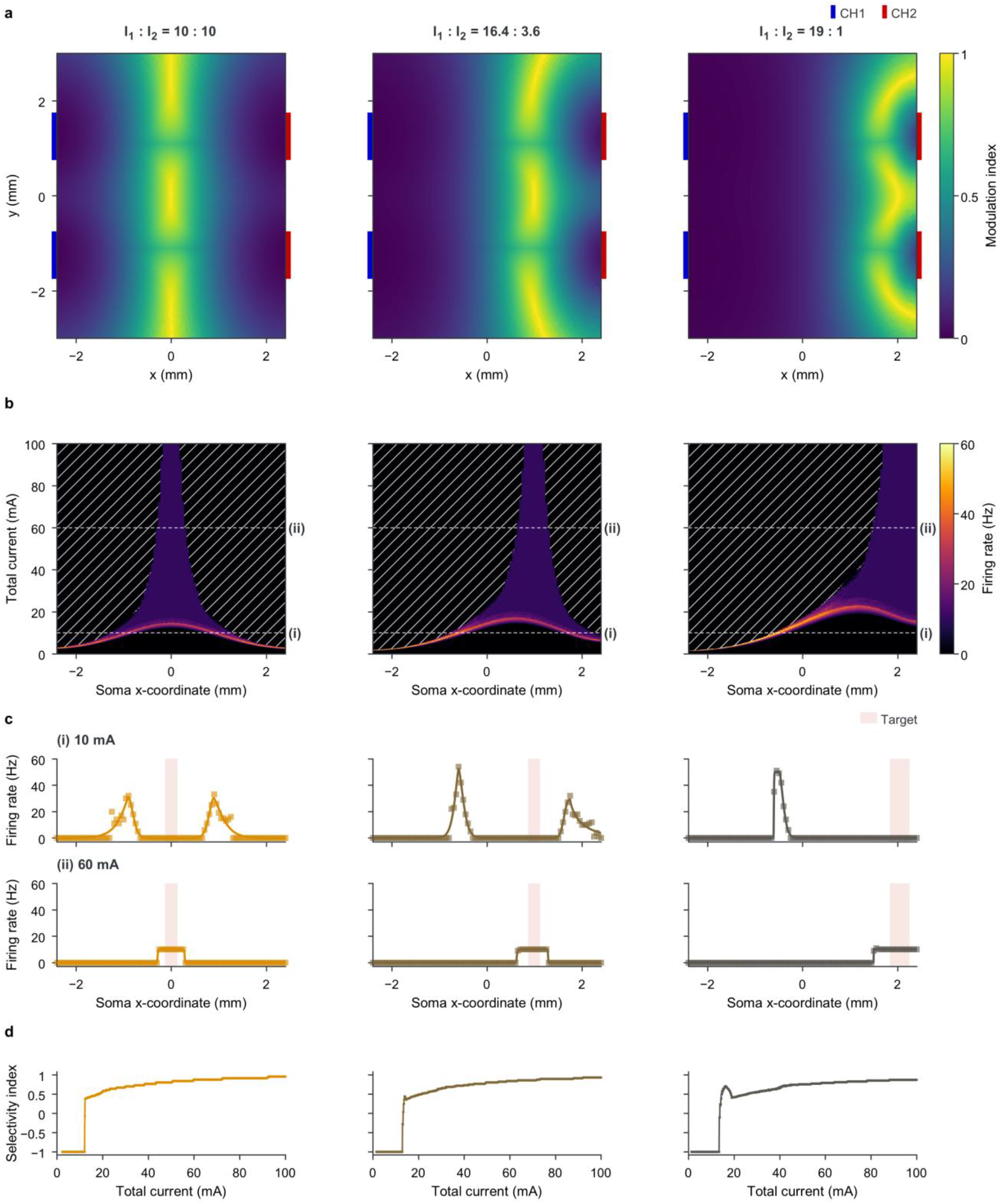
| Suprathreshold suppression supports spatially selective firing and current-ratio-dependent steering. Columns show CH1:CH2 current ratios of 10:10, 16.4:3.6 and 19:1, with total current matched across ratios at each tested amplitude. **a**, MI maps at z = 0; blue and red bars mark the CH1 and CH2 electrodes. **b**, Somatic firing rate across soma x-coordinate and total current. The colour scale saturates at 60 Hz. Hatching marks zero-AP conditions above the first firing current at each position; dashed lines mark 10 and 60 mA. **c**, Spatial firing-rate profiles at (i) 10 and (ii) 60 mA. Symbols, simulation results; lines, fitted guides; red shading, the MI-defined target (MI ≥ 0.9). **d**, Selectivity index (SI) versus total current; corresponding target and off-target activation fractions are shown in Extended Data Fig. 4. All simulations used the morphologically simplified, unmyelinated CA1 model at 34 °C, with CH1 and CH2 driven at 2,000 and 2,010 Hz. The soma was placed in the phantom at x-coordinates from −2.4 to 2.4 mm in 0.05 mm steps (y = z = 0), with a fixed neuronal orientation.

We mapped firing rates across neuronal position and total current to determine how these modulation patterns translated into spatially selective responses (Fig. 3b). At lower currents, firing was concentrated in off-target regions near the electrodes. As current increased, firing emerged within the strongly modulated target region, while previously active neurons in weakly modulated off-target regions entered sustained suppression. For the 10:10 current ratio, this reversal in the spatial firing pattern was observed within the TOW (Fig. 2a). At higher currents, the region where firing was maintained shifted with the current ratio (Fig. 3b). At 10 mA, no firing was observed within the target, whereas at 60 mA, target firing reached approximately 10 spikes/s for all three current ratios (Fig. 3c).

To quantify spatial selectivity, we calculated a selectivity index (SI) from the target and off-target activation fractions, with −1 indicating exclusively off-target activation and +1 indicating exclusively target activation (Methods). The activation fraction was the proportion of sampled locations in each region that generated at least one AP during the stimulation plateau. The target activation fraction reached unity and remained saturated at higher currents for all three current ratios (Extended Data Fig. 4d). By contrast, the off-target activation fraction initially increased and subsequently declined with increasing current. The SI showed an overall increase at higher suprathreshold currents (Fig. 3d). Varying the MI cutoff from 0.80 to 0.95 altered target extent and SI values but preserved the overall increase in selectivity at higher currents (Supplementary Fig. 5). With target recruitment already saturated, this improvement in spatial selectivity primarily reflected suppression of off-target firing.

### Persistent depolarization under HFS and periodic repolarization during TIS

We next examined the membrane dynamics underlying sustained HFS suppression and beat-locked TIS firing. At a total current of 20 mA, removing the carrier-frequency oscillations from the membrane-potential traces revealed sustained depolarization under HFS and alternating depolarized and hyperpolarized intervals under TIS, despite the zero-mean stimulation waveforms (Fig. 4a,b). The sodium-channel inactivation variable h showed a corresponding temporal pattern (Fig. 4c). Under HFS, h remained low, indicating reduced sodium-channel availability. Under TIS, h remained low during depolarized intervals but recovered during hyperpolarized intervals.

**Fig. 4.**
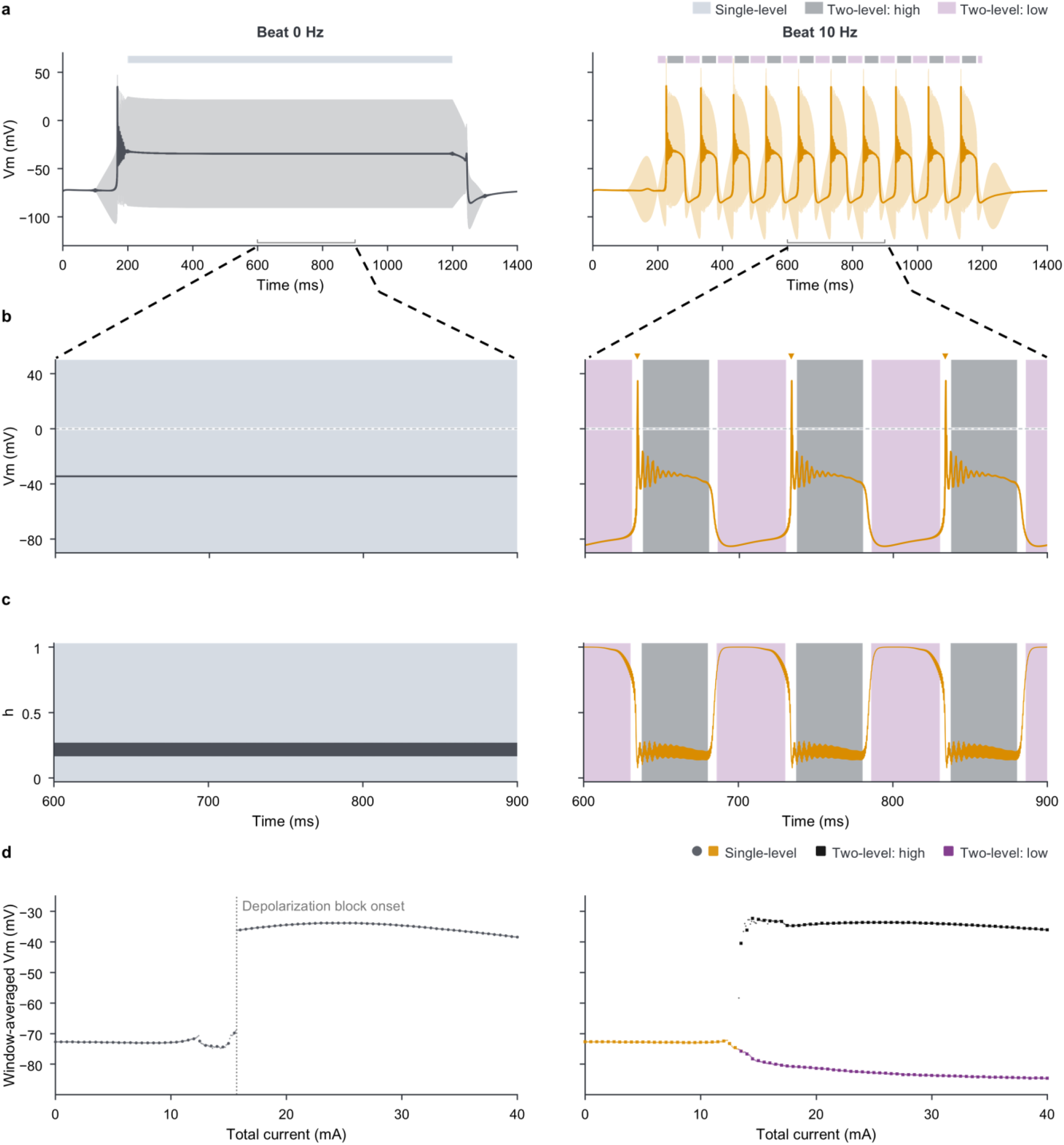
| Persistent depolarization under high-frequency stimulation contrasts with periodic recovery under temporal interference stimulation. HFS and TIS are shown in the left and right columns, respectively; a–c use 20 mA total current. **a**, Raw (pale) and filtered (dark) somatic Vm. Bands above the traces, single-level, high and low averaging windows used in d; gaps, excluded AP masks and transition intervals. Brackets, interval enlarged in b and c. **b**, Filtered Vm during 600–900 ms, with the same window colours. Dashed line, 0 mV; triangles, detected APs. **c**, Somatic Na⁺-channel inactivation gate h; h = 1 indicates no inactivation. **d**, Window-averaged raw Vm versus total current, using the window-selection procedure in Supplementary Fig. 6. Left, HFS single-level means; dotted vertical line, onset of somatic depolarization block under these conditions (15.7 mA; Fig. 2a). Right, orange squares, single-level means at currents without two-level beats; black squares, high-window means; purple squares, means pooled across single-level and low windows at currents containing two-level beats. Window categories describe the analysis and do not by themselves identify physiological states. Small dots, every simulated current (0–40 mA, 0.1 mA steps); larger markers, currents at 0.5 mA intervals. All simulations used the morphologically simplified, unmyelinated CA1 model at the phantom origin, 34 °C, a 2 kHz carrier and equal channel amplitudes, with a 10 Hz beat for TIS.

To examine how these membrane states changed with stimulation current, we quantified window-averaged membrane potentials during the stimulation plateau (Fig. 4d). For TIS, high- and low-potential windows were analyzed separately in beats classified as having two distinct potential levels (Supplementary Fig. 6 and Methods). Under HFS, the window-averaged membrane potential remained near the resting level at lower currents before shifting abruptly to a sustained depolarized level at 15.7 mA, coinciding with the onset of firing suppression (Fig. 2a). Sustained depolarization and loss of firing, accompanied by reduced sodium-channel availability, were consistent with depolarization block^26^. Under TIS, the high-level mean approached the sustained depolarized level observed under HFS as current increased, whereas the low-level mean became progressively more negative (Fig. 4d). These model dynamics support a contribution of periodic repolarization and recovery of sodium-channel availability in maintaining beat-locked firing during TIS at currents that suppress HFS responses.

### Suprathreshold response transitions support selective recruitment in an atlas-based brain model

We next extended the analysis to a rat-brain model based on the Waxholm Space atlas, incorporating morphologically detailed neurons in hippocampal CA1 and layer 4 of the primary somatosensory barrel cortex (S1-bf) (Fig. 5a,b)^27^. Neuronal models were placed within both regions and oriented according to local anatomy (Fig. 5b and Methods). The electrode montage produced a high-MI region intersecting CA1, but MI varied substantially within CA1 (Fig. 5a and Extended Data Fig. 5a). To evaluate selectivity for the strongly modulated CA1 subregion rather than CA1 as a whole, we defined the target population as CA1 neurons with a local MI ≥ 0.9 at the soma and classified all remaining modeled neurons in CA1 and S1-bf as off-target.

**Fig. 5.**
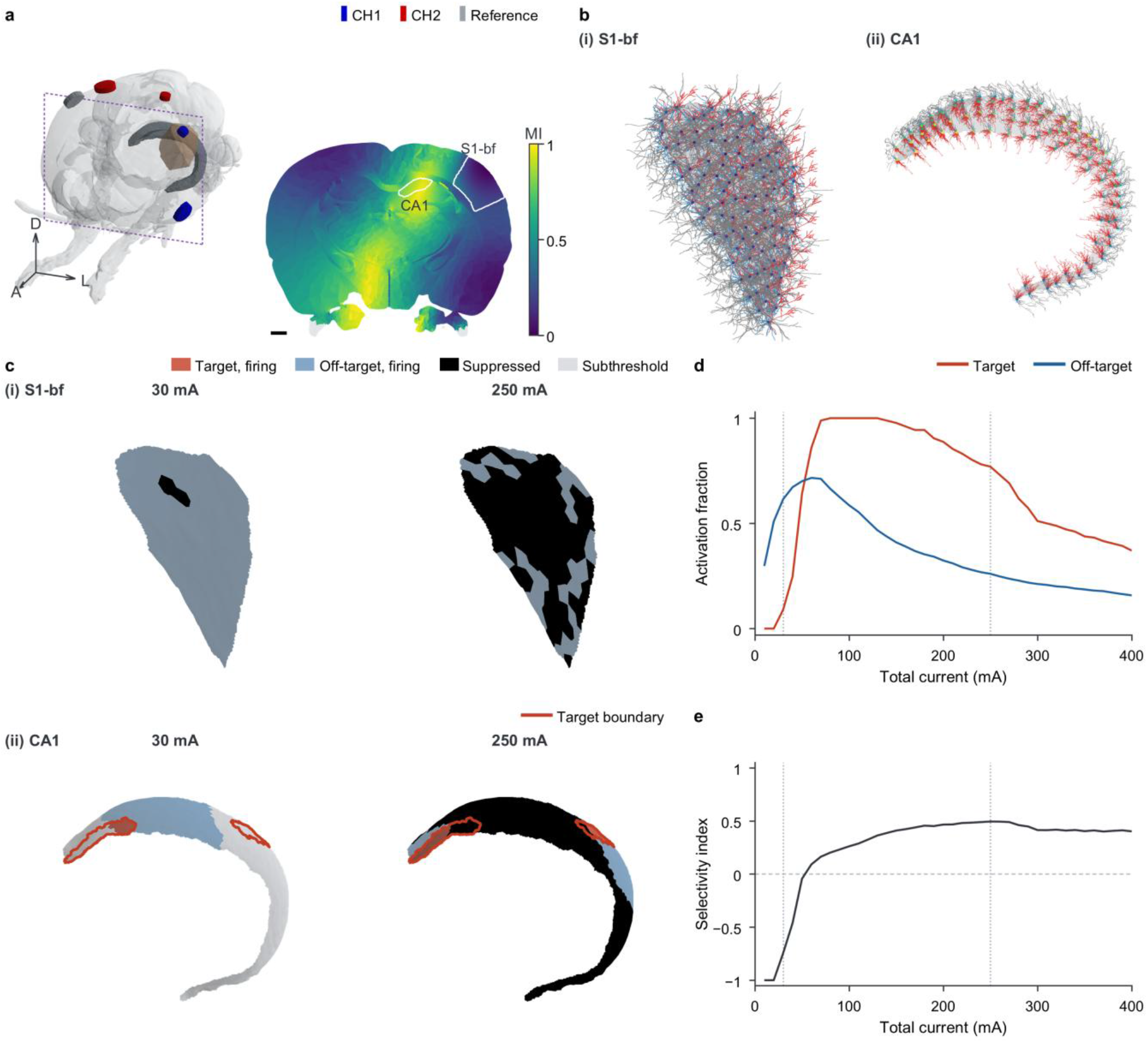
| Current-dependent target recruitment and spatial selectivity in an atlas-based rat-brain model. **a**, Brain-surface montage and coronal MI map at y = −2.0 mm, excluding scalp and skull. Grey map regions, undefined MI. Blue, red and grey electrodes, CH1, CH2 and reference, respectively. Scale bar, 1 mm. **b**, Detailed morphologies of (i) L4 PC clone 2 at 100 positions in S1-bf layer 4 and (ii) CA1 PC clone 5 at 100 of 1,000 simulated CA1 positions (Supplementary Tables 2 and 3). Apical dendrites, red. Basal dendrites, blue. Axons, grey. Soma colour indicates MI. **c**, Firing states at 30 and 250 mA for the models in b, mapped to surfaces by nearest simulated soma. Red and blue, firing target and off-target positions, respectively. Black, no plateau APs after firing at a lower tested current. Grey, positions below their first firing current. Red outlines enclose CA1 targets (somatic MI ≥ 0.9). All remaining CA1 and S1-bf L4 positions were off-target. **d**, Volume-weighted target and off-target activation fractions pooled across two CA1 clones (1,000 positions each) and nine L4 clones (100 positions each, three orientations per position). **e**, SI from the fractions in d, maximal at 250 mA among tested currents. Vertical dotted lines in d,e mark the currents in c. TIS used equal channel amplitudes at 2,000 and 2,010 Hz and total currents of 10–400 mA.

At 30 mA, firing occurred predominantly in the off-target population, whereas neurons within the high-MI CA1 target remained largely subthreshold (Fig. 5c). At 250 mA, many off-target neurons that had fired at lower currents were suppressed, whereas a large fraction of neurons within the high-MI CA1 target were in the firing state. This heterogeneity was evident even within CA1, where the activation fractions at 250 mA were 0.75 and 0 for the top and bottom 10% of locations ranked by soma-level MI, respectively (Extended Data Fig. 5a,b). The current-dependent recruitment patterns also varied among neuronal model types (Extended Data Fig. 5c).

We quantified recruitment across stimulation currents using volume-weighted activation fractions for the target and off-target populations (Fig. 5d). The off-target activation fraction initially increased and subsequently declined with increasing current. The target activation fraction increased at higher currents, reached unity and subsequently declined as some target neurons also ceased firing. Accordingly, the SI increased from −0.75 at 30 mA to a maximum of approximately 0.49 at 250 mA, before decreasing slightly to 0.40 at 400 mA (Fig. 5e). Notably, maximal spatial selectivity occurred at a higher stimulation current than maximal target recruitment, reflecting a further reduction in off-target activation relative to target activation.

### Whole-cell recordings support modulation-dependent suprathreshold responses ex vivo

To determine whether the modulation-dependent suprathreshold responses predicted computationally were reproduced ex vivo, we performed whole-cell current-clamp recordings from five layer 4 neurons in acute mouse S1-bf slices under pharmacological synaptic blockade (Fig. 6a). To control MI without relying on spatial interference, we applied premodulated waveforms through a single electrode pair. HFS, half-TIS and TIS were applied with MI values of 0, 0.5 and 1, respectively. At a representative suprathreshold current of 300 µA, HFS elicited transient APs during ramp-up but no APs during the stimulation plateau. By contrast, half-TIS and TIS maintained repetitive firing during the plateau, with TIS responses occurring approximately once per beat cycle (Fig. 6b).

**Fig. 6.**
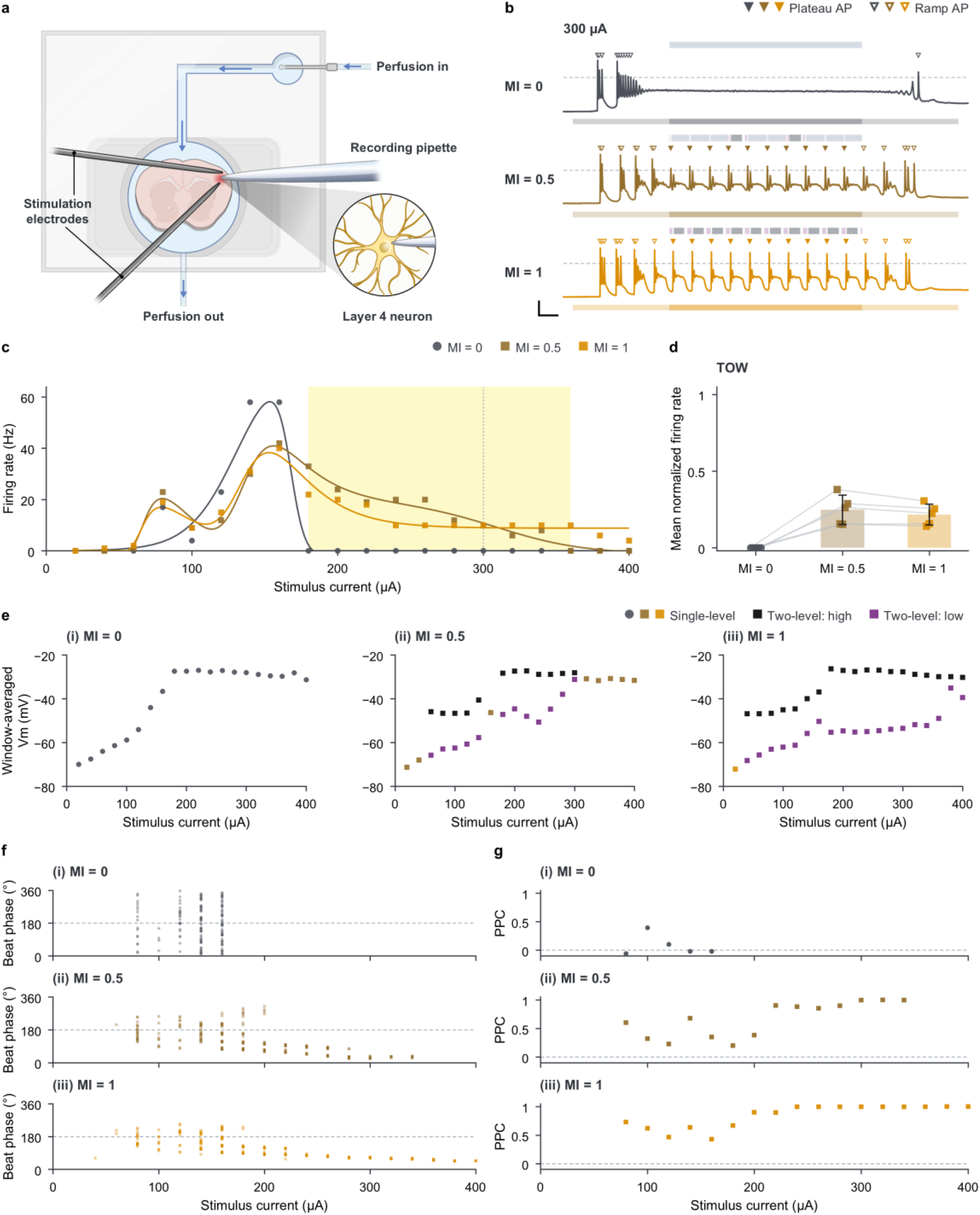
| Temporal interference stimulation maintains beat-locked firing at suprathreshold currents that suppress high-frequency-stimulation responses in brain slices. **a**, Extracellular stimulation and whole-cell current clamp of S1-bf layer 4 neurons under synaptic blockade in slices from male C57BL/6J mice aged 8–12 weeks. Panels b, c and e–g show Cell 4. **b**, Filtered V_m_ at 300 µA. Lower bars, stimulation with lighter ramps. Dashed lines, 0 mV. Filled/open triangles, plateau/ramp APs. Upper bands, single-level (pale grey), high (grey) and low (pale purple) averaging windows for e. Gaps, excluded AP masks and transitions. Scale bars, 100 ms and 30 mV. **c**, Plateau firing rate versus current. Symbols, sweeps. Lines, fitted guides. Dotted line, 300 µA. Yellow, TOW. **d**, Rates normalized to each cell’s maximum across waveforms and currents, then averaged per waveform over TOW currents valid for all waveforms. Linked symbols, cell means. Bars and error bars, mean ± s.d. (five cells, one per mouse). **e**, Window-averaged raw V_m_ (Supplementary Fig. 6). Waveform colours, means from exclusively single-level sweeps. For sweeps containing two-level segments, black, high-window means, and purple, pooled single-level/low-window means. **f**, Plateau AP phases on a common 10 Hz clock from plateau onset. Dashed line, 180°. **g**, PPC per sweep with ≥4 plateau APs. Dashed line, zero. One electrode pair delivered unmodulated 2 kHz stimulation (MI = 0), then 10 Hz premodulation (MI = 0.5, then 1), one sweep per waveform per current (20 µA increments). Ramps lasted 0.5 s each around a 1 s plateau.

We next quantified firing across stimulus currents (Fig. 6c). In the representative neuron, firing rates initially increased under all three waveforms but diverged at higher currents. Under HFS, firing declined abruptly to zero at 180 µA. Half-TIS firing declined gradually after its peak, whereas TIS maintained firing near the 10 Hz beat frequency over an extended suprathreshold range. Similar response transitions were observed across all five neurons, although the corresponding current ranges varied among cells (Extended Data Fig. 6). To compare responses across cells, normalized firing rates were averaged over each cell’s TOW. Mean normalized firing within these windows was nonzero under half-TIS and TIS, whereas HFS firing was absent (Fig. 6d).

Window-averaged membrane potentials differed across the stimulation waveforms (Fig. 6e). Under HFS, the window-averaged mean membrane potential shifted to a strongly depolarized level at stimulation currents associated with sustained suppression. For half-TIS and TIS, the window-averaged membrane potential was strongly depolarized during high-level windows and returned towards less-depolarized values during low-level windows. At higher currents, however, half-TIS transitioned to a single depolarized level, whereas TIS retained distinct high- and low-level window-averaged potentials (Fig. 6e). Similar membrane-potential dynamics were observed across the recorded neurons, although the transitions occurred at different applied currents (Extended Data Fig. 6). These membrane-potential dynamics were consistent with the persistent depolarization during HFS and periodic return towards a less-depolarized state predicted by the model.

We next quantified how spike timing changed with stimulation current under the three waveforms (Fig. 6f,g). For half-TIS and TIS, spikes at lower currents occurred preferentially near the envelope maximum. As stimulation current increased, the preferred firing phase shifted towards the envelope minimum. PPC approached unity at higher currents for the modulated waveforms, indicating strong phase locking to the beat cycle, whereas HFS showed little phase consistency relative to a nominal 10 Hz reference cycle. Similar current-dependent shifts in firing phase were observed in the other recorded neurons, although the strength of phase locking varied among cells (Extended Data Fig. 7). These shifts were consistent with the computational predictions in Fig. 2.

Finally, depolarizing current-step tests performed before and after the stimulation protocol showed that all five neurons remained capable of generating APs after stimulation (Supplementary Fig. 7). Sustained suppression was therefore not attributable to an irreversible loss of AP generation. Together, the firing-rate, membrane-potential and spike-timing measurements supported the predicted separation between sustained HFS suppression and beat-locked firing under modulated stimulation.

### Ex vivo retinal recordings reveal differential suppression and current-ratio-dependent spatial steering

To examine whether modulation-dependent firing suppression could support spatially patterned activity in retinal tissue, we performed ex vivo multielectrode-array (MEA) recordings from isolated mouse retinas in the presence of the ionotropic glutamate receptor antagonists (Fig. 7a). Unlike the premodulated waveforms used in the whole-cell experiments, TIS was generated by superimposing electric fields at 2.00 and 2.01 kHz from two bipolar electrode pairs, producing spatially varying modulation at a 10 Hz beat frequency. Representative recordings showed that spiking persisted in the TIS region but was markedly suppressed in the HFS region, with activity recovering after stimulus offset (Fig. 7b). Firing-rate heatmaps further showed that suppression became more pronounced with increasing stimulation current, particularly under HFS (Fig. 7c). Mean firing counts measured from 0.5 to 1.5 s after stimulus onset approached zero above approximately 1.5 mA under HFS, whereas spiking remained detectable under TIS at the highest current tested (Fig. 7d).

**Fig. 7.**
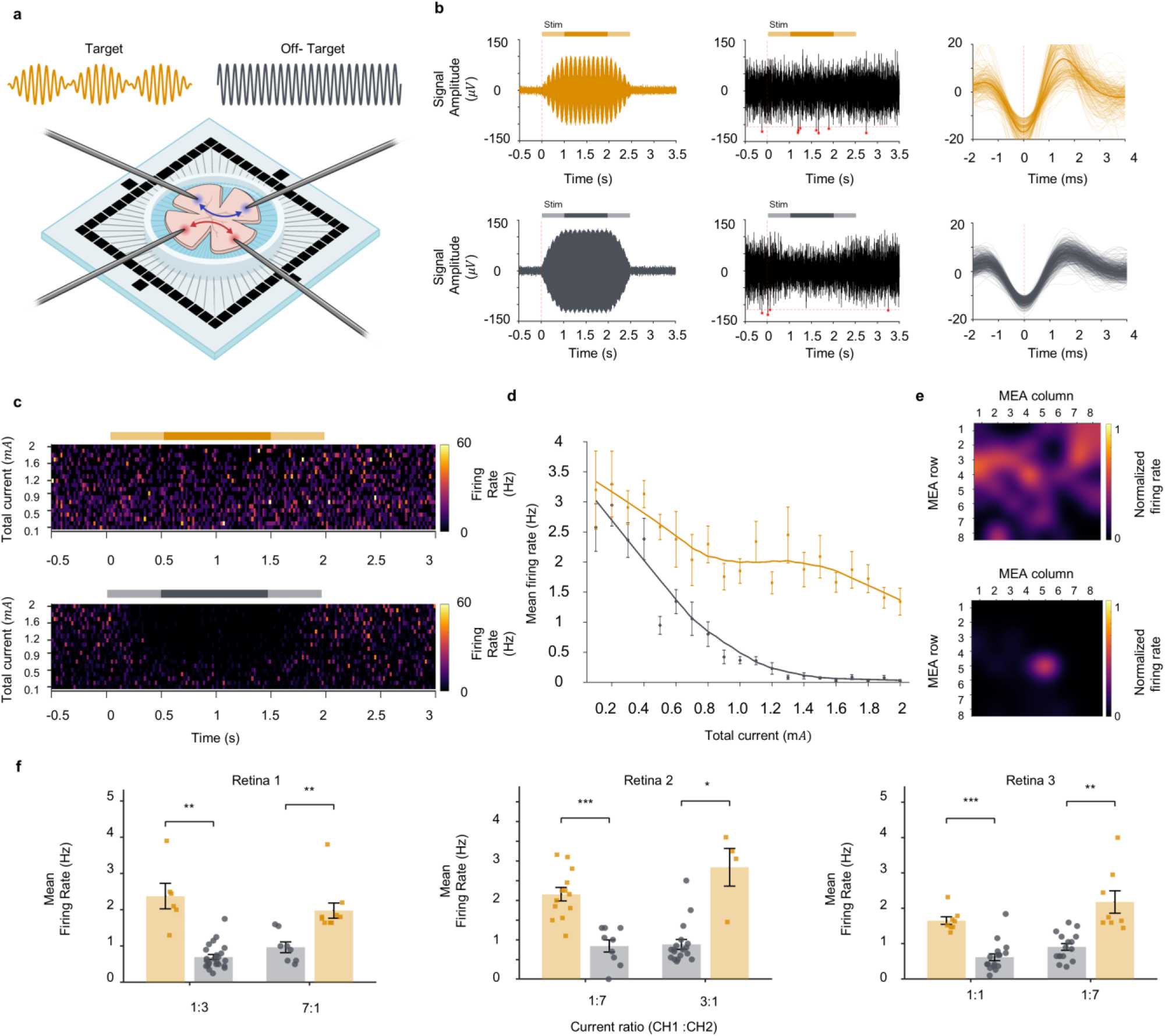
| Retinal recordings reveal differential suppression and spatial steering during temporal interference stimulation. Ex vivo MEA recordings from three isolated mouse retinae, one per mouse, each tested at two channel-current ratios. Orange and grey, recordings from the designated TIS and HFS regions, respectively, at each current ratio. **a**, Schematic of the experimental setup. Two electrode pairs delivered carriers at 2,000 and 2,010 Hz, generating spatially varying modulation with a 10 Hz beat. **b**, Representative extracellular recordings from TIS (top) and HFS (bottom) regions. Left, raw signals; middle, filtered signals with detected spikes marked by red asterisks; right, overlaid spike waveforms (thin traces, individual spikes; thick traces, means). **c**, Representative firing-rate heatmaps versus time and total current; TIS region above HFS region. Orange and grey horizontal bars in b and c, stimulation periods. **d**, Mean firing rate during the 1 s stimulation plateau versus total current, summarized across six retina–current-ratio combinations from three retinae. **e**, Spatial maps of normalized firing rate across the MEA in one retina at 200 µA (top) and 1,500 µA (bottom). **f**, Mean firing rates in two recording regions at the indicated channel-current ratios and a fixed total current of 1,500 µA, shown separately for each of the three retinae. Symbols, individual electrode measurements; bars, means. Error bars in d and f, s.e.m. Statistical comparisons between TIS and HFS regions in f used two-sided Welch’s two-sample t-tests; *\*p < 0.05, **p < 0.01 and ***p < 0.001*.

We next examined whether differential firing suppression could produce spatially selective activity and whether the location of retained firing could be steered by adjusting the current ratio. Spatial firing-rate maps showed broadly distributed activity at a lower current and a more localized region of retained activity at a higher current (Fig. 7e). At a fixed total current of 1.5 mA, adjusting the ratio between the two stimulation channels changed which of the two recording regions had the higher mean firing rate in all three retinal preparations (Fig. 7f).

### TIS increases beat-frequency locking of motor output in vivo

To examine how the timing of forelimb movements differed between HFS and TIS, we delivered HFS and 10 Hz TIS through the same bipolar electrode in the caudal forelimb area (CFA) of rat motor cortex following local injection of synaptic receptor antagonists and tracked contralateral digit-2 movements using DeepLabCut (Fig. 8a and Methods)^28, 29^. For each animal, movement trajectories under both waveforms were projected onto a shared principal movement axis. Movement-event timing was then evaluated relative to a common 100 ms reference cycle (Extended Data Fig. 8 and Methods).

**Fig. 8.**
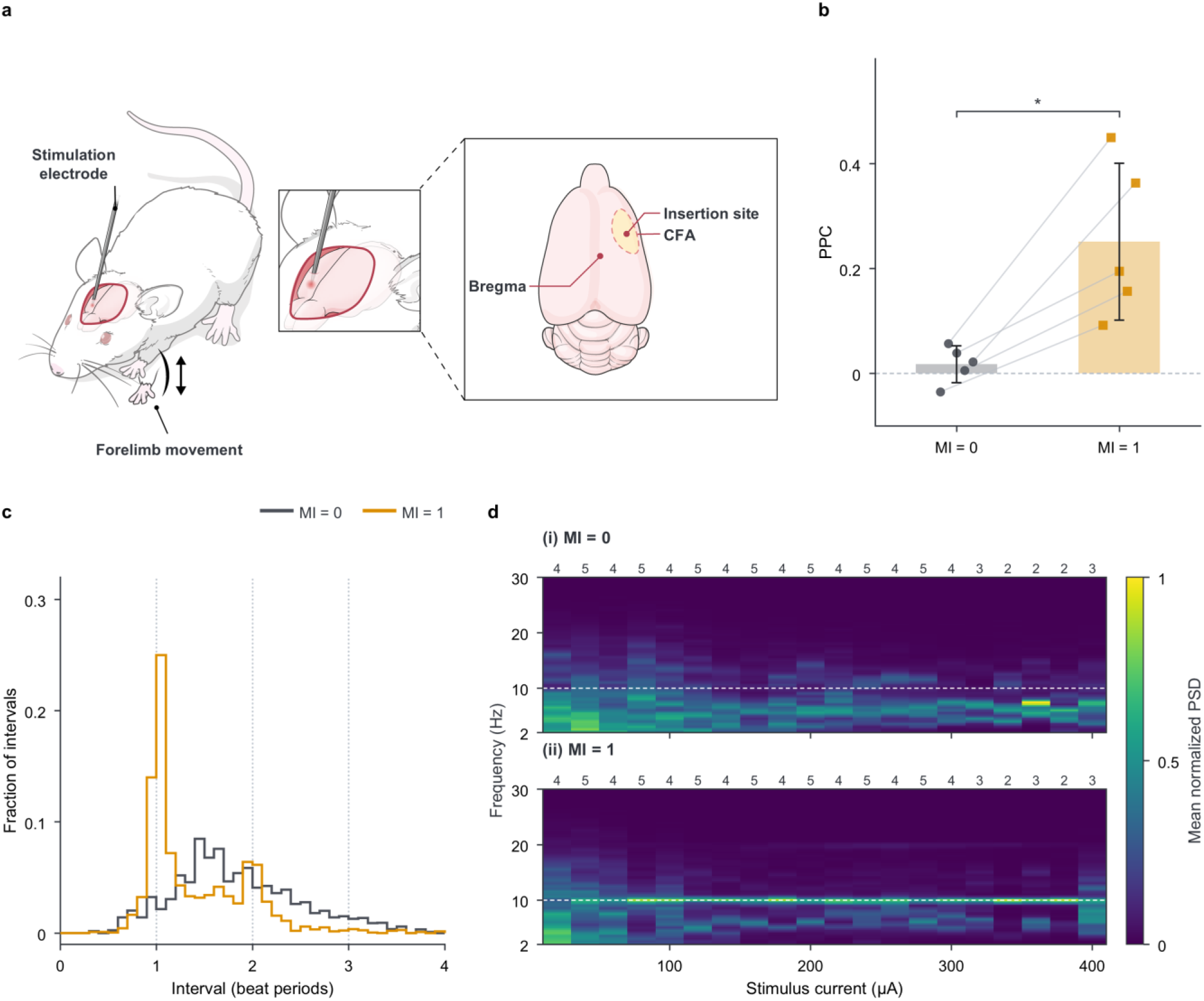
| Temporal interference stimulation enhances beat-frequency locking of forelimb movements in vivo. **a**, Cortical stimulation and contralateral forelimb movement in five rats. Digit 2 of the left forepaw was tracked. **b**, Movement-event PPC averaged per rat across matched currents with ≥4 plateau events under each waveform. Phases used a common 10 Hz clock from stimulation onset. Linked symbols, individual rat means. Bars and error bars, mean ± s.d. (n = 5). Dashed line, zero. Two-sided paired t-test with Holm correction across b and both Extended Data Fig. 9b comparisons. Adjusted *p* = 0.040. Asterisk, *p* < 0.05. **c**, Inter-event intervals in 10 Hz reference periods. Step histograms, mean interval distributions with equal weight per rat. Vertical dotted lines, one, two and three periods. **d**, Power spectral density (PSD) of movement-axis velocity versus current. Each epoch’s spectrum was normalized to its maximum over 2–30 Hz before averaging across available rats at each current. Upper and lower maps, HFS (MI = 0) and TIS (MI = 1), respectively. Column labels, numbers of contributing rats. Dashed line, 10 Hz. A single bipolar electrode in the right caudal forelimb area (CFA) delivered HFS at 2 kHz and TIS at 2,000 and 2,010 Hz. Peak amplitudes spanned 20–400 µA across rats. Each waveform–current condition comprised one 5 s train with 0.5 s ramps around a 4 s plateau. Analyses in b–d used the 0.5–4.5 s plateau.

Both HFS and TIS elicited motor responses, but the timing of movement events differed between stimulation conditions. PPC averaged across matched stimulation currents was higher under TIS than HFS in all five rats (0.251 versus 0.017; paired t-test, Holm-adjusted p=0.040, Fig. 8b), indicating stronger phase locking of movement events to the 10 Hz beat cycle. Consistent with this finding, intervals between consecutive movement events were concentrated near the 100 ms beat period under TIS but were more broadly distributed under HFS (Fig. 8c and Extended Data Fig. 8e). Normalized velocity spectra further showed greater concentration near 10 Hz under TIS across stimulation currents (Fig. 8d). The 9–11 Hz band accounted for 38.6% of the 2–30 Hz power under TIS, compared with 8.5% under HFS (Extended Data Fig. 9). Together, these results demonstrate stronger beat-frequency locking of motor output under TIS.

## Discussion

Our findings support differential suprathreshold suppression as a mechanism for spatially selective firing during TIS. Across the simulated conditions, strong amplitude modulation did not confer a consistent activation-threshold advantage over unmodulated HFS. Instead, the principal effect emerged above threshold, where HFS produced sustained firing suppression at higher stimulation currents while strongly modulated stimulation maintained beat-locked firing with periodic repolarization. Spatial variation in modulation then separated target and off-target neurons into different suprathreshold response states, enabling selective firing and current-ratio-dependent steering through preferential off-target suppression.

A cortical-neuron modeling study by Wang et al. reported comparable activation thresholds for modulated and unmodulated kilohertz stimulation and suggested that amplitude modulation could periodically interrupt conduction block, allowing repeated onset and offset responses^19^. Our study extends these findings by quantifying spatial recruitment, current-ratio-dependent steering and intensity-dependent operating windows under nonuniform fields, supported by complementary cellular and tissue recordings. The beat-locked neuronal firing reported in the original demonstration of TIS by Grossman et al. resembles the responses observed in our simulations and whole-cell recordings^8^ (Supplementary Fig. 8). This resemblance may reflect similar suprathreshold responses, although differential suppression was not directly examined as the basis of spatial selectivity in the original study.

An important implication of these findings is that neuronal responses to TIS cannot be explained simply by preferential excitation at the envelope maximum. In the representative CA1 model, simulations with a slowly varying envelope identified distinct boundaries for the onset of firing, the onset of sustained suppression and the re-emergence of firing after suppression (Extended Data Fig. 3). This response pattern explains the intensity-dependent shift in firing phase observed in simulations and whole-cell recordings. Near threshold, firing occurred preferentially around the envelope maximum. As stimulation intensity increased, however, firing shifted towards lower-envelope phases, consistent with suppression at higher envelope amplitudes and the resumption of firing as the envelope declined. Although the beat frequency determines the recurrence of the modulation cycle, intensity-dependent transitions between firing and suppression determine whether and when spikes occur within each cycle.

The response-state dynamics identified in this study establish an intrinsic cellular mechanism for the differential responses to modulated and unmodulated kilohertz stimulation. In whole-cell recordings under pharmacological synaptic blockade, modulated stimulation maintained firing at intensities that suppressed responses to HFS (Fig. 6). However, previous cortical-slice studies have shown that network interactions can also contribute to TIS-selective responses^21^. In intact tissue, TIS responses may therefore reflect both intrinsic membrane dynamics and network-dependent effects, with their relative contributions varying across neuronal populations, brain regions and circuit architectures.

In the atlas-based rat-brain model, spatial selectivity was influenced not only by the MI distribution but also by stimulation intensity and neuronal model type. Neuronal model types with distinct morphologies and intrinsic membrane properties exhibited different current-dependent recruitment profiles (Extended Data Fig. 5c). Stimulation intensity further shaped spatial selectivity, as increasing current altered the balance between target firing and off-target suppression despite an unchanged MI distribution. Consequently, maximal spatial selectivity occurred at a higher current than maximal target recruitment, reflecting a further reduction in off-target activation (Fig. 5d,e). Retinal MEA recordings in the presence of glutamatergic receptor antagonists provided complementary evidence for spatially patterned activity, showing more localized firing at higher currents and current-ratio-dependent changes in the region of greater retained activity (Fig. 7). Together, these findings indicate that optimizing spatial selectivity requires consideration of local field distributions, stimulation intensity and neuronal properties.

Compartment-resolved simulations in the representative CA1 neuron model revealed that the onset of firing suppression differed between the soma and axonal compartments under HFS (Extended Data Fig. 10 and Supplementary Fig. 9). At lower stimulation currents, repetitive firing was observed at the soma. As stimulation current increased, somatic firing was suppressed while APs remained detectable along the axon, indicating that somatic suppression did not necessarily coincide with the loss of axonal activity. At still higher currents, AP detections persisted at proximal axonal sites but disappeared distally. A local reduction in excitability associated with sustained suppression could impair the propagation of APs, potentially contributing to the conduction block reported in previous studies^30, 31^. Somatic and proximal compartments near the stimulation electrode could experience fields sufficient to induce sustained suppression, whereas more distant axonal segments may experience weaker effective stimulation. These compartment-dependent responses suggest that somatic suppression does not necessarily eliminate axonal activity. Although HFS also elicited motor responses in vivo, the contribution of preserved axonal activity to these responses remains unresolved.

The translational potential of TIS depends on the stimulation regime. In the suprathreshold regime examined here, beat-locked firing provides a basis for temporal control of evoked neuronal activity. Building on this finding, future studies could investigate whether dynamically varying the beat frequency enables programmable control of interspike intervals and whether the resulting temporal patterns can support information encoding in neural circuits. By contrast, human transcranial TIS is expected to operate predominantly in the subthreshold regime, where stimulation may modulate the timing of ongoing neuronal activity without inducing the firing and suppression transitions characterized here. Consistent with this distinction, non-human primate recordings at human-relevant field strengths demonstrated changes in spike timing without substantial changes in firing rate^32^. Translating TIS into effective neuromodulation strategies will therefore require approaches tailored to these distinct stimulation regimes, including the development of suprathreshold temporal stimulation strategies and characterization of subthreshold modulation under clinically relevant conditions.

## Methods

### Finite-element and neuronal models

We used SimNIBS^33^ to calculate channel-specific extracellular potentials in a homogeneous grey-matter phantom and a rat-brain model derived from the Waxholm Space atlas^27^ (Supplementary Fig. 1). The brain model excluded scalp and skull. Isotropic conductivities were assigned by tissue class^34^. The cerebrospinal-fluid value was used for the cochlea and vestibular apparatus^35^ and the white-matter value for the spinal-cord label^36^ (Supplementary Table 2A). Potentials at each neuronal segment were scaled by the channel currents and summed. Local MI was the maximum envelope-modulation amplitude divided by the sum of the two channel-field magnitudes^8, 37^. Locations with weak fields from either channel were masked in Fig. 5a.

We obtained models of 13 excitatory cell types in rat somatosensory cortex and one CA1 pyramidal cell type from the Blue Brain Project^22, 23, 24^. Each cell type was represented by five model instances with unmyelinated axons (Supplementary Table 1). Neurons were simulated individually in NEURON^38^ without synaptic input. Simplified phantom models retained the soma, the longest path in each neurite class and the membrane mechanisms of the retained sections. Their responses were compared with those of the detailed models (Extended Data Fig. 1).

### Neuronal placement and stimulation

Reference phantom simulations placed the soma at the origin, with position varied along the x-axis for spatial steering. In the atlas, detailed neurons were placed in left S1-bf layer 4 and the left CA1 pyramidal layer and oriented according to local anatomy^23, 24^. Placements were shared across instances within each region. Only instances whose morphologies remained within the permitted masks at every tested position and orientation were retained. These masks included neighbouring regions to accommodate cortical and CA1 projections^39, 40, 41^ (Fig. 5b and Supplementary Table 3).

Unless otherwise specified, simulations used 34 °C, a 1 s stimulation plateau and equal peak currents in the two channels. HFS used two 2 kHz carriers. TIS used 2,000 and 2,010 Hz carriers, giving a 10 Hz beat. Total current was the sum of the two channel peak amplitudes. For steering comparisons, the channel-current ratio was varied at matched total current. The unmodulated condition in the phantom current-ratio sweeps used one channel.

### Animals and ethical approval

All animals were maintained under a 12 h light/dark cycle with food and water available ad libitum. For ex vivo brain-slice electrophysiology, male C57BL/6J mice (8–12 weeks old; DBL, Korea) were housed at 22–24 °C and 40–60% relative humidity. All procedures were approved by the Institutional Animal Care and Use Committee (IACUC) of BNH Research (approval no. BnH-2026-1A). The brain-slice dataset comprised recordings from five neurons obtained from five mice, with one neuron per mouse. For ex vivo retinal electrophysiology, male C57BL/6J mice (10 weeks old, 20–25 g; Hana Biotech, Korea) were housed at 22 ± 2 °C and 55 ± 5% relative humidity. All procedures were approved by the IACUC of Pusan National University (approval no. PNU-2025-0663). The retinal dataset comprised recordings from three retinas obtained from three mice, with one retina per mouse.

For in vivo CFA stimulation, male Sprague–Dawley rats (6 weeks old, 200–220 g; Orient Bio, Korea) were housed in pairs in individually ventilated cages at 22 ± 2 °C and 55 ± 5% relative humidity, with lights on at 08:00. All procedures were conducted in accordance with the National Institutes of Health Guide for the Care and Use of Laboratory Animals and approved by the IACUC of Yonsei University (approval no. 2024-0306). Motor-response analyses included five rats.

### Experimental stimulation

An electrically isolated current stimulator based on a mirrored Howland current source^42, 43, 44^ delivered the waveforms. For brain slices and in vivo stimulation, all waveforms were synthesized before delivery through a single channel and the same electrode pair within each experiment. For retinal stimulation, two independently controlled channels each drove a separate electrode pair. Current series and recording exclusions are listed in Supplementary Table 4.

### Brain-slice preparation

Mice were deeply anaesthetized with isoflurane and transcardially perfused with ice-cold sucrose-based artificial cerebrospinal fluid (aCSF) equilibrated with 95% O₂ and 5% CO₂. Brains were rapidly removed, and 400 µm paracoronal slices containing the primary somatosensory barrel cortex were cut at 55° relative to the midline, using a 10° ramp and a vibratome (VT1000 S, Leica Biosystems). Slices recovered in standard aCSF at 35 °C for 30 min and were subsequently maintained at 23–25 °C for 1–4 h before recording.

Standard aCSF contained, in mM, 119 NaCl, 2.5 KCl, 2.5 CaCl₂, 1.3 MgSO₄, 1 NaH₂PO₄, 26.2 NaHCO₃, 11 glucose, 1 sodium pyruvate and 0.4 sodium ascorbate. Sucrose-aCSF contained, in mM, 198 sucrose, 2.5 KCl, 0.5 CaCl₂, 5.0 MgSO₄, 1 NaH₂PO₄, 26.2 NaHCO₃, 11 glucose, 1 sodium pyruvate and 0.4 sodium ascorbate. Recordings were performed at 27–29 °C in aCSF containing NBQX (5 µM), picrotoxin (100 µM) and DL-AP5 (50 µM).

### Whole-cell patch-clamp recording

Whole-cell current-clamp recordings were obtained from layer 4 neurons of the primary somatosensory barrel cortex under infrared differential interference contrast optics. Five neurons were recorded, one per mouse. Patch pipettes had resistances of 3–6 MΩ and contained, in mM, 125 potassium gluconate, 8 NaCl, 10 HEPES, 0.5 EGTA, 4 Mg-ATP and 0.4 Na-GTP, adjusted to pH 7.35 with KOH, with an osmolality of approximately 285 mOsm. Signals were recorded using a MultiClamp 700A amplifier and Digidata 1440 digitizer (Molecular Devices), low-pass filtered at 10 kHz and sampled at 100 kHz using Clampex 10.6.2.2. Input and series resistances were monitored during recording.

### Brain-slice stimulation

Extracellular stimulation was delivered through a pair of Pt/Ir monopolar electrodes (PI3PT30,01B5, Microprobes for Life Science), with one electrode positioned in layers 2/3 approximately 150 µm from the recorded neuron and the other in layer 6 approximately 500 µm away. MI = 0 used an unmodulated 2 kHz sinusoid; MI = 0.5 and 1 were synthesized by summing 2.00 and 2.01 kHz components at amplitude ratios of 3:1 and 1:1, respectively.

Each stimulus comprised a 0.5 s ramp-up, 1 s plateau and 0.5 s ramp-down. MI = 0, 0.5 and 1 were applied in that order at each current, with one 5 s recording sweep per waveform and current at 20 s onset-to-onset intervals. Currents increased in 20 µA steps from 20 to 400 µA for Cells 2–5 and to 800 µA for Cell 1. Intracellular current steps (200 pA, 50 ms) were applied before and after the extracellular series, except for the Cell 1 pre-test (150 pA, 20 ms).

### Brain-slice recording quality assessment

Four Cell 1 conditions lacked valid plateau recordings and were excluded: MI = 0 at 140 and 540 µA, MI = 0.5 at 720 µA, and MI = 1 at 740 µA. Missing responses were not assigned zero AP counts.

Baseline Vm was calculated as the mean raw Vm during the 50 ms before the extracellular-stimulation trigger or intracellular current-step onset. Intracellular current-step responses were analysed without offline filtering. APs were detected as upward 0 mV crossings, retaining the first crossing and subsequent crossings at least 1 ms after the last retained detection, and counted during the first 60 ms after step onset. First-AP amplitude was the maximum raw V_m_ during the 2 ms interval beginning at the first retained crossing minus baseline. Paired pre-test and post-test AP-count and AP-amplitude summaries excluded Cell 1 because its pre-test and post-test steps differed; baseline comparisons included all five cells.

### Retinal preparation

Mice were euthanized by CO₂ inhalation in accordance with institutional guidelines, and the eyes were immediately enucleated. The enucleated eyes were immediately transferred to oxygenated Ames’ medium (A1420, Sigma-Aldrich), and all subsequent retinal dissection procedures were performed under continuous perfusion with oxygenated Ames’ medium. The optic nerve and residual scleral tissue surrounding the optic nerve were removed using spring scissors. The limbus was then incised, and the anterior segment of the eye, including the cornea, was removed. The lens was carefully extracted from the posterior segment using forceps. The sclera and underlying choroid were subsequently removed to isolate the retina. Small peripheral incisions were made using micro-scissors, and the retina was flattened on the multielectrode array (MEA; 60squareMEA200/50iR-Ti, Multi Channel Systems MCS GmbH)^45, 46^.

### Multielectrode-array recording

The isolated retina was then transferred onto a commercial MEA, with the ganglion cell layer facing the electrode array. A fine mesh anchor was placed over the retina to maintain stable contact with the electrodes. After positioning the retina on the electrode array, the bath solution was replaced every 5 min with fresh Ames’ medium containing CNQX (25 μM; Hello Bio) and D-AP5 (25 μM; Hello Bio). Temporal interference stimulation was applied after 20 min of synaptic blocker treatment.

Neural signals were recorded using two 32-channel headstages (RHD2132, Intan Technologies) connected to Omnetics connectors (2432, Omnetics Connector Corporation) integrated into the PCB. The signals were amplified and digitized by the headstages and acquired at a sampling rate of 30 kHz using an RHD 512-channel recording controller (7310, Intan Technologies). The acquisition bandwidth was set to 100–5,000 Hz using the Intan recording software.

### Retinal two-channel stimulation

Four monopolar electrodes (PI3PT30,01B5, Microprobes for Life Science) were configured as two bipolar stimulation pairs, each consisting of two stimulation electrodes. Each electrode pair was secured to an XYZ-axis manipulator and positioned near one of the corners of the electrode array using a USB microscope (M4113ZTL, Dino-Lite). Stimulation waveforms were delivered simultaneously through the two electrode pairs. By varying the current-amplitude ratio between the two electrode pairs, two stimulation conditions were selected to interchange the target and non-target regions. The selected current-amplitude ratio was maintained throughout each stimulation protocol, while the total applied current was increased from low to high intensities. The carrier frequency was fixed at 2 kHz, with a beat frequency of 10 Hz.

### Retinal response analysis

Neural signals recorded from the MEA were analyzed using custom MATLAB scripts (MathWorks). Before spike detection, the recorded signals were digitally band-pass filtered between 100 and 500 Hz to suppress the 2 kHz stimulation artifact. Spikes were detected from the filtered signals using a negative polarity threshold set at three times the standard deviation of the background noise. A refractory period of 1 ms was applied to minimize multiple detections of the same spike. Representative spike waveforms were extracted from the detected spikes, and the mean waveform was calculated for each electrode group.

Neural responses were visualized using raster plots, peristimulus time histograms (PSTHs), and firing-rate heatmaps across the tested stimulation amplitudes. Raster plots and PSTHs were generated from a selected single electrode, with stimulation amplitude represented on the y-axis of the raster plot. Heatmaps were generated from multiple selected electrodes to visualize the firing rate at each stimulation amplitude. PSTHs and heatmaps were calculated using a 20 ms bin width. Neural activity was displayed from −0.5 to 3.0 s relative to the stimulation trigger.

For each stimulation condition, the mean firing rate (FR) of each selected electrode was calculated during the 1 s plateau (0.5–1.5 s after stimulus onset).

### Surgical procedures

Rats were anesthetized with ketamine (100 mg kg⁻¹) and xylazine (5 mg kg⁻¹) and secured in a stereotaxic frame (NARISHIGE). After a midline scalp incision, a cranial window spanning anteroposterior coordinates of 0.0 to 3.0 mm and mediolateral coordinates of 2.0 to 4.0 mm relative to bregma was created. A concentric bipolar electrode (PTM3CC02INS, World Precision Instruments) was then positioned in layer 5 of the caudal forelimb area (CFA, anteroposterior, +1.0 mm; mediolateral, −3.0 mm relative to bregma; dorsoventral, −1.5 mm from the dura mater). A synaptic blocker cocktail comprising CNQX (4.3 mM), D-AP5 (5 mM), and gabazine (0.5 mM) in 0.9% saline was injected locally into the cortical tissue adjacent to the CFA stimulation electrode using a 10 μL Gastight syringe (1701 RN, Hamilton Company) mounted on a Quintessential Stereotaxic Injector (QSI, 53311, Stoelting). Infusions were delivered at a rate of 100 nL min⁻¹. The total volumes injected were 700 nL for Rat 1, 600 nL for Rats 2 and 3, and 500 nL for Rats 4 and 5 (Supplementary Table 4C). The syringe was left in place for 10 min after injection to minimize backflow along the injection tract.

### In vivo cortical stimulation and video recording

Following local injection of synaptic receptor antagonists, cortical stimulation was delivered through the implanted concentric bipolar electrode using a single stimulation channel. HFS used a 2 kHz sinusoid; TIS was synthesized by summing 2.00 and 2.01 kHz components at an amplitude ratio of 1:1.

Each stimulus comprised a 0.5 s ramp-up, a 4 s plateau and a 0.5 s ramp-down. HFS and TIS were applied in that order at each current before increasing the current, with one stimulus per waveform and current at 30 s onset-to-onset intervals. Currents increased in 40 µA steps from 40 to 400 µA for Rat 1, in 20 µA steps from 20 to 400 µA for Rats 2 and 3, and in 20 µA steps from 20 to 300 µA for Rats 4 and 5. HFS was delivered at 360 µA in Rat 2, but the corresponding video was not recorded. This trial was therefore excluded from movement analyses.

Movements of the contralateral left forepaw were recorded from the front using a BFS-U3-19S4M-C camera (FLIR) at 120 frames s⁻¹. An LED remained illuminated throughout each stimulus, including the ramp-up and ramp-down periods.

### Movement tracking and preprocessing

Digit 2 of the left forepaw was tracked using a DeepLabCut 3.0.0^28, 29^ model trained separately for each recording session (Supplementary Table 5A). After five-frame median filtering, coordinates with tracking likelihood below 0.4 were treated as missing. Internal gaps of up to 10 frames were linearly interpolated, and coordinates were converted to millimetres using two reference points 19.70 mm apart in each video (Supplementary Table 5B).

For temporal analyses, four of 159 epochs required additional gap filling. Remaining internal gaps were linearly interpolated and boundary gaps filled with the nearest valid coordinate (Supplementary Table 5C,D). Coordinates were low-pass filtered at 20 Hz and differentiated using each video’s frame rate to obtain velocity.

### Movement events and velocity spectra

For each rat, the first principal component of two-dimensional velocities pooled across currents and waveforms defined a common movement axis. With upward displacement defined as positive, the movement axis was oriented to have a non-negative vertical component. Events were local minima in projected position with a minimum prominence of one image pixel (Supplementary Fig. 10 and Supplementary Table 5E). Plateau events (0.5–4.5 s) were referenced to a 10 Hz clock starting at stimulus onset for both waveforms. PPC was averaged within rat and waveform over matched currents with at least four plateau events in each waveform. Inter-event interval distributions were pooled across currents within each rat and waveform, then averaged with equal weight per rat.

Plateau velocity spectra were estimated using Welch’s method^47^. For display, each epoch’s spectrum was normalized to its maximum over 2–30 Hz. For statistical comparisons, unnormalized spectra at matched currents were used to calculate the mean fraction of 2–30 Hz power within 9–11 Hz and the fraction of epochs with a dominant frequency in that band for each rat and waveform. Peak displacement and path length were measured over the full train, with recording-specific exclusions detailed in Extended Data Fig. 8.

### Membrane-potential filtering and AP detection

Carrier-comb filtering attenuated carrier harmonics from CA1 simulations and whole-cell recordings. Cortical simulations used adaptive-comb filtering to attenuate additional trace-specific carrier subharmonics. APs were detected as upward 0 mV crossings in filtered membrane potential, with a minimum interval of 1 ms between detections. The two filters and a band-stop/high-pass procedure adapted from Grossman et al.^8^ were compared on the same recordings and simulated traces (Supplementary Figs. 11 and 12).

### Activation thresholds, firing rates and TIS operating windows

Activation required at least one plateau AP. Firing rate was the plateau AP count divided by plateau duration. Relative threshold was the ratio of TIS to HFS activation currents under matched conditions. A TIS operating window comprised tested suprathreshold currents for which HFS elicited no plateau APs and TIS elicited at least one AP in every beat period. Normalized rates were expressed relative to the maximum for each cell or model across all tested currents and waveforms.

### Spatial recruitment and selectivity

In the phantom, target positions had somatic MI ≥ 0.9. Target length was the summed extent of intervals meeting this criterion along the x-axis (Extended Data Fig. 4 and Supplementary Fig. 5). In the atlas, CA1 positions meeting the same criterion formed the target. All remaining sampled CA1 positions and all S1-bf layer 4 placements were off-target. Activation fractions were calculated within each group. Atlas configurations were weighted by the volume of the corresponding layer divided by the total number of model-instance, position and orientation combinations in that layer (Supplementary Table 2C). The selectivity index was the difference between target and off-target activation fractions divided by their sum and was undefined when both fractions were zero.

### Beat phase and pairwise phase consistency

For 10 Hz simulations and whole-cell recordings, plateau AP phases were referenced to a 10 Hz clock starting at plateau onset for all waveforms. For modulated waveforms, 0° and 180° corresponded approximately to the envelope minimum and maximum. Pairwise phase consistency (PPC) was the mean cosine of the pairwise phase differences for sweeps or simulations containing at least four plateau APs^25^. The same estimator was used for movement events, with phases referenced to stimulation onset. In the circular plots, the arrow indicates the mean event phase and its length the mean resultant length (Supplementary Fig. 10d).

### Membrane-potential window analysis

Voltage windows were selected from filtered traces, and raw membrane potentials were averaged after excluding APs and transitions (Supplementary Fig. 6). Each TIS beat or HFS plateau was classified as two-level when the fitted high level exceeded the fitted low level by at least 10 mV and the two-level residual cost was at most 80% of the constant-fit cost. Other segments were classified as single-level. Low-window and single-level samples were pooled where indicated. For the slow-envelope simulation, the same comparison was applied to a full 20 s segment to estimate suppression onset and release.

### Somatic and axonal AP propagation analysis

For axonal rasters, detections were linked across nearby recording sites and coloured by the site of the earliest observed upward crossing in each group. Intervening subthreshold deflections could support a link but did not count as APs. The first detected site need not be the site of AP initiation (Extended Data Fig. 10).

### Statistical analysis

Deterministic simulations were summarized descriptively across model instances, placements and orientations. Brain-slice TOW and pre-test/post-test measures were also summarized descriptively. In Fig. 6d, each cell contributed one mean per waveform over TOW currents with valid recordings for all three waveforms.

For retinal recordings, statistical comparisons between the TIS and HFS conditions were performed using a two-sided Welch’s two-sample t-test. Electrode-level mean firing rate (FR) values were treated as individual observations for statistical analysis. The significance level was set at p < 0.05. Data are presented as mean ± s.e.m., with individual electrode measurements shown where applicable.

Rat-level HFS and TIS outcomes were compared using two-sided paired t-tests (n = 5 rats). Holm correction^48^ was applied jointly to movement-event PPC (Fig. 8b), the 9–11 Hz power fraction and the fraction of epochs with a dominant frequency in that band (Extended Data Fig. 9b). Confidence intervals for paired mean differences used the t distribution with four degrees of freedom and were unadjusted for multiplicity. Rat-level summaries are shown as mean ± s.d.

## Data availability

The simulation outputs, electrophysiological data and forelimb-tracking data supporting this study will be made publicly available on GitHub upon journal publication.

## Code availability

The custom code used for the simulations and data analyses will be made publicly available on GitHub upon journal publication.

## Acknowledgements

The authors thank members of the Neural Interface Lab for their valuable feedback and Medical Illustration & Design (MID), part of the Medical Research Support Services at Yonsei University College of Medicine, for their support with medical illustrations.

## Author contributions

K.E. and J.M. conceived and designed the study. J.M. developed and validated the software. J.M. and J.C. developed the stimulation hardware. J.-H.J., J.M. and K.E. performed the whole-cell patch-clamp experiments. Y.P. and J.M. conducted the ex vivo retinal experiments. J.M., C.S.K., S.H.M., H.R.O. and H.J.L. conducted the in vivo cortical experiments. J.M., K.E. and Y.P. analysed the data and prepared the figures. S.-Y.P., B.T.C., H.H.J., S.C. and K.E. supervised the study. K.E., S.C., H.H.J. and C.S.K. acquired funding. K.E., J.M., Y.P., J.-H.J., S.H.M. and C.S.K. wrote the original manuscript draft. All authors reviewed the manuscript.

---

**Extended Data Fig. 1.**
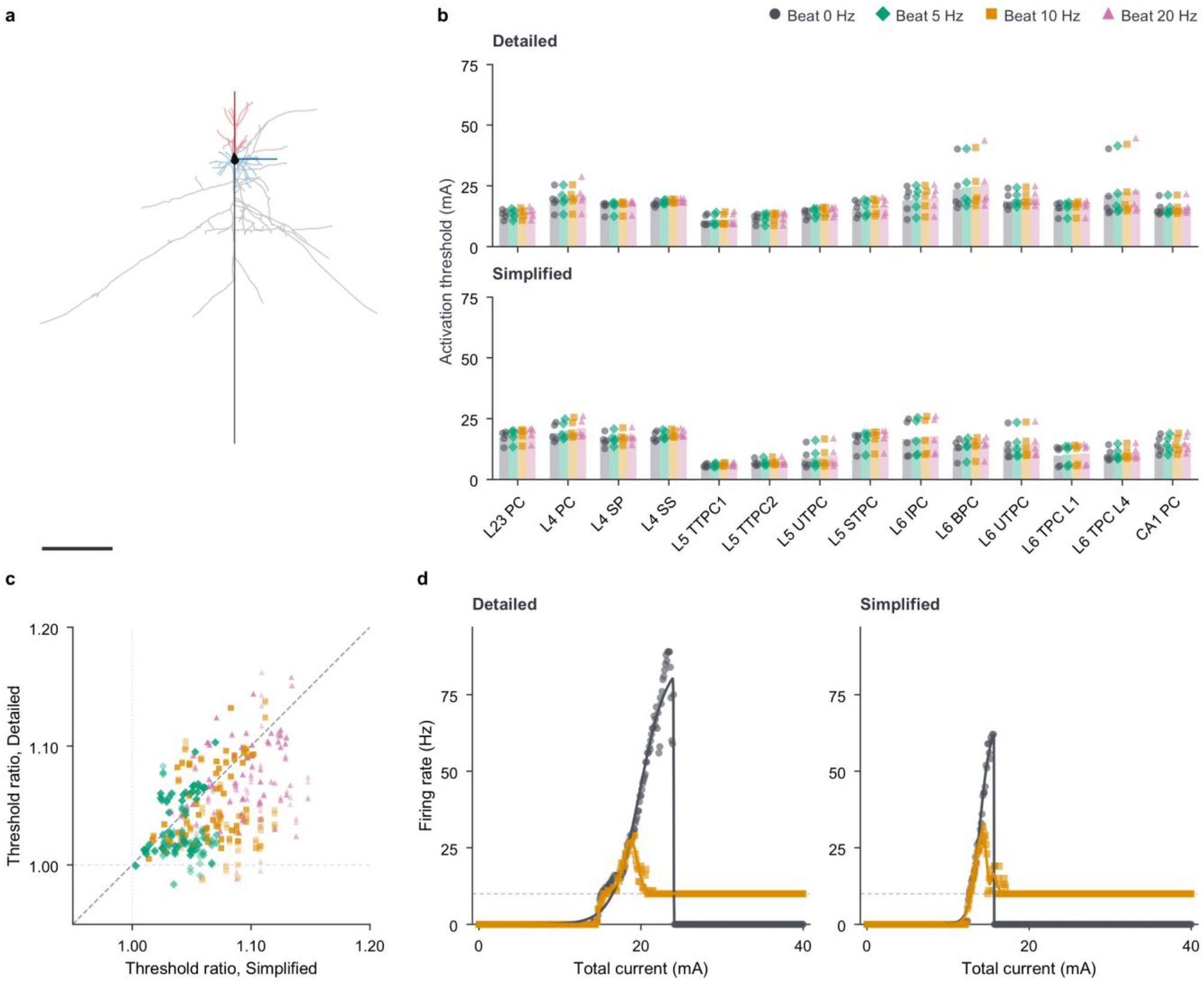
| Effects of morphological simplification on activation thresholds and firing rate. Detailed and simplified neuronal models were compared in a homogeneous phantom. **a**, Example L2/3 PC morphology: detailed morphology in pale colours and retained, straightened paths in darker colours. Scale bar, 500 µm; neurite widths and soma size are enlarged for visibility. **b**, Absolute activation thresholds for the 13 excitatory cortical model types and the CA1 PC model (Supplementary Table 1), under HFS and under TIS at beat frequencies of 5, 10 and 20 Hz. Upper, detailed models; lower, simplified models. Symbols, each clone’s median threshold across four axial rotations; bars, means of the five clone medians. **c**, Matched TIS-to-HFS activation-threshold ratios for simplified and detailed morphologies, with one point per clone, rotation and beat frequency. Horizontal and vertical reference lines mark ratio 1; the diagonal marks equality between morphologies. Some detailed-model ratios fall below 1 (minimum displayed ratio, 0.983). **d**, Somatic firing rates for CA1 PC clone 1 at 270° axial rotation, comparing HFS with TIS with a 10 Hz beat. Symbols, simulation results; dashed line, 10 Hz. All simulations used unmyelinated models at the phantom origin, 34 °C, a 2 kHz carrier and equal channel amplitudes.

**Extended Data Fig. 2.**
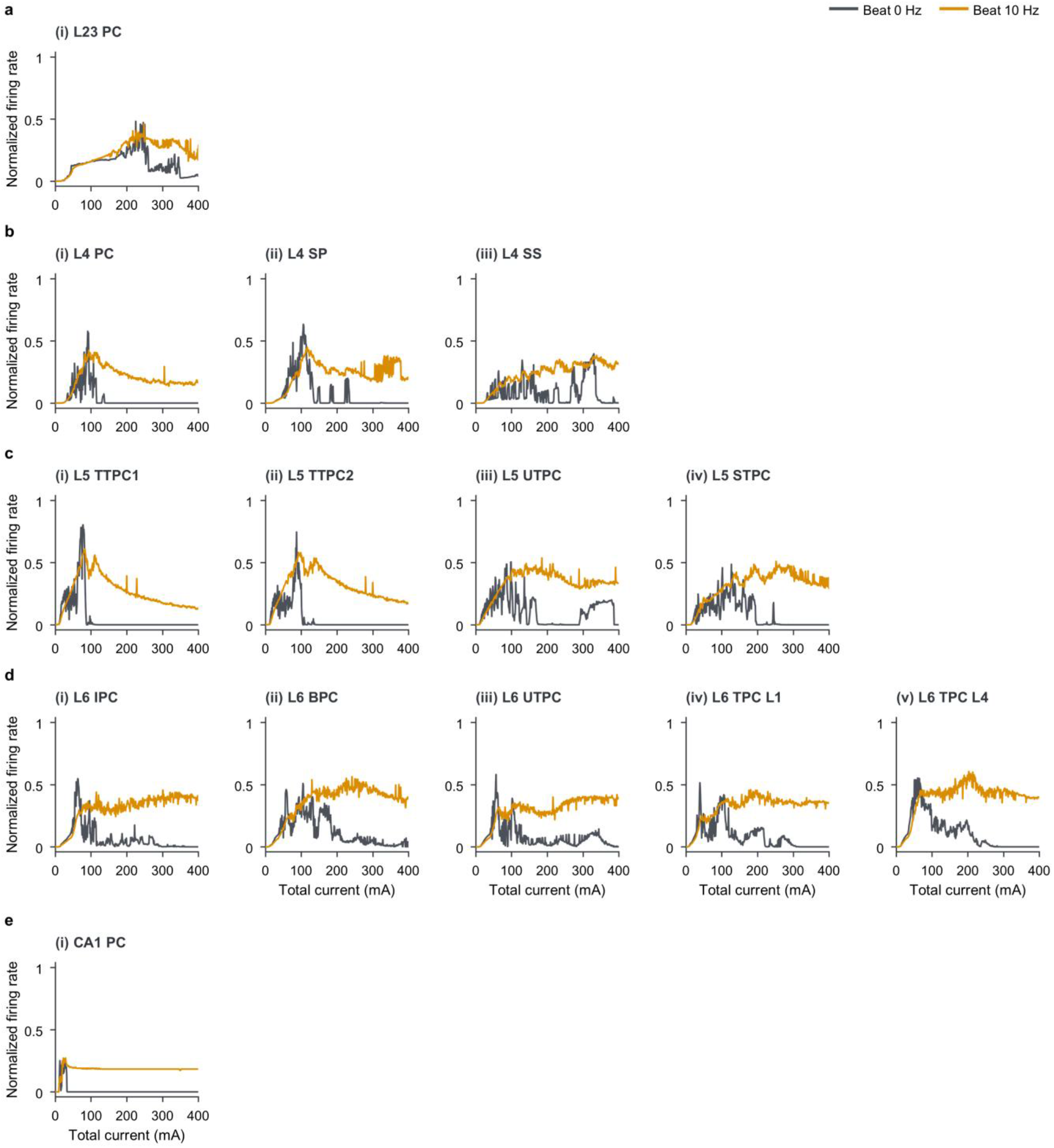
| Normalized somatic firing rates across model types under high-frequency and temporal interference stimulation. Normalized firing rates for 14 simulated model types. Total current was swept from 0 to 400 mA. Somatic APs were detected from filtered Vm and counted during the 1 s plateau using model-specific procedures (Methods; filtering comparisons in Supplementary Figs. 11 and 12). Each clone’s firing rates were normalized by its maximum across both waveforms and all tested currents; lines show the mean of the five normalized clone responses for each model type and waveform at each current. **a–d**, The 13 excitatory cortical model types grouped by layer: L2/3, L4, L5 and L6, respectively. **e**, CA1 PC model. All simulations used morphologically simplified, unmyelinated models at the phantom origin, 34 °C, a 2 kHz carrier and equal channel amplitudes, with a 10 Hz beat for TIS.

**Extended Data Fig. 3.**
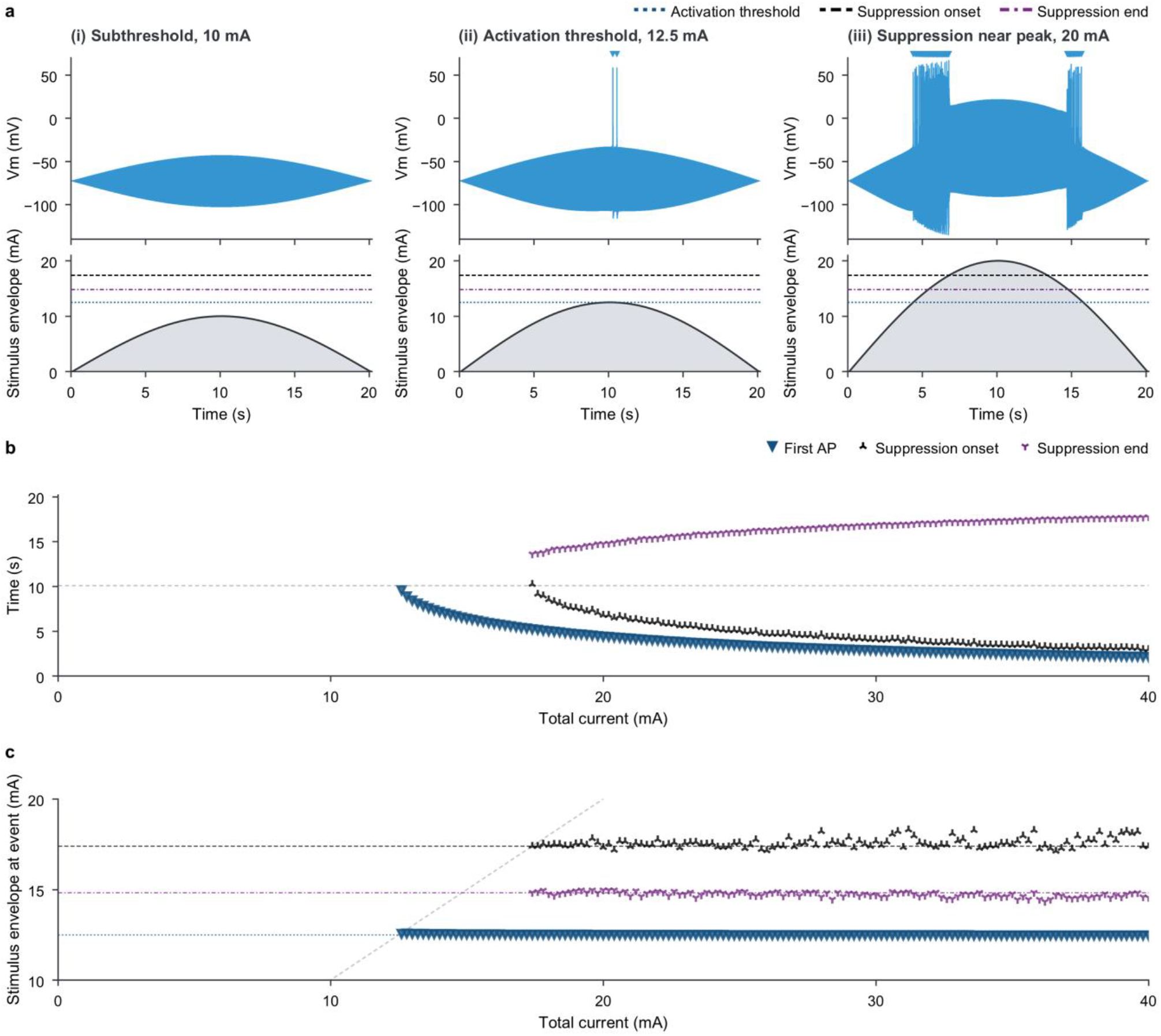
| Firing onset and suppression boundaries occur at nearly constant envelope amplitudes during slowly modulated stimulation. **a**, One envelope period at (i) 10, (ii) 12.5 and (iii) 20 mA total current. Top, raw somatic Vm; triangles, APs detected from filtered Vm. Bottom, stimulus envelope. At 20 mA, firing on the rising and falling portions is separated by an AP-free interval near the envelope peak. Suppression boundaries were identified from filtered Vm using the window-selection procedure (Methods and Supplementary Fig. 6). Dark-blue dotted line, activation threshold, 12.5 mA. Black dashed and purple dash-dot lines, envelope amplitudes at suppression onset and end, 17.4 and 14.8 mA, respectively, obtained from the 17.4 mA simulation, the lowest tested current showing suppression. **b,c**, Times of the first AP and of suppression onset and end (b), and the corresponding envelope amplitudes (c), across the 0–40 mA sweep. Grey dashed lines mark the envelope peak: 10.1 s in b and total current in c. All simulations used the morphologically simplified, unmyelinated CA1 model at the phantom origin, 34 °C, equal channel amplitudes and carrier frequencies of 2,000 and 2,000.05 Hz. Each stimulus spanned one 20 s envelope period, rising to its maximum over 10 s and falling over the next 10 s, without additional onset or offset ramps.

**Extended Data Fig. 4.**
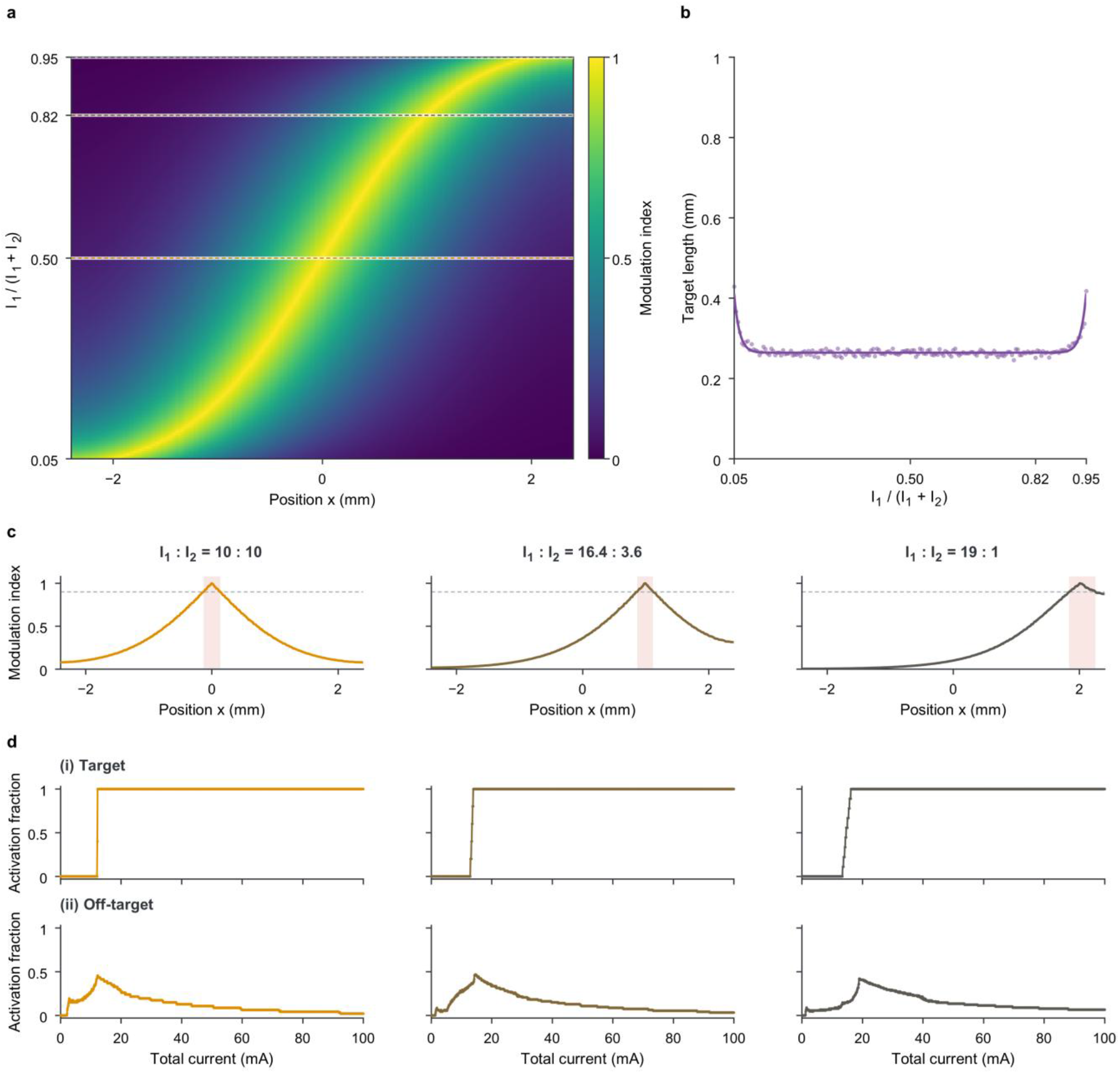
| Modulation index–defined target geometry and activation fractions underlying spatial selectivity. In **c** and **d**, columns show CH1:CH2 current ratios of 10:10, 16.4:3.6 and 19:1. **a**, MI along y = z = 0 as a function of the CH1 current fraction, I₁/(I₁ + I₂), where I₁ and I₂ are the peak current amplitudes in CH1 and CH2, respectively. Dashed lines mark the three ratios used in Fig. 3. **b**, Summed length of regions satisfying MI ≥ 0.9 along y = z = 0; symbols, calculated lengths; line, fitted guide. **c**, MI profiles along y = z = 0. Red shading, the target; dashed line, MI = 0.9. **d**, Activation fractions of sampled positions within (i) the target and (ii) the off-target region versus total current. Total current was matched across ratios at each tested amplitude. These activation fractions were used to calculate SI in Fig. 3d. Neuronal responses were sampled at x-coordinates from −2.4 to 2.4 mm in 0.05 mm steps (y = z = 0). Curve colours in c and d identify current ratios.

**Extended Data Fig. 5.**
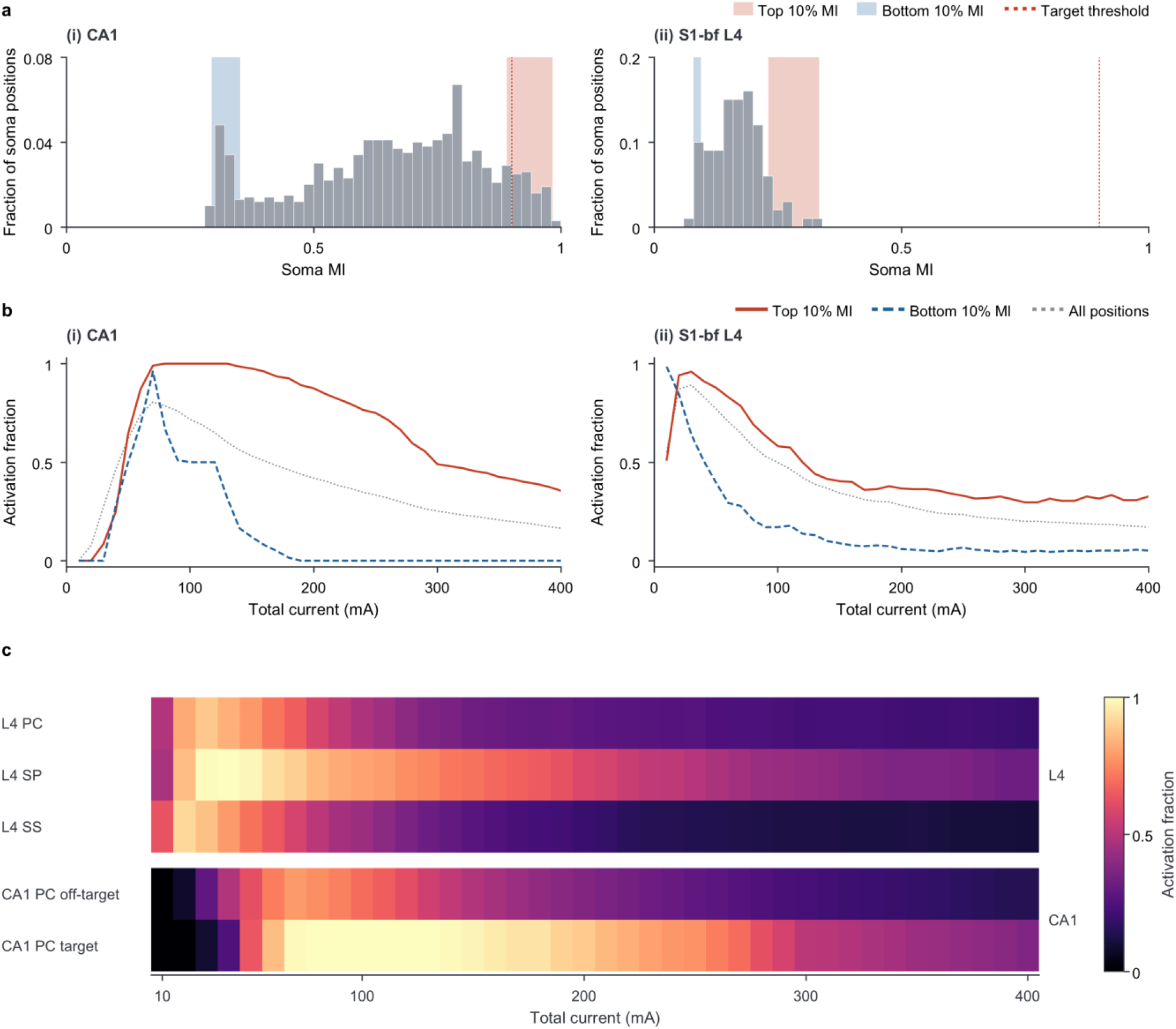
| Neuronal recruitment by somatic modulation index and cell type in the atlas-based rat-brain model. **a**, Somatic MI distributions across unique positions in (i) CA1 and (ii) S1-bf layer 4. Red and blue bands, MI ranges of the top and bottom 10% of positions ranked by somatic MI within each region, respectively; dotted line, the target MI threshold used in Fig. 5 (MI = 0.9). **b**, Activation fractions versus total current for (i) CA1 and (ii) S1-bf layer 4, comparing the top and bottom 10% of positions ranked by somatic MI with all positions. Positions were ranked separately within each region, and all clones and orientations at the selected positions were included. Red solid and blue dashed lines, upper and lower MI deciles, respectively; grey dotted lines, all positions. **c**, Activation fractions versus total current for each L4 cell type, with CA1 target and off-target placements shown separately. Clones were pooled within each row: four L4 PC, one L4 SP, four L4 SS and two CA1 PC clones (Supplementary Table 3). Each CA1 clone occupied the same 1,000 positions; each L4 clone occupied the same 100 positions with three orientations per position. Fractions in b and c were calculated within each displayed group; regional volume weighting was applied when combining off-target groups in Fig. 5d. All simulations used TIS with carrier frequencies of 2,000 and 2,010 Hz and equal channel amplitudes. Total current ranged from 10 to 400 mA.

**Extended Data Fig. 6.**
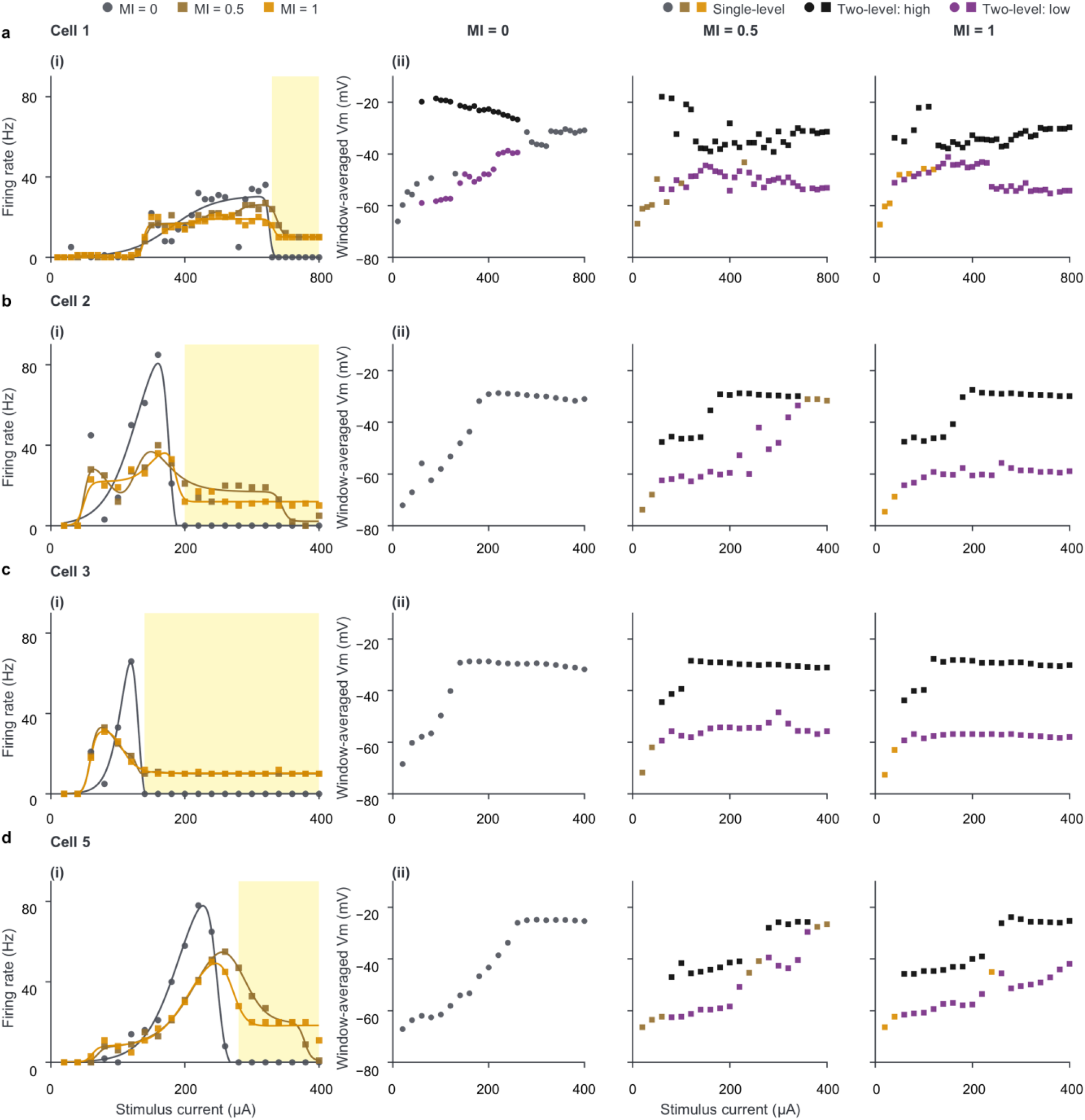
| Firing rate and membrane potential in the four additional cortical neurons. Whole-cell recordings from the four remaining neurons in the Fig. 6 dataset: **a–d**, Cells 1, 2, 3 and 5, respectively (n = 4 cells, one per mouse). **(i)**, Plateau firing rate versus current. Symbols, individual sweeps; lines, fitted guides; yellow shading, TOW. Complete plateau recordings were unavailable for four conditions in Cell 1, and these conditions were excluded: MI = 0 at 140 and 540 µA, MI = 0.5 at 720 µA, and MI = 1 at 740 µA. Shading spans the MI = 1 condition at 740 µA, for which the TOW criterion could not be evaluated. **(ii)**, Window-averaged raw Vm for the same sweeps, with columns corresponding to MI = 0, 0.5 and 1. Waveform-coloured symbols, single-level means in sweeps without two-level segments; black symbols, high-window means; purple symbols, pooled single-level and low-window means in sweeps containing two-level segments. Stimulation and analysis were as in Fig. 6. Tested currents ranged from 20 to 400 µA for Cells 2, 3 and 5 and from 20 to 800 µA for Cell 1.

**Extended Data Fig. 7.**
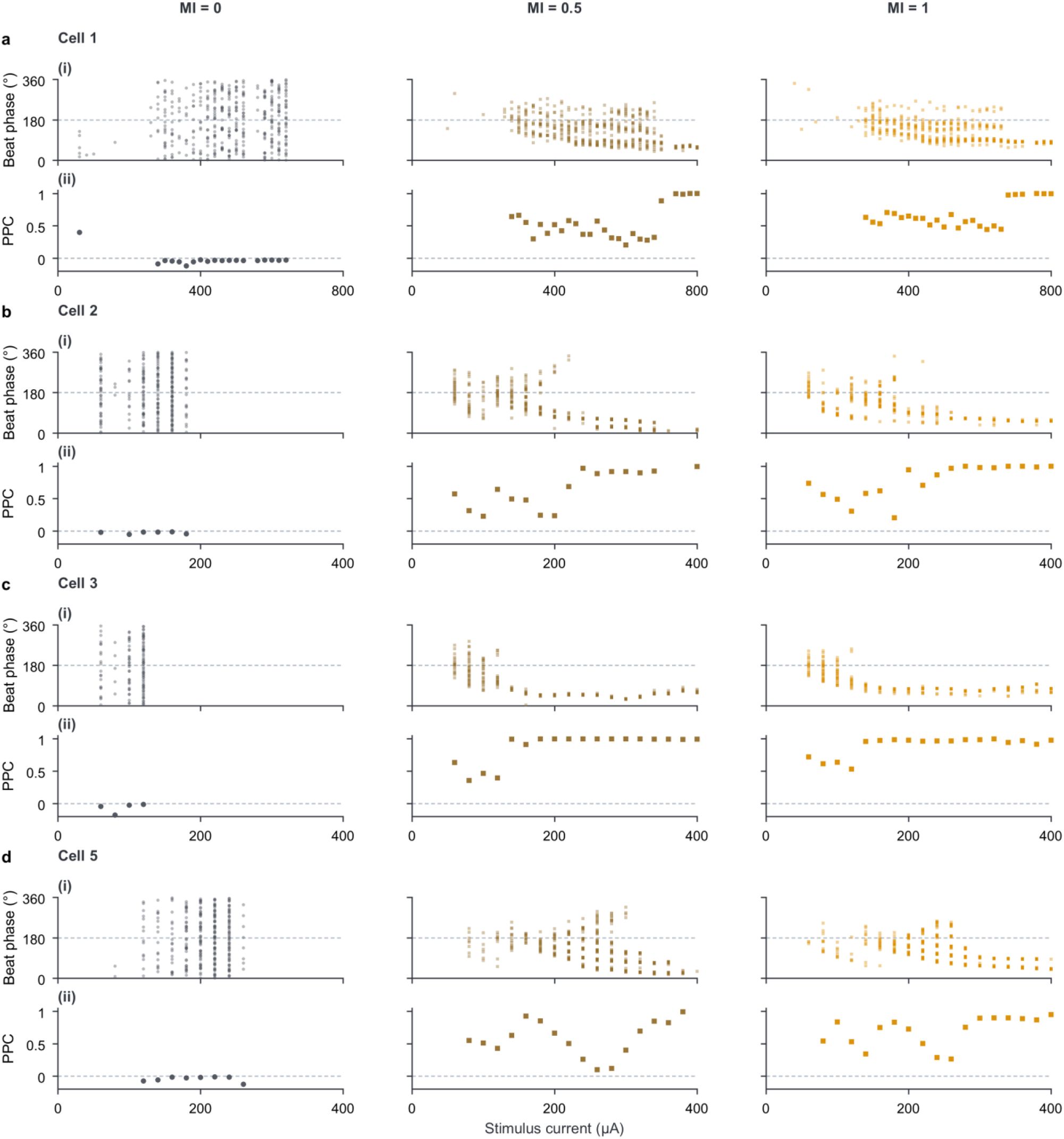
| Action-potential phase and phase consistency in four additional cortical neurons. Whole-cell recordings from the four remaining neurons in the Fig. 6 dataset: **a–d**, Cells 1, 2, 3 and 5, respectively (n = 4 cells, one per mouse); columns, MI = 0, 0.5 and 1. **(i)**, Phase of individual plateau APs, referenced to a 10 Hz clock starting at plateau onset for all three waveforms; dashed line, 180°. **(ii)**, PPC per sweep containing at least four plateau APs; dashed line, zero. Stimulation and analysis were as in Fig. 6. Tested currents ranged from 20 to 400 µA for Cells 2, 3 and 5 and from 20 to 800 µA for Cell 1.

**Extended Data Fig. 8.**
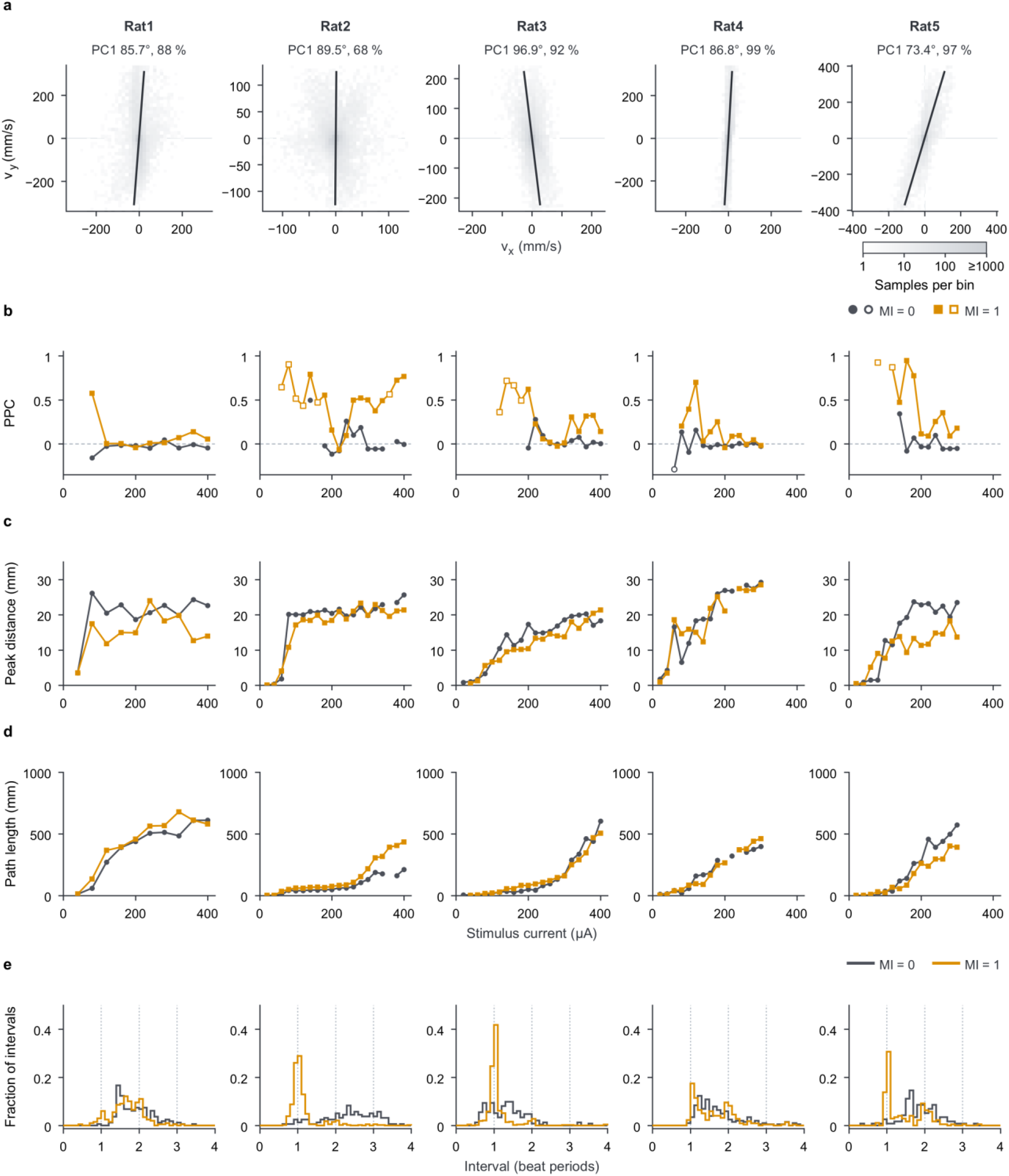
| Forelimb movement responses across stimulation currents in individual rats. Columns, Rats 1–5 from Fig. 8. **a**, Two-dimensional histograms of velocity samples pooled across waveforms and currents within each rat. Display ranges contain 99% of each rat’s samples. Lines, first principal component (PC1). Labels give the axis angle and percentage of variance explained. **b**, PPC per epoch with ≥4 plateau events, referenced to a 10 Hz clock from stimulation onset. Filled symbols, epochs included in Fig. 8b animal means (≥4 plateau events for each waveform at the same current). Open symbols, other eligible epochs. Dashed line, zero. **c**, Peak displacement of digit 2 from its pre-onset position over 0–5 s. Three epochs were omitted because the reference position was unavailable or the peak occurred in a gap-filled frame. **d**, Total path length of digit 2 over 0–5 s. Four epochs containing gap-filled frames were omitted. In b–d, symbols represent individual stimulation epochs and lines connect neighbouring tested currents with valid values. **e**, Step histograms of plateau inter-event intervals pooled across currents for each rat and waveform, expressed in units of the 10 Hz reference period. Fractions include all intervals, including those beyond the plotted range. Vertical dotted lines, one, two and three periods. Stimulation was as in Fig. 8. Analyses in b and e used the 0.5–4.5 s plateau.

**Extended Data Fig. 9.**
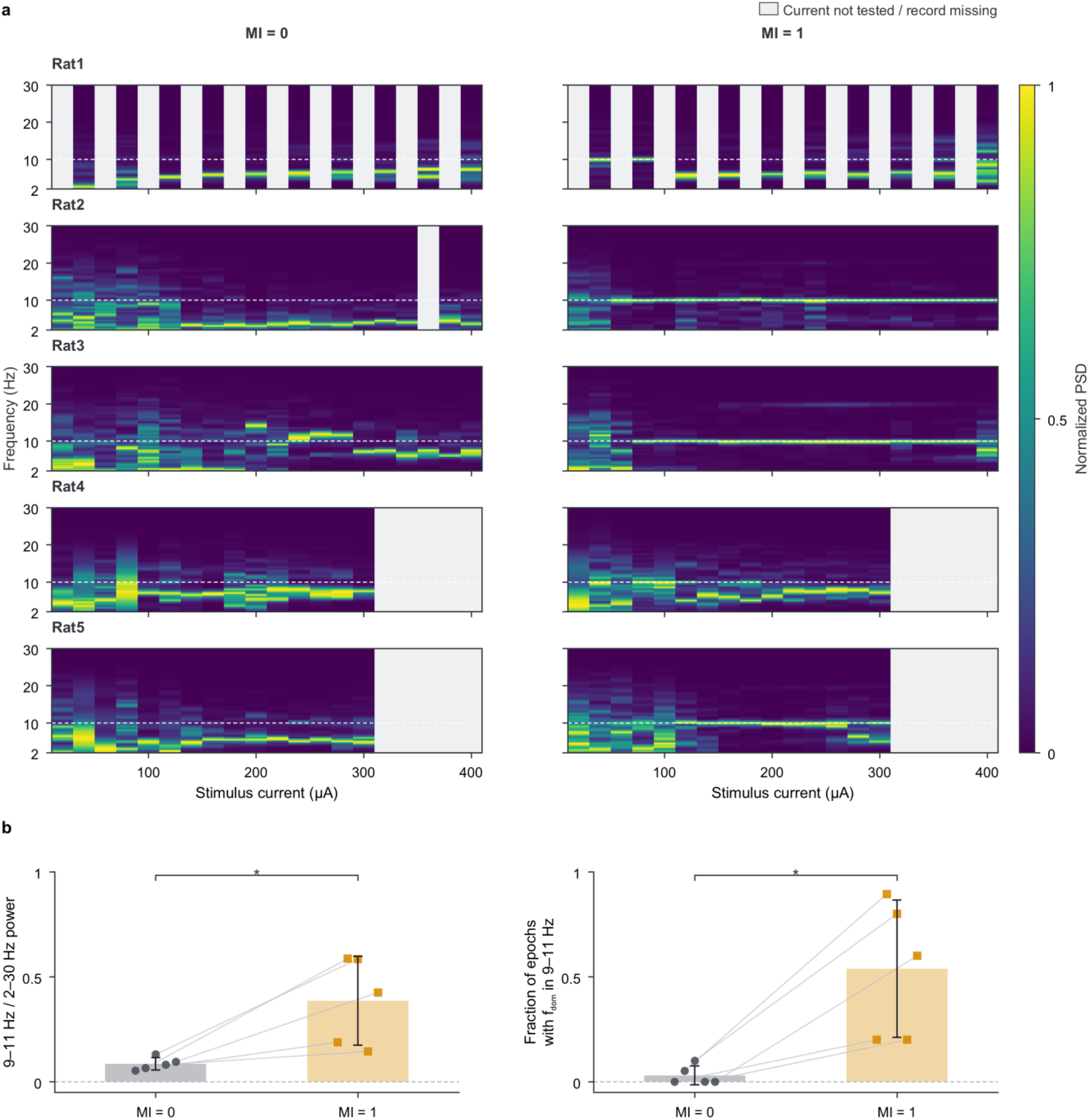
| Forelimb velocity spectra and beat-frequency measures in individual rats. Forelimb movement analyses for the five rats in Fig. 8. **a**, PSD of movement-axis velocity versus stimulus current. Rows, Rats 1–5; left and right columns, MI = 0 and 1, respectively. Each epoch’s PSD was normalized to its maximum over 2–30 Hz. Dashed lines, 10 Hz; light-grey columns, currents not tested or without a recording. **b**, Spectral measures calculated using matched currents with valid spectra for both waveforms. Left, fraction of 2–30 Hz power within 9–11 Hz, averaged across currents within each animal. Right, fraction of epochs with a dominant frequency within 9–11 Hz. The dominant frequency was defined as the frequency of the maximum PSD within 2–30 Hz. Symbols and connecting lines, individual animals; bars and error bars, mean ± s.d. (n = 5 rats); dashed lines, zero. Two-sided paired t-tests with Holm correction across these two comparisons and the PPC comparison in Fig. 8b; adjusted *p* = 0.040 for both comparisons; asterisks, *p* < 0.05. Stimulation was as in Fig. 8. All analyses used the 0.5–4.5 s plateau.

**Extended Data Fig. 10.**
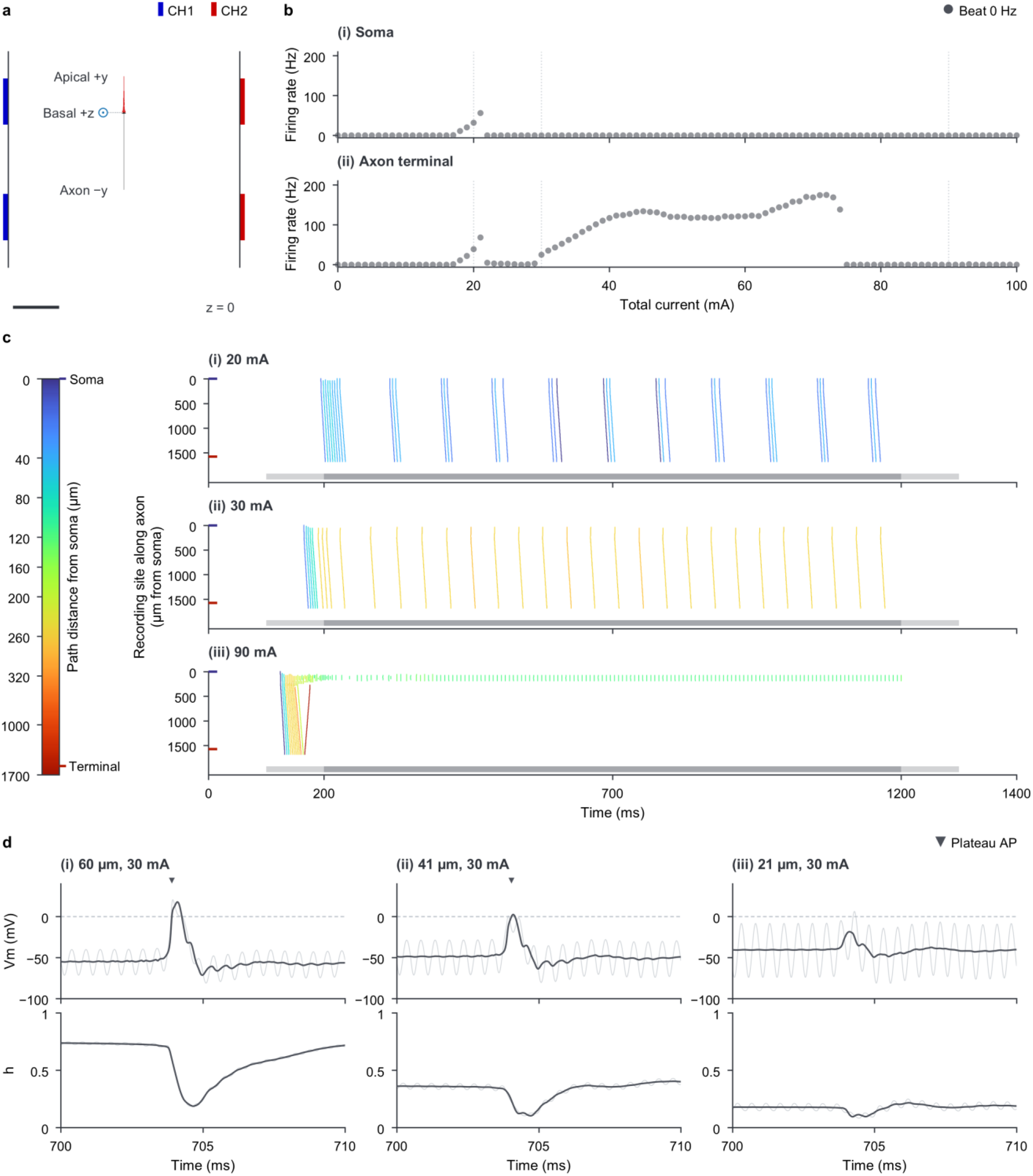
| Axonal activity can persist despite somatic firing suppression under high-frequency stimulation. **a**, Placement and morphology projected onto the z = 0 plane; the basal dendrite points out of the plane. Scale bar, 1 mm. **b**, Plateau firing rate versus total current at (i) the soma and (ii) the central segment of the terminal axonal section, at a path distance of 1,576 µm from the soma. Symbols, simulation results; vertical dotted lines, currents shown in c. **c**, AP-detection rasters from filtered Vm at (i) 20, (ii) 30 and (iii) 90 mA, sampled at the soma and 85 axonal sites. Vertical position, path distance from the soma. Colours, path distance of the site with the earliest upward 0 mV crossing in each linked group of detections (Methods); the colour scale is expanded for proximal distances. Coloured axis ticks, the soma and the central segment of the terminal axonal section used in b. Horizontal bars, stimulation periods; lighter segments, ramps. **d**, Vm (top) and Na⁺-channel inactivation gate *h* (bottom) at axonal sites (i) 60, (ii) 41 and (iii) 21 µm from the soma during 700–710 ms of the 30 mA simulation. Light and dark grey traces, raw and filtered signals, respectively; triangles, detected APs; dashed lines, 0 mV. All simulations used the model and HFS settings of Fig. 2a, with the soma positioned at (x, y, z) = (0, 1, 0) mm in the phantom.

## References

1. Wagner, T., Valero-Cabre, A. & Pascual-Leone, A. Noninvasive Human Brain Stimulation. Annu Rev Biomed Eng 9, 527–565 (2007).

2. Polanía, R., Nitsche, M. A. & Ruff, C. C. Studying and modifying brain function with non-invasive brain stimulation. Nat Neurosci 21, 174–187 (2018).

3. Eom, K. & Byun, K. M. Advanced non-invasive brain neuromodulation: a comprehensive review of electric, magnetic, ultrasonic, and optical modalities. Biomed Eng Lett 16, 1095–1116 (2026).

4. Datta, A. et al. Gyri-precise head model of transcranial direct current stimulation: Improved spatial focality using a ring electrode versus conventional rectangular pad. Brain Stimul 2, 201–207.e1 (2009).

5. Opitz, A., Paulus, W., Will, S., Antunes, A. & Thielscher, A. Determinants of the electric field during transcranial direct current stimulation. Neuroimage 109, 140–150 (2015).

6. Edwards, D. et al. Physiological and modeling evidence for focal transcranial electrical brain stimulation in humans: A basis for high-definition tDCS. Neuroimage 74, 266–275 (2013).

7. Kuo, H. I. et al. Comparing Cortical Plasticity Induced by Conventional and High-Definition 4 × 1 Ring tDCS: A Neurophysiological Study. Brain Stimul 6, 644–648 (2013).

8. Grossman, N. et al. Noninvasive Deep Brain Stimulation via Temporally Interfering Electric Fields. Cell 169, 1029–1041.e16 (2017).

9. Rampersad, S. et al. Prospects for transcranial temporal interference stimulation in humans: A computational study. Neuroimage 202, 116124 (2019).

10. Lee, S., Lee, C., Park, J. & Im, C. H. Individually customized transcranial temporal interference stimulation for focused modulation of deep brain structures: a simulation study with different head models. Sci Rep 10, 11730 (2020).

11. Song, X., Zhao, X., Li, X., Liu, S. & Ming, D. Multi-channel transcranial temporally interfering stimulation (tTIS): application to living mice brain. J Neural Eng 18, 036003 (2021).

12. Botzanowski, B. et al. Focal control of non-invasive deep brain stimulation using multipolar temporal interference. Bioelectron Med 11, 7 (2025).

13. Savvateev, I. et al. Multipair phase-modulated temporal interference electrical stimulation combined with fMRI. Cell Syst 17, 101610 (2026).

14. Violante, I. R. et al. Non-invasive temporal interference electrical stimulation of the human hippocampus. Nat Neurosci 26, 1994–2004 (2023).

15. Missey, F. et al. Non-invasive temporal interference stimulation of the hippocampus suppresses epileptic biomarkers in patients with Epilepsy: biophysical differences between kilohertz and amplitude modulated stimulation. Brain Stimul 19, 102981 (2026).

16. Yang, C. et al. Transcranial temporal interference stimulation targeting the subthalamic region for motor symptoms in Parkinson’s disease: a pilot, randomised, double-blind, sham-controlled crossover study. EBioMedicine 127, 106245 (2026).

17. Huang, Y., Datta, A. & Parra, L. C. Optimization of interferential stimulation of the human brain with electrode arrays. J Neural Eng 17, 036023 (2020).

18. Mirzakhalili, E., Barra, B., Capogrosso, M. & Lempka, S. F. Biophysics of Temporal Interference Stimulation. Cell Syst 11, 557–572.e5 (2020).

19. Wang, B., Aberra, A. S., Grill, W. M. & Peterchev, A. V. Responses of model cortical neurons to temporal interference stimulation and related transcranial alternating current stimulation modalities. J Neural Eng 19, 066047 (2023).

20. Opančar, A. et al. The same biophysical mechanism is involved in both temporal interference and direct kHz stimulation of peripheral nerves. Nat Commun 16, 9006 (2025).

21. Caldas-Martinez, S. et al. Cell-specific effects of temporal interference stimulation on cortical function. Commun Biol 7, 1076 (2024).

22. Migliore, R. et al. The physiological variability of channel density in hippocampal CA1 pyramidal cells and interneurons explored using a unified data-driven modeling workflow. PLoS Comput Biol 14, e1006423 (2018).

23. Romani, A. et al. Community-based reconstruction and simulation of a full-scale model of the rat hippocampus CA1 region. PLoS Biol 22, e3002861 (2024).

24. Markram, H. et al. Reconstruction and Simulation of Neocortical Microcircuitry. Cell 163, 456–492 (2015).

25. Vinck, M., van Wingerden, M., Womelsdorf, T., Fries, P. & Pennartz, C. M. A. The pairwise phase consistency: A bias-free measure of rhythmic neuronal synchronization. Neuroimage 51, 112–122 (2010).

26. Lee, J. I., Werginz, P., Kameneva, T., Im, M. & Fried, S. I. Membrane depolarization mediates both the inhibition of neural activity and cell-type-differences in response to high-frequency stimulation. Commun Biol 7, 734 (2024).

27. Kleven, H. et al. Waxholm Space atlas of the rat brain: a 3D atlas supporting data analysis and integration. Nat Methods 20, 1822–1829 (2023).

28. Mathis, A. et al. DeepLabCut: markerless pose estimation of user-defined body parts with deep learning. Nat Neurosci 21, 1281–1289 (2018).

29. Nath, T. et al. Using DeepLabCut for 3D markerless pose estimation across species and behaviors. Nat Protoc 14, 2152–2176 (2019).

30. Bhadra, N., Lahowetz, E. A., Foldes, S. T. & Kilgore, K. L. Simulation of high-frequency sinusoidal electrical block of mammalian myelinated axons. J Comput Neurosci 22, 313–326 (2007).

31. Fisher, K. M., Jillani, N. E., Oluoch, G. O. & Baker, S. N. Blocking central pathways in the primate motor system using high-frequency sinusoidal current. J Neurophysiol 113, 1670–1680 (2015).

32. Vieira, P. G., Krause, M. R. & Pack, C. C. Temporal interference stimulation disrupts spike timing in the primate brain. Nat Commun 15, 4558 (2024).

33. Saturnino, G. B. et al. SimNIBS 2.1: A Comprehensive Pipeline for Individualized Electric Field Modelling for Transcranial Brain Stimulation. In Brain and Human Body Modeling: Computational Human Modeling at EMBC 2018 (eds Makarov, S., Horner, M. & Noetscher, G.) 3–25 (Springer International Publishing, Cham, 2019).

34. Puonti, O. et al. Accurate and robust whole-head segmentation from magnetic resonance images for individualized head modeling. Neuroimage 219, 117044 (2020).

35. D’Alessandro, S., Handler, M., Saba, R., Garnham, C. & Baumgarten, D. Computer Simulation of the Electrical Stimulation of the Human Vestibular System: Effects of the Reactive Component of Impedance on Voltage Waveform and Nerve Selectivity. J Assoc Res Otolaryngol 23, 815–833 (2022).

36. Jackson, M. P. et al. Safety parameter considerations of anodal transcranial Direct Current Stimulation in rats. Brain Behav Immun 64, 152–161 (2017).

37. Karimi, N., Amirfattahi, R. & Zeidaabadi Nezhad, A. Neuromodulation effect of temporal interference stimulation based on network computational model. Front Hum Neurosci 18, 1436205 (2024).

38. Hines, M. L. & Carnevale, N. T. Neuron: A Tool for Neuroscientists. Neuroscientist 7, 123–135 (2001).

39. Staiger, J. F. et al. Functional Diversity of Layer IV Spiny Neurons in Rat Somatosensory Cortex: Quantitative Morphology of Electrophysiologically Characterized and Biocytin Labeled Cells. Cereb Cortex 14, 690–701 (2004).

40. Narayanan, R. T. et al. Beyond Columnar Organization: Cell Type- and Target Layer-Specific Principles of Horizontal Axon Projection Patterns in Rat Vibrissal Cortex. Cereb Cortex 25, 4450–4468 (2015).

41. Cenquizca, L. A. & Swanson, L. W. Spatial organization of direct hippocampal field CA1 axonal projections to the rest of the cerebral cortex. Brain Res Rev 56, 1–26 (2007).

42. Bertemes-Filho, P., Vincence, V. C., Santos, M. M. & Zanatta, I. X. Low power current sources for bioimpedance measurements: a comparison between Howland and OTA-based CMOS circuits. Journal of Electrical Bioimpedance 3, 66–73 (2012).

43. Grando Sirtoli, V., Coelho Vincence, V. & Bertemes-Filho, P. Mirrored enhanced Howland current source with feedback control. Rev Sci Instrum 90, 024702 (2019).

44. FallahRad, M. et al. Electrophysiology equipment for reliable study of kHz electrical stimulation. J Physiol 597, 2131–2137 (2019).

45. Lee, D. H. et al. Microelectrothermoforming (μETF): one-step versatile 3D shaping of flexible microelectronics for enhanced neural interfaces. npj Flex Electron 9, 5 (2025).

46. Park, Y. et al. Focused Ultrasound as a Novel Non-Invasive Method for the Delivery of Gold Nanoparticles to Retinal Ganglion Cells. Transl Vis Sci Technol 13, 5 (2024).

47. Welch, P. The use of fast Fourier transform for the estimation of power spectra: A method based on time averaging over short, modified periodograms. IEEE Trans. Audio Electroacoust. 15, 70–73 (1967).

48. Holm, S. A Simple Sequentially Rejective Multiple Test Procedure. Scand. J. Stat. 6, 65–70 (1979).

