## Supplementary Information for "Suprathreshold suppression underlies spatial selectivity in temporal interference stimulation"

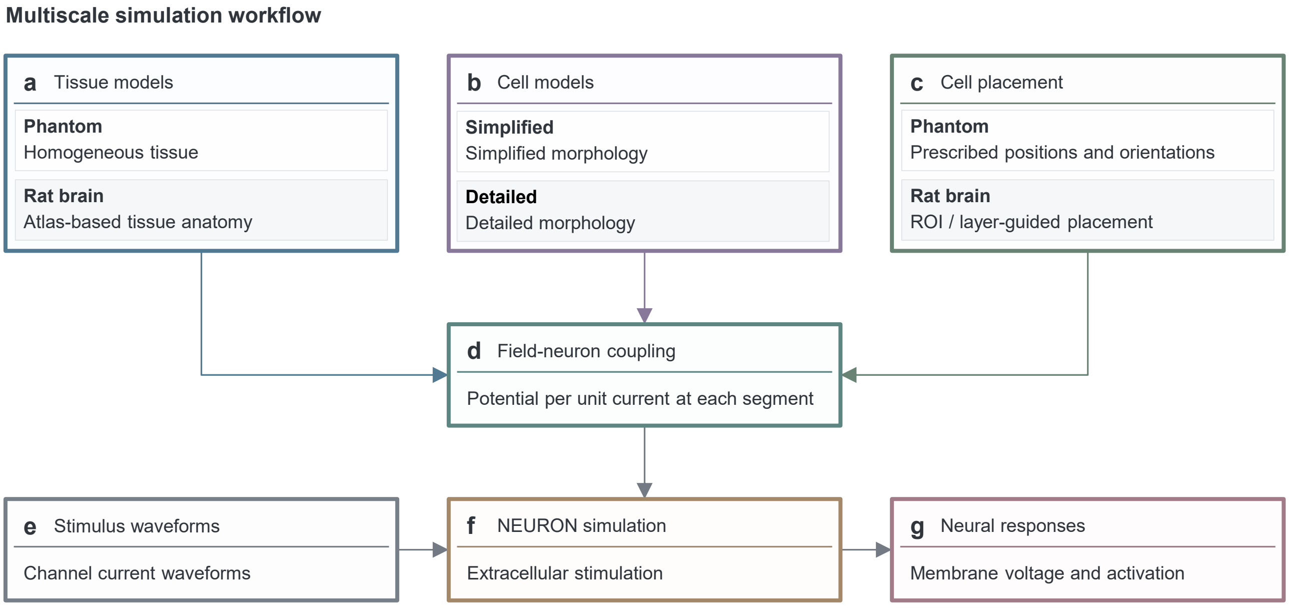


**Supplementary Fig. 1 | Multiscale simulation workflow for extracellular electrical stimulation.**
Boxes indicate modelling stages and arrows indicate data flow. **a**, Tissue volume-conductor models: a homogeneous phantom for Figs. 1–4 and an atlas-based rat brain for Fig. 5. Extracellular potentials are computed separately for each stimulation channel. **b**, Simplified neuronal morphologies used in Figs. 1–4 and detailed morphologies used in Fig. 5. **c**, Prescribed neuronal positions and orientations in the phantom, and ROI- and layer-guided placement in the rat brain. **d**, Evaluation of each channel's extracellular potential per unit current at every neuronal segment. **e**, Channel current waveforms for electrical stimulation. **f**, Time-dependent extracellular potentials are applied to the neuronal models in NEURON. **g**, Membrane potentials and ion-channel state variables are recorded and analysed to characterize neuronal responses.

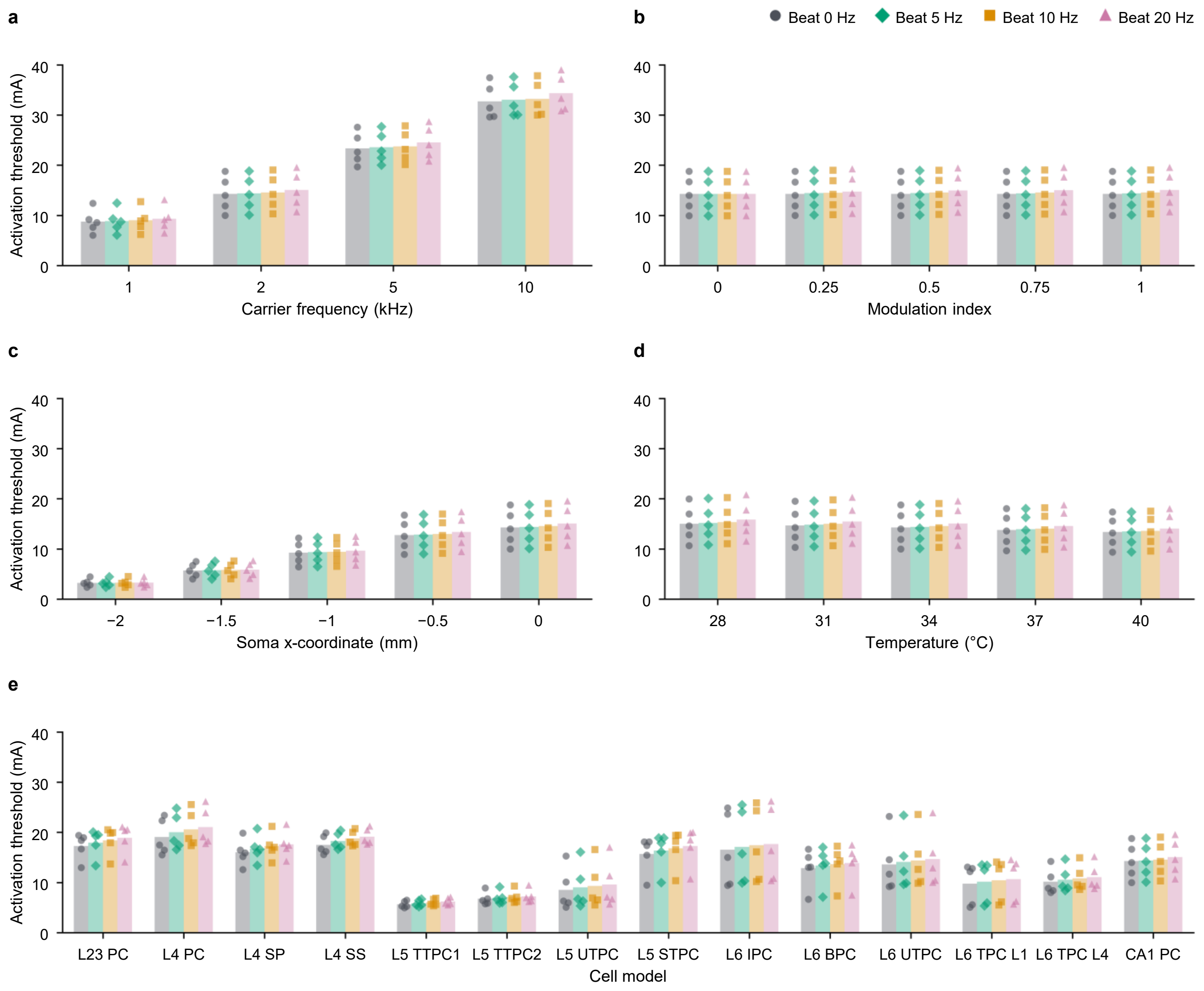


**Supplementary Fig. 2 | Absolute activation thresholds for HFS and TIS.**
Absolute activation thresholds, expressed as total current, for HFS and for TIS at beat frequencies of 5, 10 and 20 Hz, corresponding to Fig. 1c(i–v). **a**, Carrier frequency. **b**, Somatic MI; channel-current ratios of 1:0, 7:1, 3:1, 5:3 and 1:1 give MI values of 0, 0.25, 0.50, 0.75 and 1.00, respectively. At MI = 0, only CH1 was active. **c**, Soma x-coordinate. **d**, Temperature. **e**, Thirteen excitatory cortical model types and the CA1 PC model (Supplementary Table 1). Bars, means across five clones; symbols, individual clones. HFS is shown as grey circles and bars. Unless varied, simulations used the morphologically simplified, unmyelinated CA1 model at the phantom origin, 34 °C, a 2 kHz carrier and equal channel amplitudes.

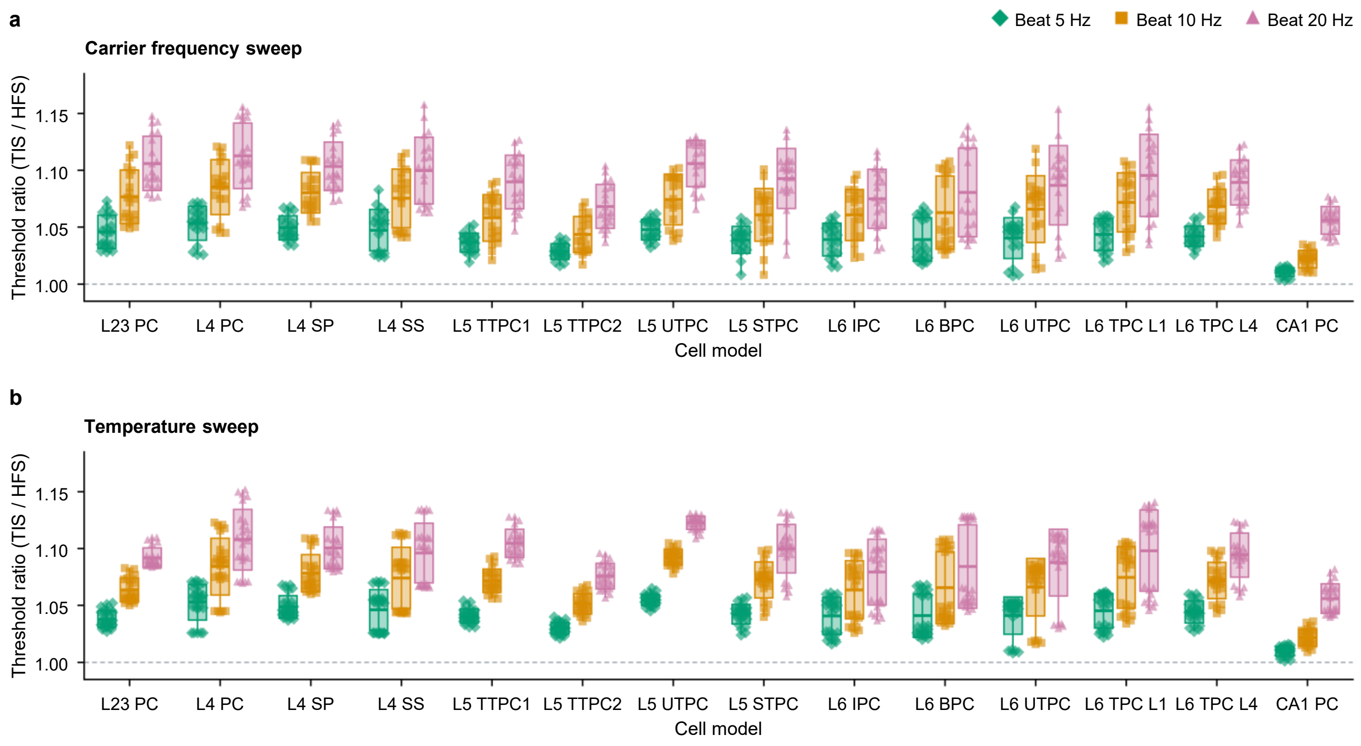


**Supplementary Fig. 3 | Activation-threshold ratios across carrier frequencies and temperatures.**
TIS-to-HFS activation-threshold ratios for the 13 excitatory cortical model types and CA1 PC model, each represented by five clones. TIS beat frequencies were 5, 10 and 20 Hz. **a**, Ratios pooled across carrier frequencies of 1, 2, 5 and 10 kHz at 34 °C: 20 clone–frequency combinations per model type and beat frequency. **b**, Ratios pooled across temperatures of 28, 31, 34, 37 and 40 °C at a carrier frequency of 2 kHz: 25 clone–temperature combinations per model type and beat frequency. Symbols, individual ratios from each clone and condition. Boxes span mean ± s.d. across the pooled combinations, horizontal lines mark means and thin vertical lines span the full range. Dashed line, ratio 1. Unless varied, simulations used the morphologically simplified, unmyelinated models at the phantom origin, 34 °C, a 2 kHz carrier and equal channel amplitudes.

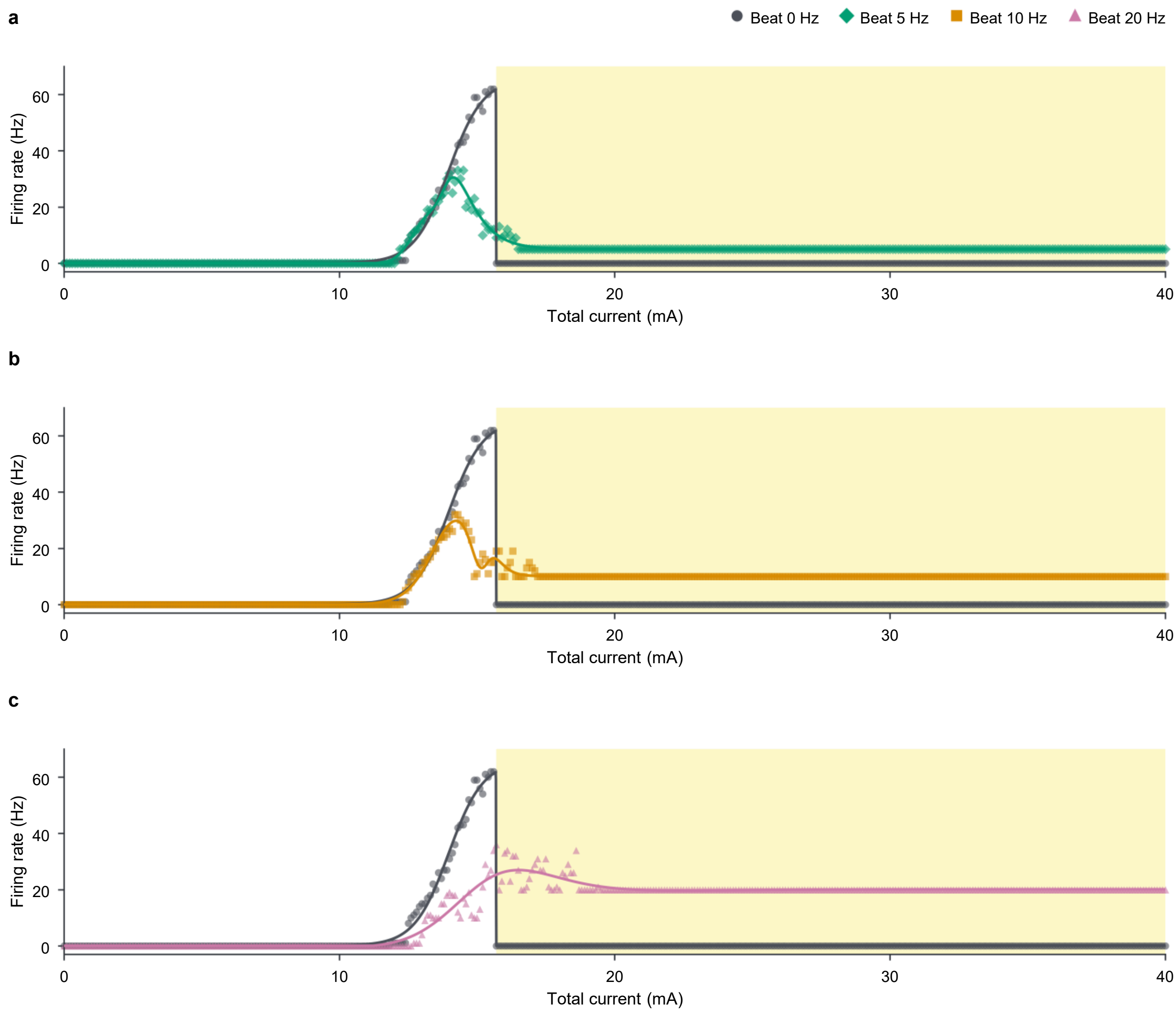


**Supplementary Fig. 4 | Beat-frequency dependence of suprathreshold somatic firing.**
Firing rate versus total current for TIS at beat frequencies of **a**, 5 Hz; **b**, 10 Hz; and **c**, 20 Hz, compared with the same HFS response. Symbols, simulation results; lines, fitted guides. Yellow shading, TOW. All simulations used the morphologically simplified, unmyelinated CA1 model at the phantom origin, 34 °C, a 2 kHz carrier and equal channel amplitudes.

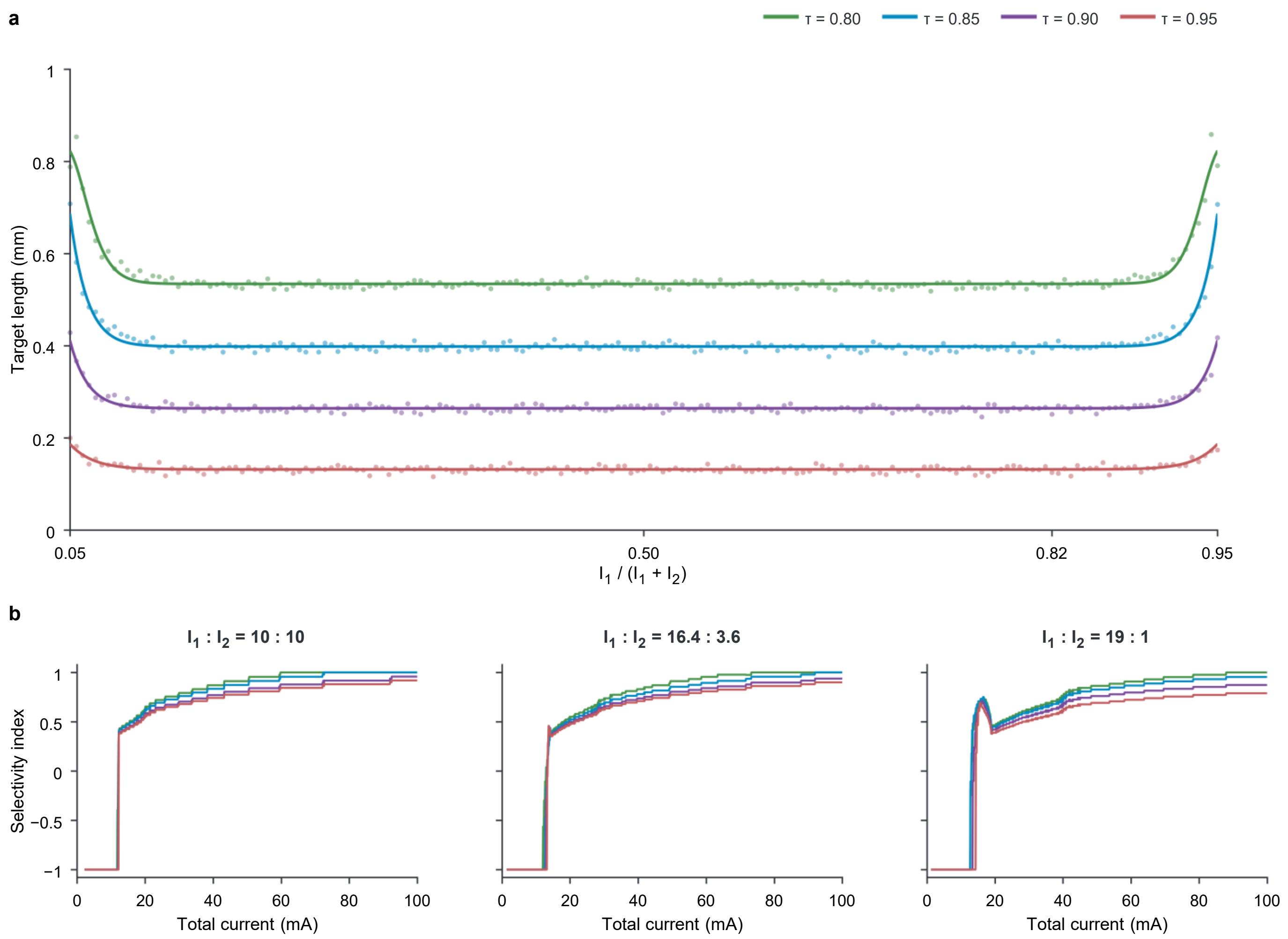


**Supplementary Fig. 5 | Sensitivity of target length and spatial selectivity to the modulation-index cutoff for target definition.**

The field maps and neuronal responses from Fig. 3 were reanalysed by defining target regions as locations with MI ≥ τ, with τ = 0.80, 0.85, 0.90 and 0.95. The main analysis used τ = 0.90. **a**, Summed target length along y = z = 0 versus the CH1 current fraction, $I_1/(I_1 + I_2)$, where $I_1$ and $I_2$ are the peak current amplitudes in CH1 and CH2, respectively. Symbols, calculated lengths; lines, fitted guides. At highly unequal current ratios, targets for τ = 0.80 and 0.85 approach the phantom boundary. **b, SI** versus total current. Columns show CH1:CH2 current ratios of 10:10, 16.4:3.6 and 19:1. SI was recalculated after reclassifying the same sampled positions as target or off-target at each τ.

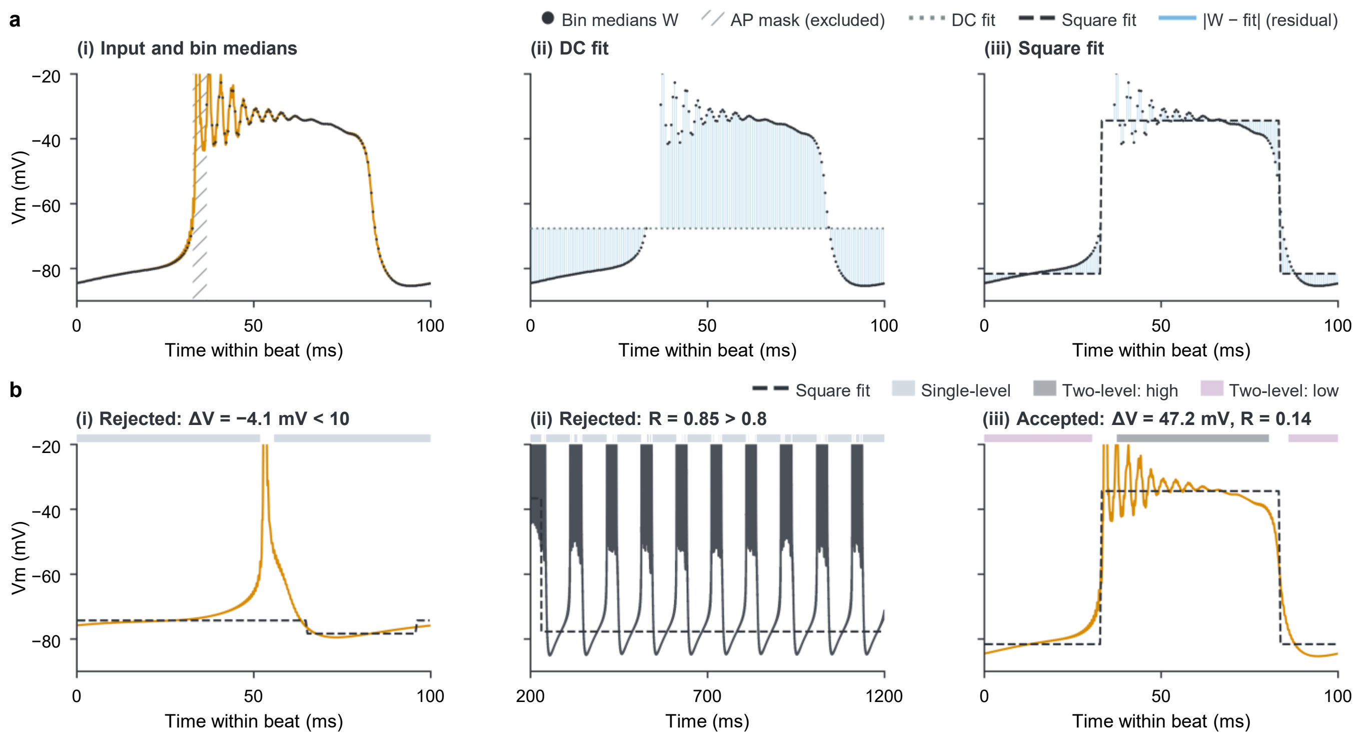


**Supplementary Fig. 6 | Selection of membrane-potential averaging windows.**
Window selection used AP-masked, filtered Vm. Each segment—one TIS beat or the HFS plateau—was fitted by a constant (DC) and a two-level function (Square), minimizing the sum of absolute differences between bin medians and fitted values. A two-level fit was accepted when its level difference ΔV was at least 10 mV and its Square-to-DC cost ratio R was at most 0.8; otherwise, the segment was classified as single-level. **a**, TIS at 20 mA, 600–700 ms: (i) filtered Vm and medians within 0.5 ms bins (W); hatched region, AP mask excluded from fitting and averaging; (ii) DC fit; (iii) Square fit. Light-blue lines, absolute residuals between W and the fitted values. The Square fit was accepted (ΔV = 47.2 mV, R = 0.14). **b**, Filtered Vm, candidate Square fits and the resulting averaging windows. (i) TIS at 13 mA, 900–1,000 ms: rejected because ΔV = −4.1 mV. (ii) HFS at 15 mA: rejected because R = 0.85. (iii) Accepted example from a. Bands above the traces, single-level, high and low averaging windows; gaps, excluded AP masks and transition intervals. Raw Vm was averaged within these windows. Time is relative to beat onset in the TIS panels and to simulation onset in the HFS panel. Vm is clipped above −20 mV for display only. Window categories describe the analysis and do not by themselves identify physiological states. All simulations used the morphologically simplified, unmyelinated CA1 model at the phantom origin, 34 °C, a 2 kHz carrier and equal channel amplitudes, with a 10 Hz beat for TIS.

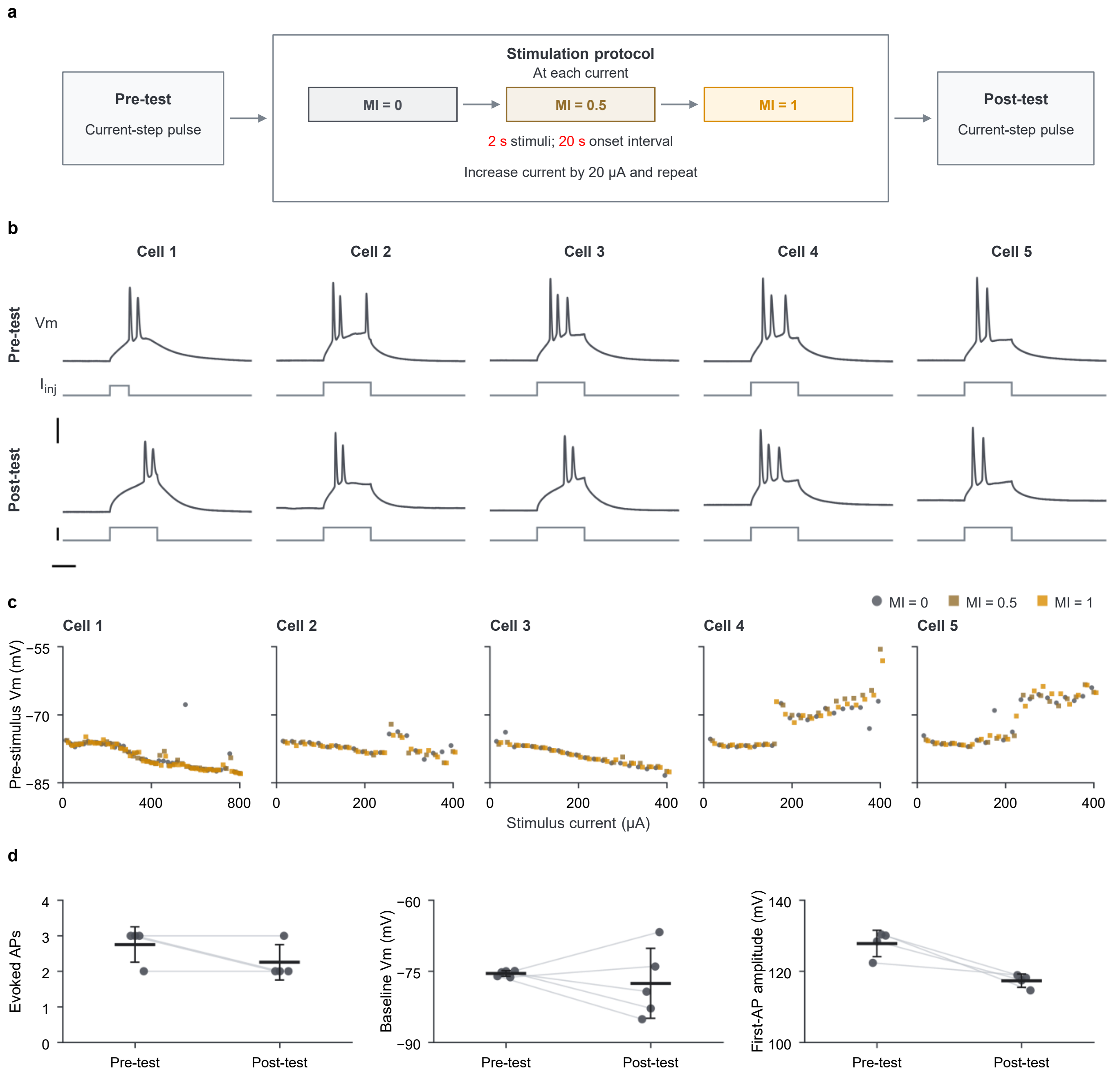


**Supplementary Fig. 7 | Current-step responses and baseline membrane potential across the extracellular stimulation protocol.**

Whole-cell recordings from the five neurons in the Fig. 6 dataset (n = 5 cells, one per mouse). **a**, Schematic of intracellular current-step tests before and after the extracellular stimulation series. **b**, Raw Vm and current-step commands during the pre-test and post-test; columns, Cells 1–5. Steps were 200 pA for 50 ms, except during the Cell 1 pre-test (150 pA for 20 ms). Scale bars, 25 ms, 40 mV and 200 pA. **c**, Mean raw Vm during the 50 ms preceding extracellular stimulation in each sweep, plotted against stimulus current. Symbols, individual sweeps, horizontally offset for visibility. **d**, Paired pre-test and post-test measurements, from left to right: AP count during the 60 ms after current-step onset, mean baseline Vm during the preceding 50 ms, and first-AP amplitude relative to the pre-step baseline. Symbols and grey connecting lines, individual cells; black horizontal lines and error bars, mean ± s.d. AP count and amplitude summaries include four cells, excluding Cell 1 because its pre-test step differed; baseline Vm includes all five cells.

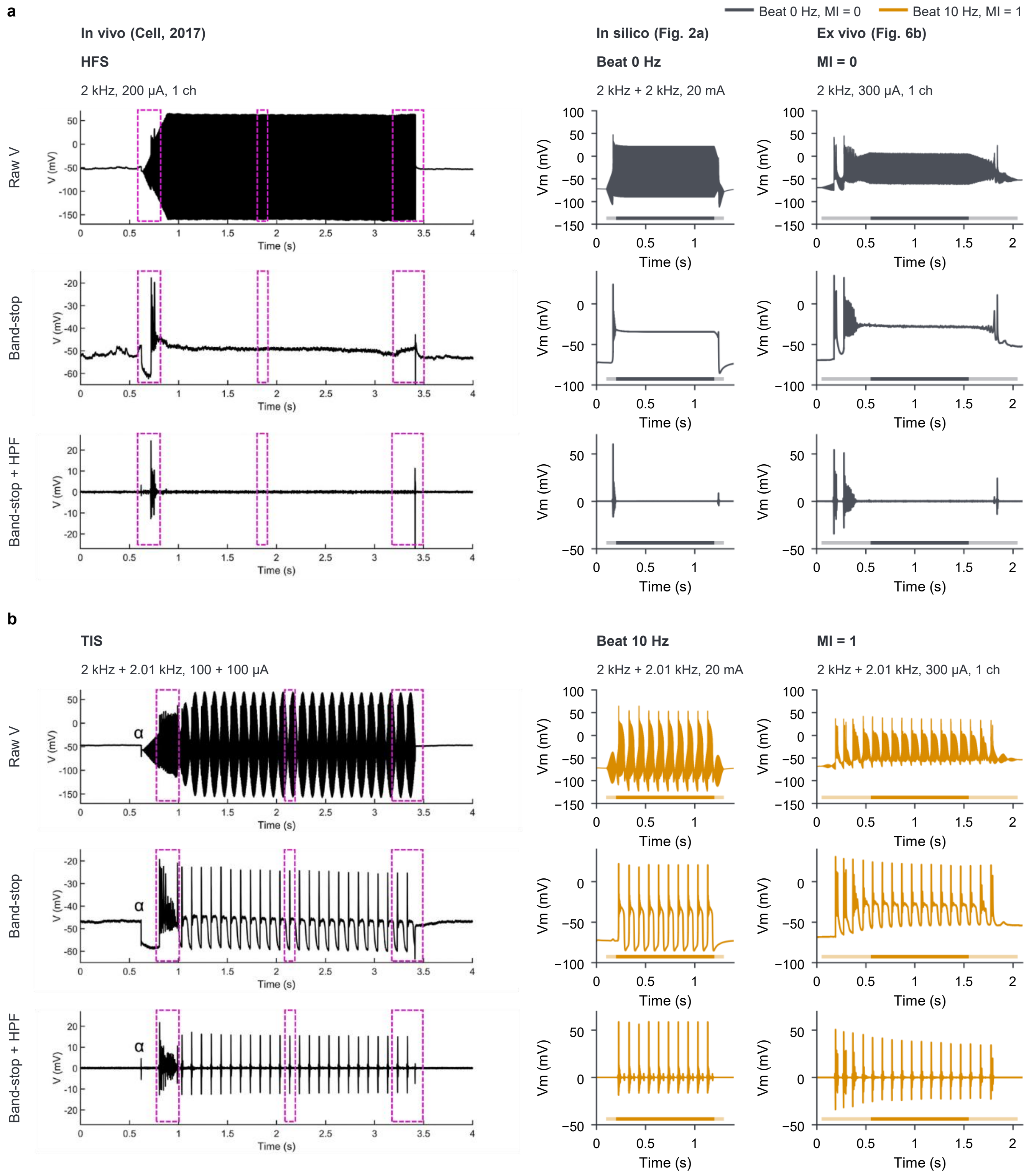


**Supplementary Fig. 8 | Simulated and ex vivo membrane-potential responses resemble published in vivo recordings.** Simulated and ex vivo membrane-potential traces were processed using the band-stop and high-pass filter settings reported by Grossman et al. (2017) for qualitative comparison with the published in vivo recordings. **a**, Unmodulated 2 kHz stimulation. **b**, TIS with components at 2,000 and 2,010 Hz. Columns, from left: published in vivo whole-cell recordings from anaesthetized mouse cortex (Grossman et al., 2017, Fig. S1D(i)–F(i) in a and S1A(i)–C(i) in b); CA1 model simulations under the conditions of Fig. 2a at 20 mA total current; and ex vivo whole-cell recordings from mouse S1-bf layer 4 (Fig. 6b, Cell 4) at 300 μA peak current. Rows, from top: raw Vm; Vm after 1–15 kHz band-stop filtering; and Vm after subsequent 100 Hz high-pass filtering. Zero in the high-pass-filtered traces does not represent resting Vm. Published traces were reproduced as images and were not reanalysed. All columns share a horizontal time scale. Voltage scales match between the model and ex vivo columns within each row; published panels retain their own scales. Coloured horizontal bars in the model and ex vivo columns, stimulation periods; lighter segments, ramps. Magenta boxes, original zoom regions; α, ground-connection artefact in the published recordings. Published stimulation used one channel at 200 μA in a and two channels at 100 μA each in b. The model used two equal-amplitude channels; both ex vivo waveforms were delivered through the same electrode pair using one stimulation channel. Current amplitudes do not imply equivalent local fields. Published panels adapted from Grossman et al., *Cell* 169, 1029–1041.e16 (2017), under CC BY 4.0: cropped, original panel labels and zoom numerals removed, uniformly scaled and rearranged.

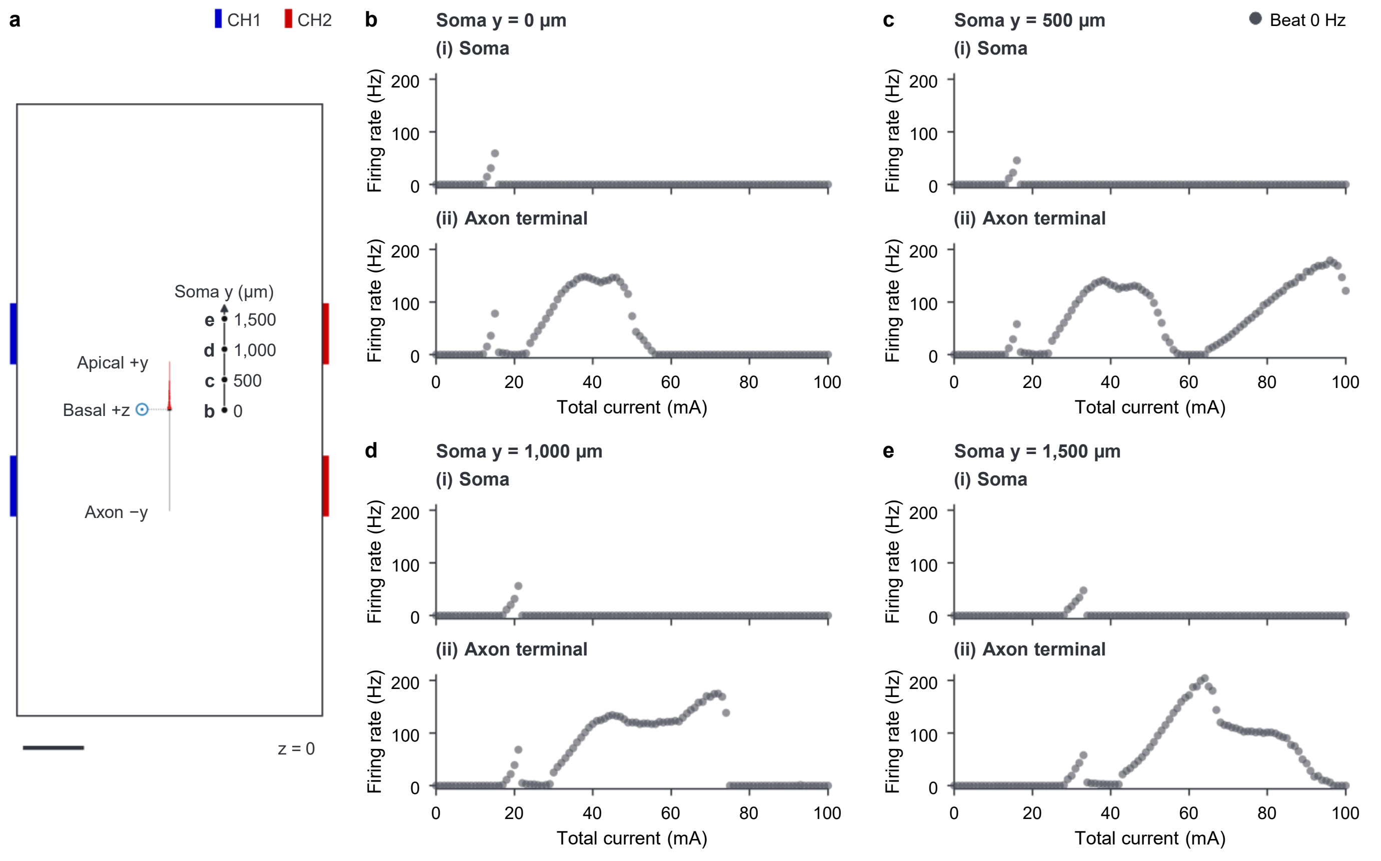


**Supplementary Fig. 9 | Position-dependent somatic and axonal firing under high-frequency stimulation.**
**a**, Placement schematic in the z = 0 plane. Scale bar, 1 mm. **b–e**, Plateau firing rate versus total current at soma positions y = 0, 500, 1,000 and 1,500 μm, respectively. Each panel shows responses at (i) the soma and (ii) the central segment of the terminal axonal section, at a path distance of 1,576 μm from the soma. Symbols, simulation results. The position at y = 1,000 μm in **d** was selected for the detailed analysis in Extended Data Fig. 10. All simulations used the model and HFS settings of Fig. 2a. The whole model was translated along the y axis of the phantom, with the soma at x = z = 0 and neuronal morphology and orientation held fixed.

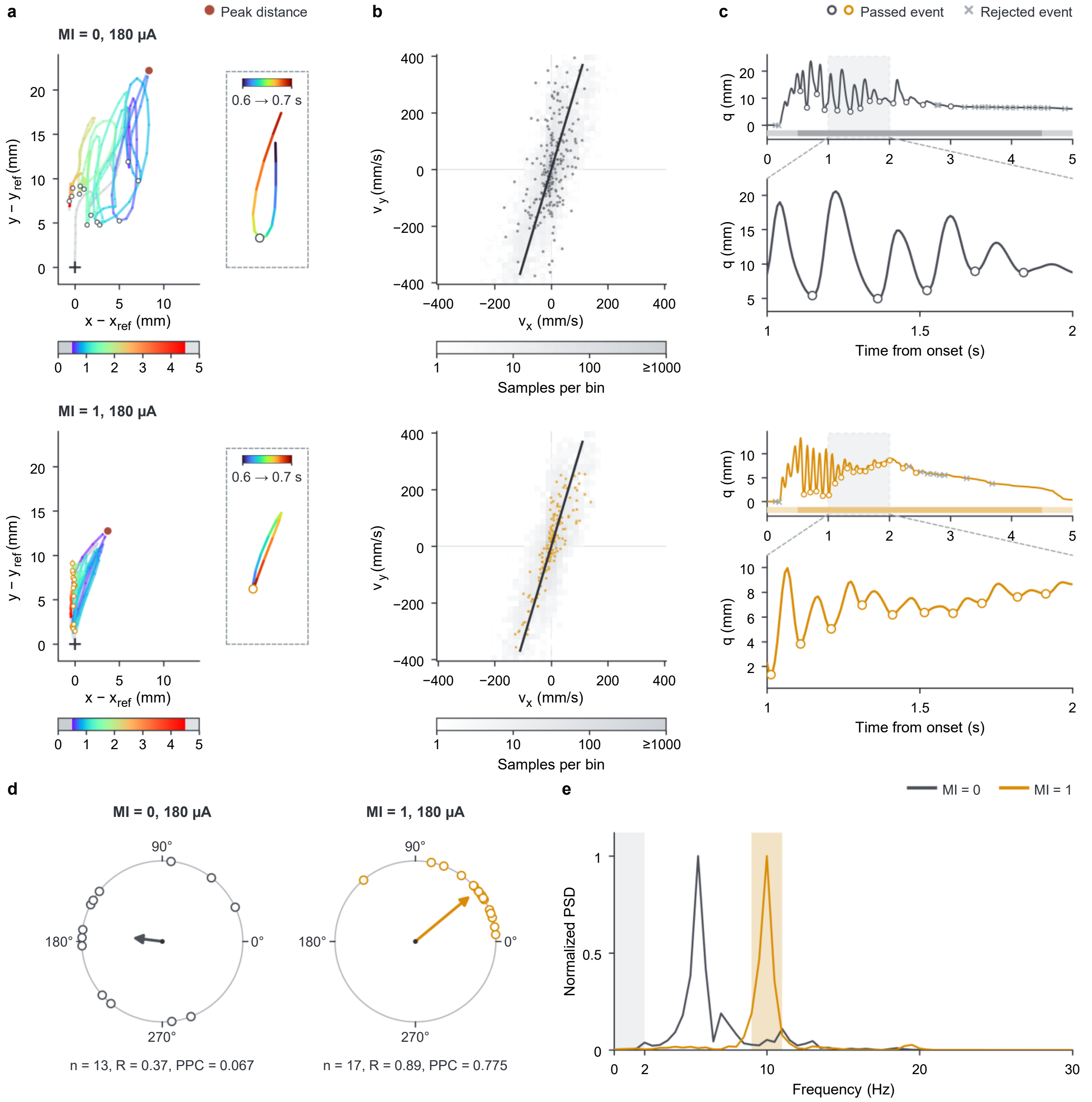


**Supplementary Fig. 10 | Detection of forelimb movement events and quantification of beat-frequency locking.** Illustration of the movement analysis using HFS and TIS epochs from Rat 5 at 180 µA. HFS is shown above TIS in **a–c** and to its left in **d**. **a**, Digit-2 trajectories during the 5 s stimulus relative to the pre-onset position. Grey segments, ramps; coloured segments, plateau time with logarithmic colour mapping. Cross, pre-onset position; filled dot, peak displacement; open circles, detected movement events. Right, trajectories over 0.6–0.7 s, shown at a shared spatial scale with separate time colour bars. **b**, Two-dimensional histograms of velocity samples pooled across both waveforms and all tested currents in this animal. Line, first principal component (PC1) defining the movement axis; overlaid points, velocity samples from the illustrated epoch. **c**, Position projected onto the movement axis over 0–5 s, with an enlarged 1–2 s interval. Open circles, accepted plateau minima; crosses, minima rejected by the prominence criterion. Horizontal bars, stimulation periods; lighter segments, ramps. **d**, Movement-event phases referenced to a 10 Hz clock starting at stimulation onset for both waveforms. Arrow, mean phase vector with length *R*; *n*, number of plateau events. **e**, Power spectral density (PSD) of movement-axis velocity during the plateau, normalized within each epoch to its maximum over 2–30 Hz. Orange shading, 9–11 Hz; grey shading, frequencies below 2 Hz. Stimulation was as in Fig. 8. Event and spectral analyses used the 0.5–4.5 s plateau. Processing parameters are provided in Methods and Supplementary Table 5.

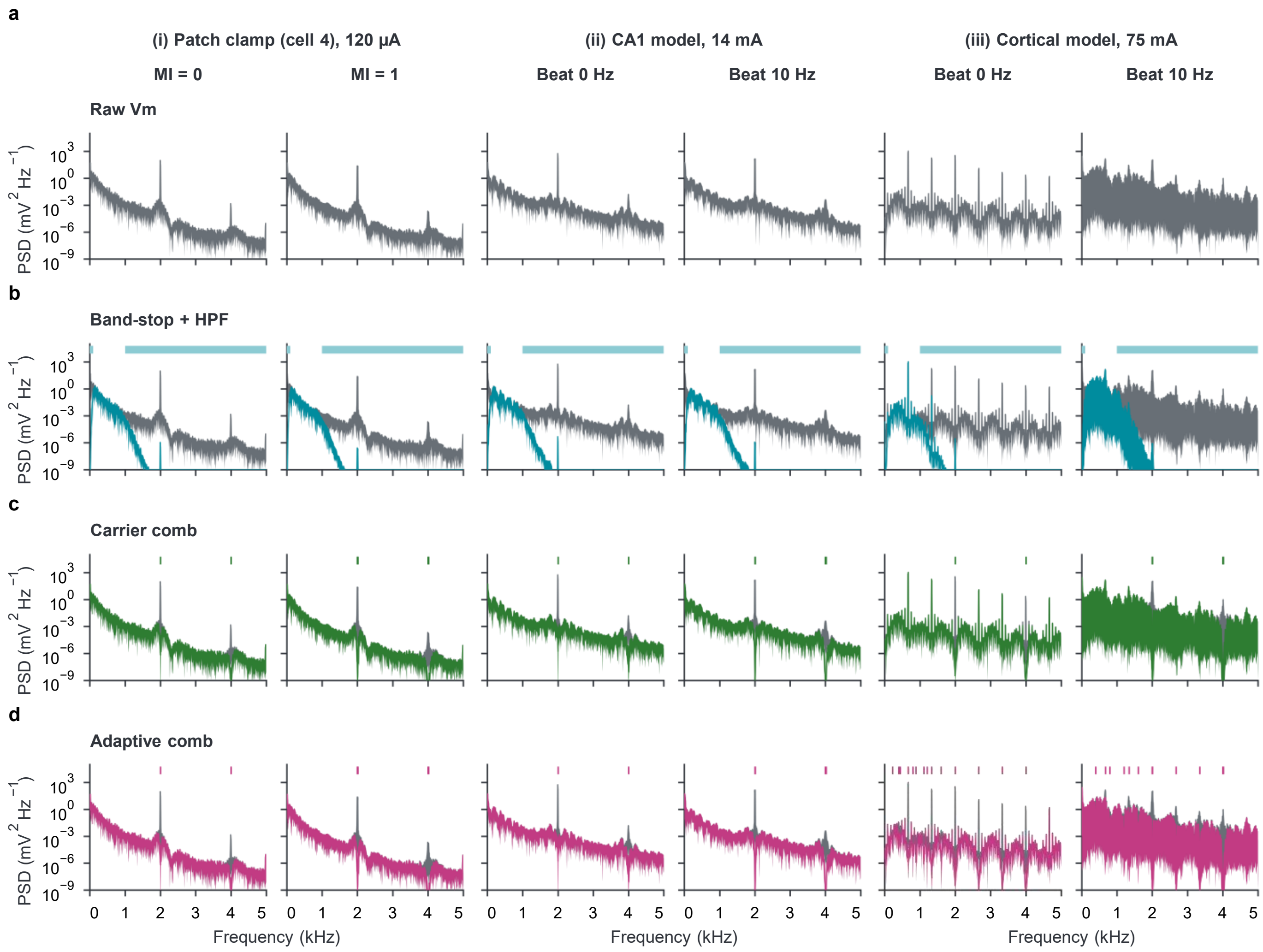


**Supplementary Fig. 11 | Comparison of membrane-potential spectra across filtering procedures.**
Power spectral density (PSD) of raw and filtered somatic Vm during the 1 s stimulation plateau. Column groups: (i) Cell 4 from Fig. 6, an S1-bf layer 4 neuron, at 120 µA; (ii) CA1 PC clone 1 at 14 mA; and (iii) L4 PC clone 1 at 75 mA. Within each group, HFS is shown on the left and TIS on the right. **a**, Raw Vm spectra (grey), also shown in the background in **b–d**. **b**, Spectra after 1–15 kHz band-stop and 100 Hz high-pass filtering (teal; adapted from Grossman et al., 2017). Horizontal bars, stop bands. **c,d**, Spectra after carrier-comb (**c**, green) and adaptive-comb (**d**, raspberry) filtering. Vertical ticks, notch frequencies within the displayed range. Additional subharmonic targets were selected only for the cortical example; carrier-comb and adaptive-comb spectra coincide in **i** and **ii**. Simulation placement and stimulation settings were as in Fig. 2a; whole-cell recording and stimulation settings were as in Fig. 6.

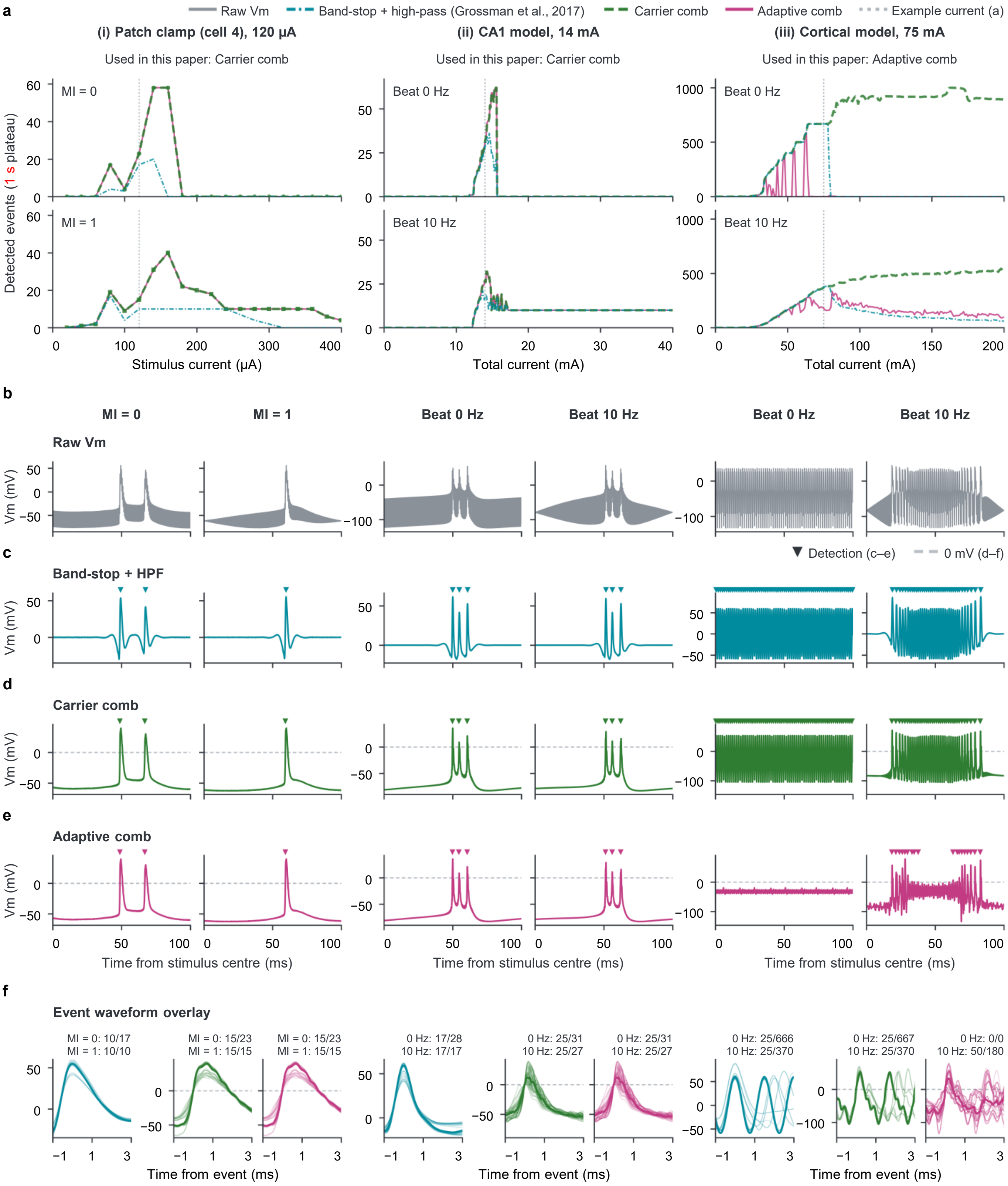


**Supplementary Fig. 12 | Dependence of detected events on filtering and detection procedures.**
The cell and neuronal models, column order and common recording and stimulation protocols are as in Supplementary Fig. 11. Procedures comprise band-stop plus high-pass filtering followed by peak detection (teal dash-dot; adapted from Grossman et al., 2017), carrier-comb filtering (green dashed) and adaptive-comb filtering (raspberry). Both comb procedures use upward 0 mV crossings; the band-stop/high-pass procedure uses peaks at least 40 mV above a 20 ms running mean and omits the original waveform-shape criterion. **a**, Detected events per 1 s plateau versus current, with HFS above TIS. One sweep or simulation was analysed per waveform and current. Lines, connections between adjacent tested currents; vertical dotted lines, the examples in **b–f** at 120 µA, 14 mA and 75 mA in **i–iii**, respectively. **b–e**, Raw Vm (**b**) and outputs after band-stop plus high-pass (**c**), carrier-comb (**d**) and adaptive-comb (**e**) filtering over the 100 ms interval starting at the stimulus midpoint. Within each pair, HFS is shown on the left and TIS on the right. Triangles, detected events; dashed lines, 0 mV in **d–f**. **f**, Filtered waveforms aligned to each procedure's detected events, from −1 to +3 ms; procedures are arranged in the order shown in **c–e**. Equal numbers of waveforms were sampled from each stimulation condition, up to 25 per condition, or up to 50 from one condition when the other had none. Thin traces, individual waveforms; thick traces, medians of the displayed waveforms pooled across stimulation conditions; annotations, displayed/total detected events per condition. Currents denote experimental peak amplitude and simulated total current.

### Supplementary Table 1

#### Neuronal model types and abbreviations.

Each of the 13 cortical cell types and the CA1 pyramidal-cell type was represented by five clones, giving 70 model instances. Cortical models were obtained from the Blue Brain Neocortical Microcircuit Collaboration Portal (Markram et al., 2015); CA1 models were obtained from the Hippocampus Hub (Migliore et al., 2018; Romani et al., 2024). All axons were unmyelinated. Detailed and morphologically simplified versions used the same source membrane-mechanism assignments in retained sections.

##### A. Model labels

| Figure label | Cell type | Figure label | Cell type |
|---|---|---|---|
| L23 PC | Layer 2/3 pyramidal cell | L5 STPC | Layer 5 slender-tufted pyramidal cell |
| L4 PC | Layer 4 pyramidal cell | L6 IPC | Layer 6 pyramidal cell with inverted apical-like dendrites |
| L4 SP | Layer 4 star pyramidal cell | L6 BPC | Layer 6 pyramidal cell with bipolar apical-like dendrites |
| L4 SS | Layer 4 spiny stellate cell | L6 UTPC | Layer 6 untufted pyramidal cell |
| L5 TTPC1 | Layer 5 thick-tufted pyramidal cell; late apical bifurcation | L6 TPC L1 | Layer 6 tufted pyramidal cell with apical dendrites reaching layer 1 |
| L5 TTPC2 | Layer 5 thick-tufted pyramidal cell; early apical bifurcation | L6 TPC L4 | Layer 6 tufted pyramidal cell with apical dendrites reaching layer 4 |
| L5 UTPC | Layer 5 untufted pyramidal cell | CA1 PC | CA1 pyramidal cell |

### Supplementary Table 2

#### Volume-conductor parameters and neuronal placement.

Conductivities were assigned isotropically. The homogeneous phantom measured 5 × 10 × 10 mm; rat-brain geometry used the Waxholm Space Sprague Dawley atlas v4.01 (Kleven et al., 2023). Conductivity sources are cited in Methods.

##### A. Tissue conductivities

| Tissue class | Conductivity (S $m^{-1}$) | Anatomical basis or assigned value |
|---|---|---|
| White matter | 0.126 | WHS white-matter labels |
| Grey matter; phantom tissue | 0.275 | WHS grey-matter labels and homogeneous phantom tissue |
| Ventricular system | 1.654 | CSF value for ventricular labels |
| Inner-ear organs | 1.654 | CSF value for the cochlea and vestibular apparatus |
| Inner-ear nerves and spiral ganglion | 0.126 | White-matter value for the cochlear nerve, vestibular nerve and spiral ganglion |
| Spinal cord | 0.126 | White-matter value for the entire spinal-cord label (SpC, 45), based on visual inspection of peripheral white matter in the represented segment |

Tissue grouping followed the anatomical hierarchy in the WHS v4 atlas supplement (Kleven et al., 2023) and the v4.01 label definitions.

##### B. Electrode montage

| Conductor | Electrode | Diameter (mm) |
|---|---|---|
| Phantom | Reference | 1 |
| Phantom | CH1 + | 1 |
| Phantom | CH1 − | 1 |
| Phantom | CH2 + | 1 |
| Phantom | CH2 − | 1 |
| Rat brain | Reference | 2 |
| Rat brain | CH1 + | 1 |
| Rat brain | CH1 − | 2 |
| Rat brain | CH2 + | 1 |
| Rat brain | CH2 − | 2 |

Each channel used a pair of active pads and a common reference pad with zero net applied current. In the rat-brain model the pads were applied to the brain surface, with scalp and skull omitted.

#### Supplementary Table 2 (continued)

##### C. Placement and population weighting

| Region / analysis | Soma positions | Orientations per position | Placement | Layer volume ($mm^3$) |
|---|---|---|---|---|
| Phantom steering | 97 | 1 | Positions along the x-axis | — |
| S1-bf layer 4 | 100 per clone | 3 | Normalized depth 0.4 | 1.57803 |
| CA1 pyramidal layer | 1,000 per clone | 1 | Normalized depth 0.7 | 1.47635 |

Positions were shared across clones within each region. S1-bf depth runs from the pial boundary (0) to the white-matter boundary (1); CA1 depth runs from the dentate-facing boundary (0) to the opposite boundary (1). Reference layer thicknesses informed the selected depths in S1-bf (Markram et al., 2015) and CA1 (Romani et al., 2024). Fig. 5 pooled nine L4 clones (300 configurations each) and two CA1 clones (1,000 configurations each). Each configuration was weighted by its regional layer volume divided by the number of configurations in that regional population; the weighting does not encode measured cell-type abundance.

### Supplementary Table 3

#### Morphology boundary checks across candidate placement depths.

Counts of placements with at least one morphology point outside the permitted atlas mask. The selected depths were 0.4 for S1-bf layer 4 and 0.7 for CA1. A clone was included in Fig. 5 only when all tested placements passed at the selected depth.

##### A. S1-bf layer 4

| Model | Depth 0.35 | Depth 0.4 (selected) | Included in Fig. 5 |
|---|---|---|---|
| L4 PC clone 1 | 300 | 300 | No |
| L4 PC clone 2 | 0 | 0 | Yes |
| L4 PC clone 3 | 0 | 0 | Yes |
| L4 PC clone 4 | 0 | 0 | Yes |
| L4 PC clone 5 | 0 | 0 | Yes |
| L4 SP clone 1 | 0 | 0 | Yes |
| L4 SP clone 2 | 248 | 54 | No |
| L4 SP clone 3 | 81 | 31 | No |
| L4 SP clone 4 | 80 | 2 | No |
| L4 SP clone 5 | 298 | 155 | No |
| L4 SS clone 1 | 0 | 0 | Yes |
| L4 SS clone 2 | 26 | 60 | No |
| L4 SS clone 3 | 6 | 0 | Yes |
| L4 SS clone 4 | 0 | 0 | Yes |
| L4 SS clone 5 | 0 | 0 | Yes |

Failure counts are out of 300 configurations per clone at each depth (100 soma positions × three axial orientations). The containment mask includes S1-bf and neighbouring cortical grey matter.

### B. CA1

| Model | Depth 0.7 (selected) | Included in Fig. 5 |
|---|---|---|
| CA1 PC clone 1 | 7 | No |
| CA1 PC clone 2 | 2 | No |
| CA1 PC clone 3 | 2 | No |
| CA1 PC clone 4 | 0 | Yes |
| CA1 PC clone 5 | 0 | Yes |

Failure counts are out of 1,000 soma positions per clone.

##### C. Anatomical masks for morphology boundary checks

| Region | Permitted anatomical regions |
|---|---|
| S1-bf | S1-bf and neighbouring cortical grey matter. |
| CA1 | CA1 and selected neighbouring structures, including the entire ipsilateral unspecified ventricular label (V-u). |

Masks were defined in the left hemisphere. A placement failed if any sampled morphology point lay outside its permitted mask.

### Supplementary Table 4

#### Experimental current series and data inclusion.

Experimental sampling and inclusion for the brain-slice, retinal and in vivo datasets. Each brain-slice neuron and retina was obtained from a different mouse. Experimental protocols and pharmacological conditions are described in Methods.

##### A. Brain-slice current series and inclusion

| Cell | Current range (μA) | Valid / planned sweeps | Matched TOW currents (μA) | Excluded conditions |
|---|---|---|---|---|
| 1 | 20–800 | 116 / 120 | 660, 680, 700, 760, 780, 800 | MI = 0: 140, 540 μA; MI = 0.5: 720 μA; MI = 1: 740 μA |
| 2 | 20–400 | 60 / 60 | 200–400 | None |
| 3 | 20–400 | 60 / 60 | 140–400 | None |
| 4 | 20–400 | 60 / 60 | 180–360 | None |
| 5 | 20–400 | 60 / 60 | 280–400 | None |

Currents were tested in 20 μA steps, with one sweep per waveform and current. Matched TOW currents are those used in Fig. 6d, requiring valid recordings for MI = 0, 0.5 and 1. Listed ranges include every 20 μA step. Excluded conditions lacked a complete plateau recording; missing responses were not assigned zero firing.

##### B. Retinal steering configurations

| Retina | CH1:CH2 ratio A | CH1:CH2 ratio B |
|---|---|---|
| 1 | 1:3 | 7:1 |
| 2 | 1:7 | 3:1 |
| 3 | 1:1 | 1:7 |

Each retina was tested at both listed CH1:CH2 current ratios. Fig. 7f compares the two recording regions separately within each retina and ratio at 1.5 mA total current.

##### C. In vivo current series and inclusion

| Rat | Current range; step (μA) | HFS / TIS epochs | Matched PPC currents (n) | Total injected volume (nL) |
|---|---|---|---|---|
| 1 | 40–400; 40 | 10 / 10 | 9 | 700 |
| 2 | 20–400; 20 | 19 / 20 | 12 | 600 |
| 3 | 20–400; 20 | 20 / 20 | 11 | 600 |
| 4 | 20–300; 20 | 15 / 15 | 12 | 500 |
| 5 | 20–300; 20 | 15 / 15 | 9 | 500 |

One train per waveform and current. Rat 2 received HFS at 360 μA, but no video was recorded. The movement dataset contains 159 epochs and 79 matched HFS–TIS current pairs. Fig. 8b used 53 matched pairs with at least four plateau events in both waveforms, summarized as five animal means. The spectral comparisons in Extended Data Fig. 9b used all 79 matched pairs. Injection volumes are cumulative totals by the analysed recording stage.

#### Supplementary Table 5

##### Movement tracking, analysis parameters and sensitivity.

Training and evaluation of the session-specific tracking models, movement-analysis settings, coordinate completeness and sensitivity to event prominence.

##### A. Tracking model training and evaluation

| Rat | Labelled frames | Training frames | Test frames | Digit-2 test error (pixels) |
|---|---|---|---|---|
| 1 | 64 | 54 | 10 | 1.38 |
| 2 | 60 | 51 | 9 | 1.82 |
| 3 | 62 | 52 | 10 | 1.79 |
| 4 | 62 | 52 | 10 | 1.26 |
| 5 | 60 | 51 | 9 | 1.42 |

##### B. Coordinate, event and spectral analysis parameters

| Parameter | Value or rule |
|---|---|
| Median filter / likelihood cutoff | 5 frames / 0.4 |
| Initial gap interpolation | Linear interpolation of internal gaps of up to 10 frames |
| Spatial calibration | Per-video median separation of two reference points 19.70 mm apart |
| Epoch inclusion | At least 50% valid coordinates after initial preprocessing; all 159 epochs passed |
| Additional temporal-analysis filling | Remaining internal gaps: linear interpolation; boundary gaps: nearest valid coordinate |
| Coordinate low-pass filter | Second-order Butterworth, 20 Hz, forward and backward |
| Velocity sampling frequency | Frame rate recorded in the video metadata |
| Movement axis | PC1 of pooled two-dimensional velocity samples from both waveforms and all currents within each animal |
| Movement event / prominence | Projected-position minimum at a negative-to-positive velocity crossing / at least 1 image pixel |
| Plateau window | 0.5–4.5 s after stimulus onset |
| PPC comparison inclusion | At least four plateau events in both waveforms at the same current |
| Phase clock / interval bins | 10 Hz from stimulus onset / 0.1 reference period |

### Supplementary Table 5 (continued)

#### B. Coordinate, event and spectral analysis parameters (continued)

| Parameter | Value or rule |
|---|---|
| Spectral display | Each epoch normalized to its maximum within 2–30 Hz. |
| Band-power fraction | Integral over 9–11 Hz divided by integral over 2–30 Hz, with interpolation at exact boundaries |
| Dominant frequency | Frequency of maximum native PSD within 2–30 Hz |

Event prominence corresponded to approximately 0.094–0.136 mm across videos. Interval distributions were normalized using all intervals, including those outside the displayed range.

#### C. Tracking completeness during stimulation

| Rat | Stimulation frames | Low-likelihood frames, n (%) | Initial interpolation, n (%) | Additional filling, n (%) |
|---|---|---|---|---|
| 1 | 12,040 | 42 (0.349) | 42 (0.349) | 0 (0.000) |
| 2 | 23,476 | 21 (0.089) | 21 (0.089) | 0 (0.000) |
| 3 | 24,079 | 67 (0.278) | 1 (0.004) | 66 (0.274) |
| 4 | 18,058 | 92 (0.509) | 21 (0.116) | 71 (0.393) |
| 5 | 18,049 | 38 (0.211) | 38 (0.211) | 0 (0.000) |
| Total | 95,702 | 260 (0.272) | 123 (0.129) | 137 (0.143) |

Counts cover the 159 LED-on stimulation epochs, including ramps. Percentages use the stimulation-frame total in each row. A total of 155 epochs were complete after initial preprocessing; the four remaining epochs are listed in D.

#### D. Epochs requiring additional coordinate filling

| Rat | Waveform | Current (μA) | Filled / total frames | Gap locations |
|---|---|---|---|---|
| 3 | TIS | 20 | 66 / 602 | 50 internal; 16 trailing |
| 4 | HFS | 200 | 34 / 602 | 18 + 16 internal |
| 4 | HFS | 240 | 16 / 602 | 16 leading |
| 4 | TIS | 220 | 21 / 602 | 21 leading |

All four epochs were retained for the primary temporal analysis. Path-length analyses omitted these epochs. Peak-displacement analyses excluded epochs without a valid pre-onset reference or with the peak at a filled frame.

### Supplementary Table 5 (continued)

#### E. Sensitivity to movement-event prominence

| Prominence (pixels) | Matched current pairs | Retained events | PPC difference (95% CI) | Raw p | Holm-adjusted p |
|---|---|---|---|---|---|
| 0 | 79 | 4,602 | 0.1644 (0.0168–0.3120) | 0.03647 | 0.05524 |
| 0.25 | 61 | 2,819 | 0.2548 (0.1278–0.3818) | 0.00509 | 0.01526 |
| 0.5 | 57 | 2,622 | 0.2495 (0.1472–0.3517) | 0.00248 | 0.00743 |
| 1 (primary) | 53 | 2,418 | 0.2336 (0.0801–0.3871) | 0.01342 | 0.04027 |
| 2 | 52 | 2,233 | 0.2667 (0.0698–0.4635) | 0.01975 | 0.05437 |
| 4 | 49 | 2,027 | 0.2823 (0.0664–0.4981) | 0.02214 | 0.05437 |

Differences are TIS minus HFS animal means with unadjusted 95% confidence intervals. Two-sided paired t-tests used five animals. Holm correction covered PPC and the two spectral outcomes separately for each prominence threshold. The matched-current set changes with the event-count criterion. TIS exceeded HFS in all five animals at every threshold; adjusted $p < 0.05$ was not retained at 0, 2 or 4 pixels.
